# Using genetic instruments to estimate interactions in Mendelian Randomization studies

**DOI:** 10.1101/544734

**Authors:** Teri-Louise North, Neil M Davies, Sean Harrison, Alice R Carter, Gibran Hemani, Eleanor Sanderson, Kate Tilling, Laura D Howe

**Author notes:** **Data and code availability:** The analytic scripts used to clean the data and produce the results are available on request from the authors. UK Biobank data are available on request from the study directly. Please contact for further information.

## Abstract

**BACKGROUND:** The interactive effect of two exposures on an outcome can be confounded. We demonstrate the use of Mendelian Randomization (MR) to estimate unconfounded additive interactions.

**METHODS:** Using simulation, we test an extension to multivariable MR using two-stage least squares to estimate the additive interaction between two continuous exposures on a continuous outcome, including scenarios where one exposure has a causal effect on the other (mediation). The interaction parameters were set to be one third of the main effect’s parameters to impose a limit on the variance explained by interaction terms. We compare the performance of the two-stage least squares estimator to a Factorial MR design, in which genetic risk scores for each exposure are dichotomised to create four groups, akin to a factorial randomized controlled trial. As an illustrative example, we apply factorial MR and the 2SLS estimator to the interactive effect of education and BMI on systolic blood pressure in UK Biobank.

**RESULTS:** Our simulations demonstrate that factorial MR has very low statistical power; 5-7% at N=50,000 and 8-23% at N=500,000 across the range of parameters tested. The two-stage least squares estimator had higher power to detect interactions than factorial MR and a lower type I error. For N=500,000, the two-stage least squares estimator had a power ranging from 29.7-92.9% and a type I error ranging from 4-6%. 95% Monte Carlo confidence intervals suggested that the estimator was unbiased to a reasonable degree of accuracy at this sample size. In comparison, the power at N=50,000 was 7-55%.

**CONCLUSIONS:** A two-stage least squares estimator using genetic risk scores for each exposure is a more powerful alternative for detecting an unconfounded additive interaction of two exposures on an outcome than existing approaches that rely on dichotomising genetic risk scores, but it requires a large sample size and instruments of adequate strength.

## INTRODUCTION

Epidemiology typically relies on observational data, which can suffer from confounding and reverse causality. Mendelian Randomization (MR) is an application of instrumental variable (IV) analysis, in which genetic variants robustly related to an exposure of interest are used to infer causal associations between the exposure and other phenotypic outcomes (1, 2). Children inherit their genomes from their parents at random. Such randomization means that reverse causality can be ruled out and genetic variants are often less related to environmental confounders (1). Multivariable MR (MVMR) is a method that can be used to estimate the effect of two or more exposures on an outcome (3). The approach is suitable in a range of scenarios, e.g. when two exposures are closely related, or when one exposure is hypothesised to mediate the effect of the other. MVMR requires a set of genetic variants that are associated with the exposure variables but do not affect the outcome other than through these exposure variables.

In many epidemiological research topics, it is of interest to explore whether the effect of one exposure depends on another exposure, i.e. to investigate an interaction (4). A recent study(5) used genetic IVs to apply a factorial design to examine whether lowering LDL-C using ezetimibe, statins or a combination of both is most effective for lowering Coronary Heart Disease (CHD) risk. This ‘2 x 2 factorial mendelian randomization study design’ mimics a factorial Randomized Controlled Trial (RCT) in which participants are randomly allocated to one of four treatment regimens (treatment 1, treatment 2, both treatments 1 and 2, or neither treatment). Genetic risk scores for each treatment exposure were generated and were split at the median, and the resultant four groups (reference, Ezetimibe, Statin, both) compared with respect to their association with CHD (5). While this approach can give an indication of whether interaction might be present, it does not quantify the interaction, and the dichotomisation of genetic risk scores means power is likely to be low. To date, MR has not been used to estimate interaction terms (with the exception of gene-environment interactions(6)) and such methods in an MR context have not yet been thoroughly investigated. External to the MR literature, instrumental variable approaches for interaction terms have been considered, and our proposed approach draws on the models used in these non-genetic IV analyses (7, 8).

A common approach to implement MR is two-stage least squares (2SLS), where the exposure is predicted from its instrument and regressed on the outcome with statistical adjustment of the standard errors in the second stage. In this study, we propose an extension to MVMR, using a 2SLS estimator to estimate additive interactions between two continuous exposures on a continuous outcome. We use simulations to assess the performance of the estimator, including scenarios where one exposure has a causal effect on the second exposure, i.e. there is mediation in addition to interaction. We compare the 2SLS estimator with the two-by-two factorial MR design. As an illustrative example, we apply the proposed 2SLS method to the interactive effect of years of schooling and BMI on systolic blood pressure in UK Biobank (9).

## METHODS

We explore an extension to MVMR that enables the estimation of an additive interaction between two exposures, by using the genetic risk scores for each exposure (Z_1_ and Z_2_) and the product of the two genetic risk scores, i.e. Z=(Z_1_,Z_2_,Z_1_Z_2_) as the instrumental variables (Z). When the first exposure has a causal effect on the second exposure, i.e. exposure 2 mediates the effect of exposure 1 on the outcome, the instrument is Z=(Z_1_,Z_2_,Z_1_Z_2_,Z_1_Z_1_). We use simulations to evaluate the performance of the 2SLS MVMR estimator for assessing additive interactions, with and without a causal effect of exposure 1 on exposure 2 (mediation), and compare these estimators with a factorial MR design. For both approaches, we consider power and type I error, and for the 2SLS estimators we also evaluate bias and coverage of the interaction coefficient.

### Simulation study: data generating process

Figure 1 provides a schematic diagram of the system excluding confounders and error terms.

**Figure 1:**
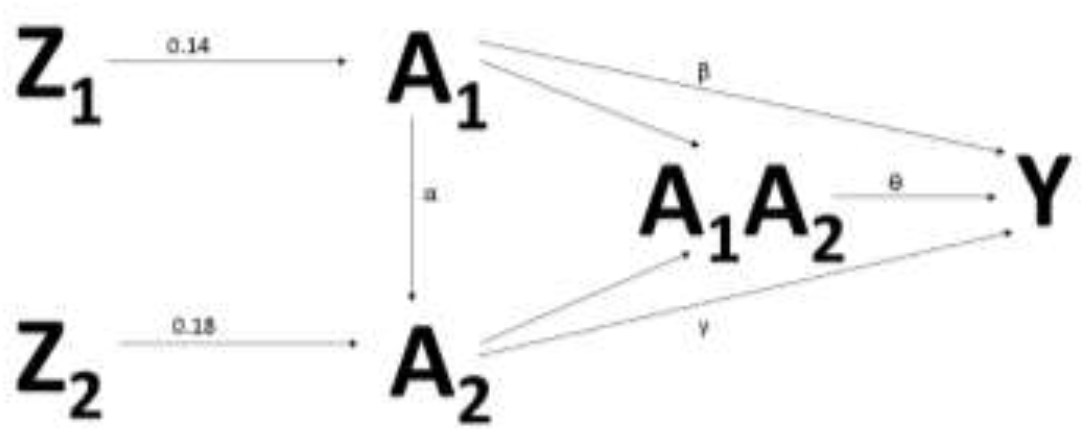
Schematic diagram of simulated data system, excluding confounder and error terms

**Figure 2:**
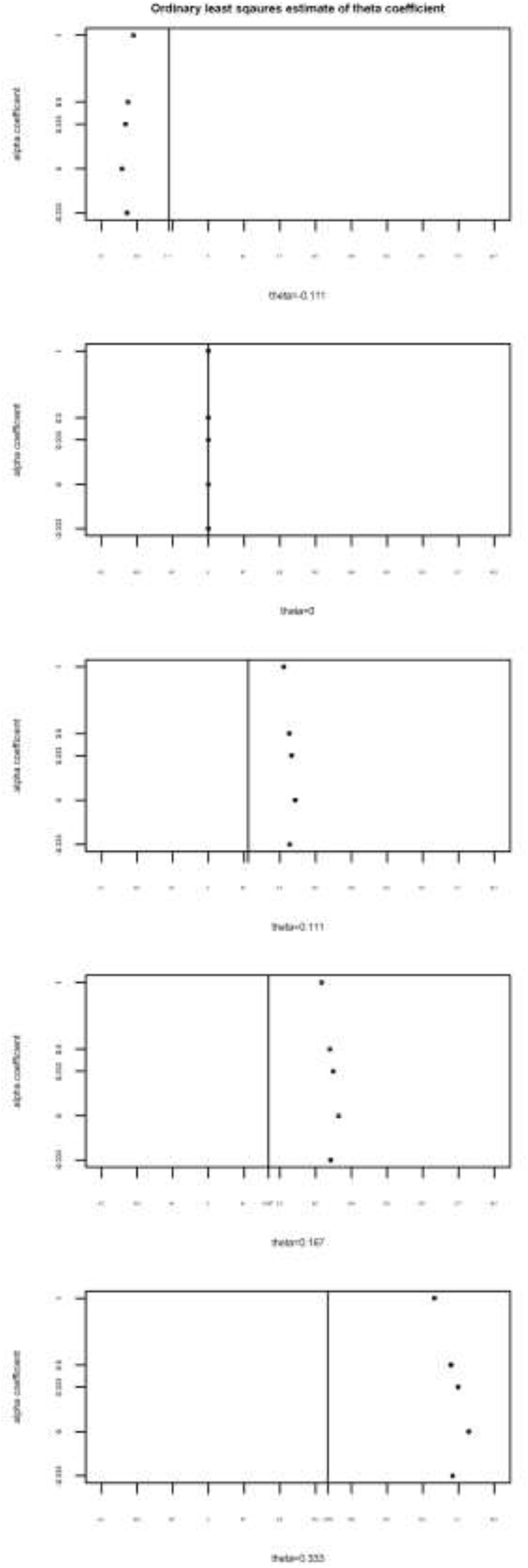
Coefficient estimates for ordinary least squares when N=100,000. Theta is the interaction coefficient (A_1_A_2_), and alpha is the effect of A1 on A2. The vertical lines show the true theta (interaction) coefficient and the OLS estimates with their Monte Carlo 95% confidence interval are overlaid.

Standard random normal variables were generated to represent error terms (U, V and E), confounders (C) and genetic instruments (Z_1_ and Z_2_) (1 - 3). Z_1_ is the genetic risk score for exposure 1 (A_1_) and Z_2_ is the (independent) genetic risk score for exposure 2 (A_2_).

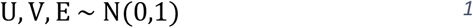

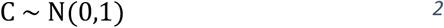

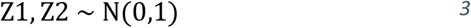

The random variables were used to generate values for each exposure (A_1_ and A_2_) and the outcome (Y), as shown in Equation 4.

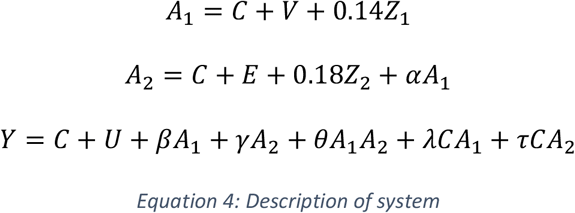

Interactions between the confounder variable (C) and A_1_ and A_2_ separately were included in the Y equation (10, 11). A_1_ was allowed to affect A_2_, implying the effect of A_1_ on Y is partially mediated by A_2_. We tested 25 alternative scenarios, which represented permutations of 5 coefficients for A_1_ in the A_2_ equation (α in Equation 4, took values −0.333, 0, 0.333, 0.5 and 1), 5 coefficients for A_1_ and A_2_ in the Y equation (β and γ in equation 4, took values −0.333, 0, 0.333, 0.5 and 1) and 5 coefficients for A_1_A_2_, CA_1_ and CA_2_ in the Y equation (θ, λ and τ in equation 4, took values −0.111, 0, 0.111, 0.167 and 0.333).

25 combinations of these parameters were tested, which were representative of the parameter space, including scenarios with and without an interaction between A_1_ and A_2_, and with and without a causal effect of A_1_ on A_2_. A table of parameter permutations is provided in the Supplement (Section 1). The interaction parameters in the Y equation (θ, λ and τ) were set to be one third of the main effect’s parameters (β and γ) to impose a limit on the variance explained by interaction terms.

The coefficients of 0.14 and 0.18 for the effect of the instrument on the exposures (A_1_ and A_2_) in Equation 4 were chosen to impose an approximate proportion of variance explained by the genetic risk scores of 1-2%. This aims to emulate the R^2^ from a typical genetic risk score in the literature.

We simulated twelve sample sizes: N=10,000 to N=100,000 (in steps of 10,000), N=500,000 and N=1,000,000. Models were run 1,000 times for each sample size and results pooled across repeats. All regression analyses and subsequent testing were performed using functions in the R systemfit package (12) version 1.1-22, the AER package (13) version 1.2-5, or using R’s lm function.

### Two-stage least squares estimation

Instrumental variable regression in AER (13) (using the *ivreg* command) or systemfit (12) was implemented. This uses the instruments Z to generate predicted values for A_1_, A_2_ and A_1_A_2_ (interaction coefficient), and regresses these predicted values against Y. The standard errors in the second stage are adjusted to account for the fact that the A_1_, A_2_ and A_1_A_2_ are predictions based on the instruments Z. For all simulations, we evaluated two IV: i) the genetic risk score for each exposure (Z_1_ and Z_2_) and the product of the two genetic risk scores, i.e. Z=(Z_1_,Z_2_,Z_1_Z_2_), and ii) an IV that allows for when the first exposure has a causal effect on the second exposure, Z= (Z_1_,Z_2_,Z_1_Z_2_,Z_1_Z_1_). The justification for including the square of the genetic risk score for exposure 1 (Z_1_Z_1_) is explained in full in Supplemental Methods section 11. The instruments are described as vectors here because all of the instruments are included in the first stage regression models in 2SLS to predict the values of each of the exposures.

### Factorial MR estimation

We modelled the factorial MR approach, by splitting the two genetic instruments Z_1_ and Z_2_ at the median to generate four subgroups representing low A_1_-low A_2_, low A_1_-high A_2_, high A_1_-low A_2_ and high A_1_-high A_2_. An ordinary least squares regression of the outcome against these four categories was performed (low-low was treated as the reference category) and a linear combination of the regression coefficients tested using an F statistic and Wald test (using the-linearHypothesis-R function from the car package (14)) to conclude presence or absence of an interaction between A_1_ and A_2_ on Y. The linear hypothesis we tested was that the sum of the regression coefficients for the low A_1_-high A_2_ and the high A_1_-low A_2_ categories equalled the high A_1_-high A_2_ coefficient. Rejection of this hypothesis at p=0.05 was regarded as suggestive of an interaction between the two exposures. This formal interaction test was not included in the original application of factorial MR (5) but we have adopted it to enable comparisons with the 2SLS approaches.

### Evaluation of approaches to estimate interaction

Coefficients and standard errors were collapsed across the 1,000 simulation runs; the regression coefficients were averaged to give 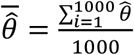. This provides a measure of the bias. The 95% Monte Carlo confidence interval (MC CI) (15) for these coefficients was calculated using the formula 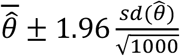, where sd is the sample standard deviation. If the 95% MC CI includes the true value we conclude that the regression coefficient from this estimating procedure is unbiased (15). We also examined the mean estimated standard error of 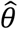 across simulation repeats and calculated the standard deviation of the regression coefficients 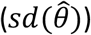 across repeats to evaluate precision (16). As the number of simulation repeats tends to infinity, 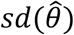 estimates the true precision of the estimator and deviation from the mean estimated standard error of 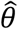 is indicative of bias in the estimate of precision. We also considered power, type I error and coverage. Power was defined as detecting an interaction at the 5% level (two-sided test) when an interaction (θ≠0) was present. A power of 68% therefore represents 680 of the 1,000 repeats detecting an interaction at p≤0.05 when the true interaction parameter was non-zero. Type I error was similarly calculated by counting the number of repeats where an interaction was detected at p≤0.05 when the true parameter value was zero. ‘Interaction detection’ was defined as the 95% confidence interval not including 0 for the 2SLS and observational linear regression approaches. Coverage was calculated as the percentage of repeats where the 95% confidence interval contained the true parameter value. Due to the nature of the factorial design, coverage could not be calculated because a confidence interval for the interaction parameter is not available. However, power and type I error were calculated by testing for an interaction using the linear contrast described previously.

### Sensitivity analyses

#### i. Pleiotropy

Pleiotropy refers to deviation from the assumption that an IV only affects the outcome through the exposure (17). A sensitivity analysis allowing for a pleiotropic effect of A_2_’s instrument on A_1_ was run with N=500,000. We compared the performance of the 2SLS approach using the instrument Z=(Z_1_,Z_2_,Z_1_Z_2_,Z_1_Z_1_) under a model of no pleiotropy versus pleiotropy using data from 1,000 random variable simulations generated by ten new seeds. We also considered the power and type I error of the factorial MR approach. The updated A_1_ equation for the pleiotropic scenario is given by equation 5.

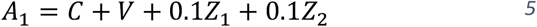

#### ii. Instrumental variable strength

With N=50,000, we changed the coefficients of Z_1_ and Z_2_ in the A_1_ and A_2_ equations respectively (Equation 6) to increase the percentage variance explained by the instruments (achieving 2.4-8.9% for A_2_).

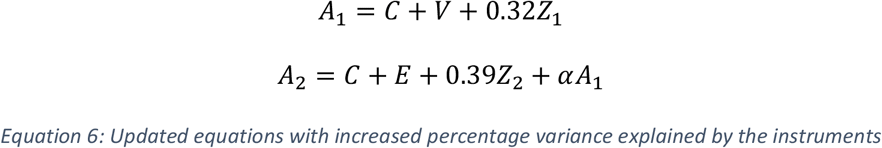

#### iii. Robust standard errors

To explore whether the use of robust standard errors (e.g. to account for deviations from assumptions of normality or other potential violations of assumptions) affected our findings, the observational and 2SLS regressions (using Z=(Z_1_,Z_2_,Z_1_Z_2_,Z_1_Z_1_)) in the main analysis were repeated with heteroskedastic robust standard errors using the *coeftest* R function or the *robust.se* R function (the former from the lmtest (18) R package and the latter from the ivpack (19) R package) with the results examined at N=50,000 and N=500,000.

#### iv. Weak instrument bias

The Sanderson-Windmeijer F statistics (20) were examined after the 2SLS regressions in the main analysis to test for weak-instrument bias and the correlation with sample size.

## ILLUSTRATIVE EXAMPLE

We used data from the UK Biobank (9), a cohort of middle aged to older individuals, to estimate the interactive effect of body mass index (BMI) and educational attainment on systolic blood pressure using ordinary least squares regression, 2SLS (with both Z=(Z_1_,Z_2_,Z_1_Z_2_) and Z=(Z_1_,Z_2_,Z_1_Z_2_,Z_1_Z_1_)) and factorial MR. We assumed a causal effect of years of education on BMI. All phenotype measures were taken from the baseline assessment clinic in 2006-2010. Educational attainment was used as a continuous variable of years of completed education (data field 845). BMI was calculated as weight in kilograms divided by height in metres squared. Systolic blood pressure was taken from the second reading, generally measured using an Omron 705 IT electronic blood pressure monitor (OMRON Healthcare Europe B. V. Kruisweg 577 2132 NA Hoofddorp). Pregnant (or possibly pregnant) women (n=372) were removed from the analysis because of changes to both BMI and blood pressure during pregnancy. We created genetic risk scores for BMI and years of schooling for all UK Biobank participants using the results of recent GWAS studies (21, 22). The discovery datasets excluded UK Biobank (the SNPs for years of schooling were taken from the dataset EduYears_Discovery_5000.txt available on the SSGAC website https://www.thessgac.org/data). We selected SNPs from each study with a P value below genome-wide significance (P ≤ 5 × 10-8). We pruned SNPs from the BMI genetic risk score to only include those independent from the education genetic risk score. Prior to pruning, the BMI genetic risk score included 69 SNPs and after pruning, this reduced to 67, while the education genetic risk score included 74 SNPs. Each genetic risk score was calculated as the sum of the effect alleles for all SNPs associated with BMI or education, with each SNP weighted by the regression coefficient from the GWAS from which the SNP was identified. We searched for proxy SNPs with an R^2^ above 0.8 for any SNP missing from UK Biobank using genotype data from European individuals (CEU) from phase 3 (version 5) of the 1000 Genomes project (23).

Age in whole years during baseline assessment centre attendance and sex were used as covariates in all analyses. Genetic analyses were also adjusted for the first 40 principal components to minimise the effects of population stratification. Sensitivity analyses were run with and without adjusting for genotyping array. Further information on UK Biobank and the quality control filtering applied to the data in this manuscript is provided in the Supplement. Quality Control filtering of the UK Biobank data was conducted by R.Mitchell, G.Hemani, T.Dudding, L.Paternoster (24).

Instrumental variable regression was implemented in Stata (StataCorp, Texas) (25) using ivreg2 (26). Factorial MR and observational analyses were implemented using-regress-.STATA’s-lincom-command was used to combine coefficients for the purposes of interaction detection assessment for the factorial MR approach. The main effects for BMI and years of schooling were included in the linear regression and instrumental variable models. Details of the predictors used in each model are provided in the supplement (Section 11.3).

## RESULTS

### Simulation study

#### Bias in estimates of the interaction coefficient

Ordinary least squares analyses revealed clear bias when the interaction coefficient was non-zero (**Error! Not a valid bookmark self-reference.** N=100,000, Supplementary Figures in Sections 13.1-13.2 N=50,000 and N=500,000). The 2SLS estimates were markedly improved, but there was still some evidence of bias for some parameter combinations at N=50,000 and N=100,000 for both Z=(Z_1_,Z_2_,Z_1_Z_2_,) and Z=(Z_1_,Z_2_,Z_1_Z_2_,Z_1_Z_1_). At N=50,000, 2SLS using Z=(Z_1_,Z_2_,Z_1_Z_2_,Z_1_Z_1_) produced coefficient estimates that were generally closer to the true value and had a smaller standard error and mean standard error compared with using Z=(Z_1_,Z_2_,Z_1_Z_2_) (Figures 3 and 4; Supplementary Figures in section 13.3-13.4). At N=500,000, 2SLS estimates were very close to the true parameter values, with minimal differences between Z=(Z_1_,Z_2_,Z_1_Z_2_,Z_1_Z_1_) and Z=(Z_1_,Z_2_,Z_1_Z_2_) (Figure 5; Supplementary Figure in section 13.5). Full results for bias analysis are presented in Supplementary Tables 2.1-5.5.

**Figure 3:**
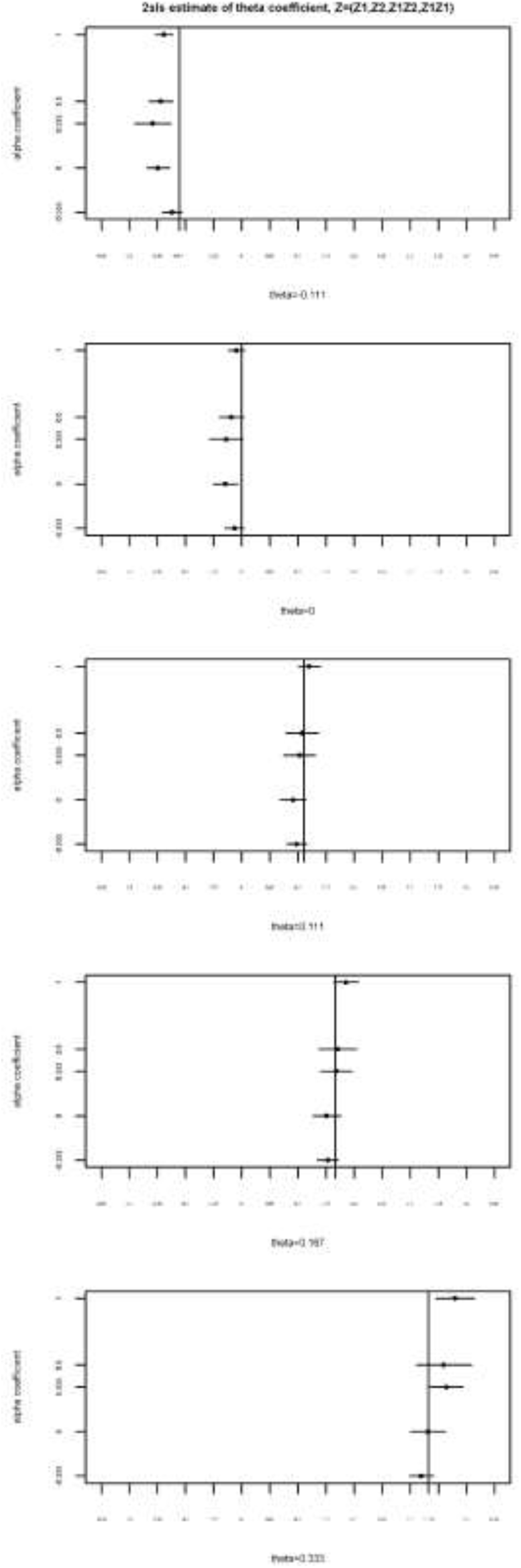
Coefficient estimates for 2SLS using Z=(Z_1_,Z_2_,Z_1_Z_2_,Z_1_Z_1_) when N=50,000. Theta is the interaction coefficient (A_1_A_2_), and alpha is the effect of A1 on A2. The vertical lines show the true theta (interaction) coefficient and the 2SLS estimates with their Monte Carlo 95% confidence interval are overlaid.

**Figure 4:**
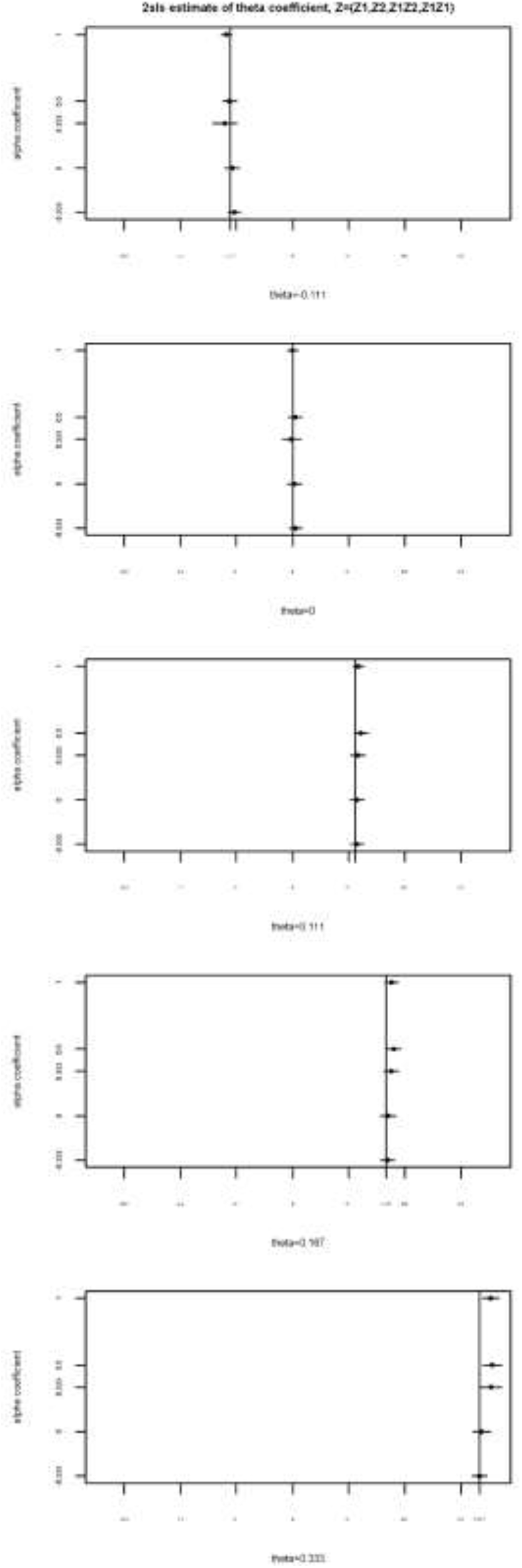
Coefficient estimates for 2SLS using Z=(Z_1_,Z_2_,Z_1_Z_2_,Z_1_Z_1_) when N=100,000. Theta is the interaction coefficient (A_1_A_2_), and alpha is the effect of A1 on A2. The vertical lines show the true theta (interaction) coefficient and the 2SLS estimates with their Monte Carlo 95% confidence interval are overlaid.

**Figure 5:**
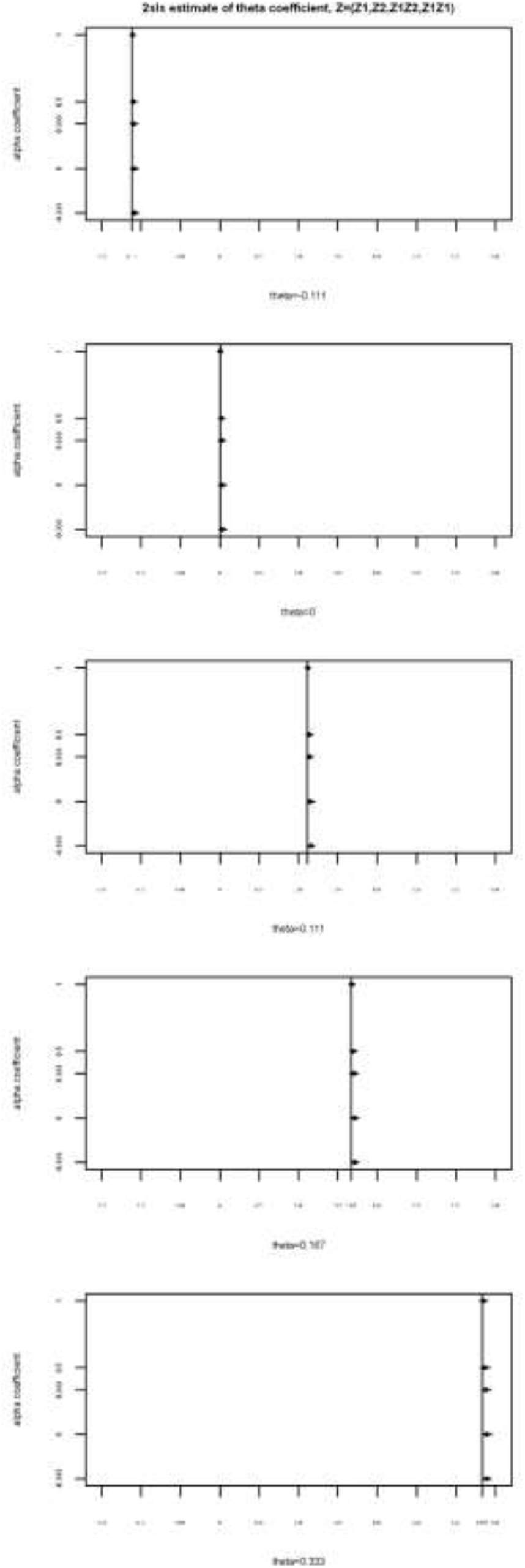
Coefficient estimates for 2SLS using Z=(Z_1_,Z_2_,Z_1_Z_2_,Z_1_Z_1_) when N=500,000. Theta is the interaction coefficient (A_1_A_2_), and alpha is the effect of A1 on A2. The vertical lines show the true theta (interaction) coefficient and the 2SLS estimates with their Monte Carlo 95% confidence interval are overlaid.

#### Power, coverage and type I error for the interaction coefficient

Power for factorial MR was extremely low; at N=50,000 power ranged from 5%-7%, and at N=500,000, the power of factorial MR was between 8-23%. Power for 2SLS was considerably higher than for factorial MR, but was still low at smaller sample sizes. At N=50,000, 2SLS using Z=(Z_1_,Z_2_,Z_1_Z_2_,Z_1_Z_1_) did not reach a power greater than 55%, and was generally much lower than this with lowest power 7%. Power for 2SLS using Z=(Z_1_,Z_2_,Z_1_Z_2_,Z_1_Z_1_) at N=500,000 ranged from 29.7% - 92.9%. Coverage for 2SLS using Z=(Z_1_,Z_2_,Z_1_Z_2_,Z_1_Z_1_) ranged from 90.6 to 99.6% for N=50,000 and 93.2 to 96.4% for N=500,000. Coverage and power metrics for FMR vs 2SLS are plotted in Figure 6 (N=50,000) and Figure 7 (N=500,000). Power and coverage were similar when using Z=(Z_1_,Z_2_,Z_1_Z_2_,Z_1_Z_1_) or using Z=(Z_1_,Z_2_,Z_1_Z_2_). When θ=0 (no interaction present), OLS had a Type I error 4-5% at N=500,000. In comparison, the rate was 5.8% for factorial MR and ranged from 4 to 6% for 2SLS using Z=(Z_1_,Z_2_,Z_1_Z_2_,Z_1_Z_1_).

**Figure 6:**
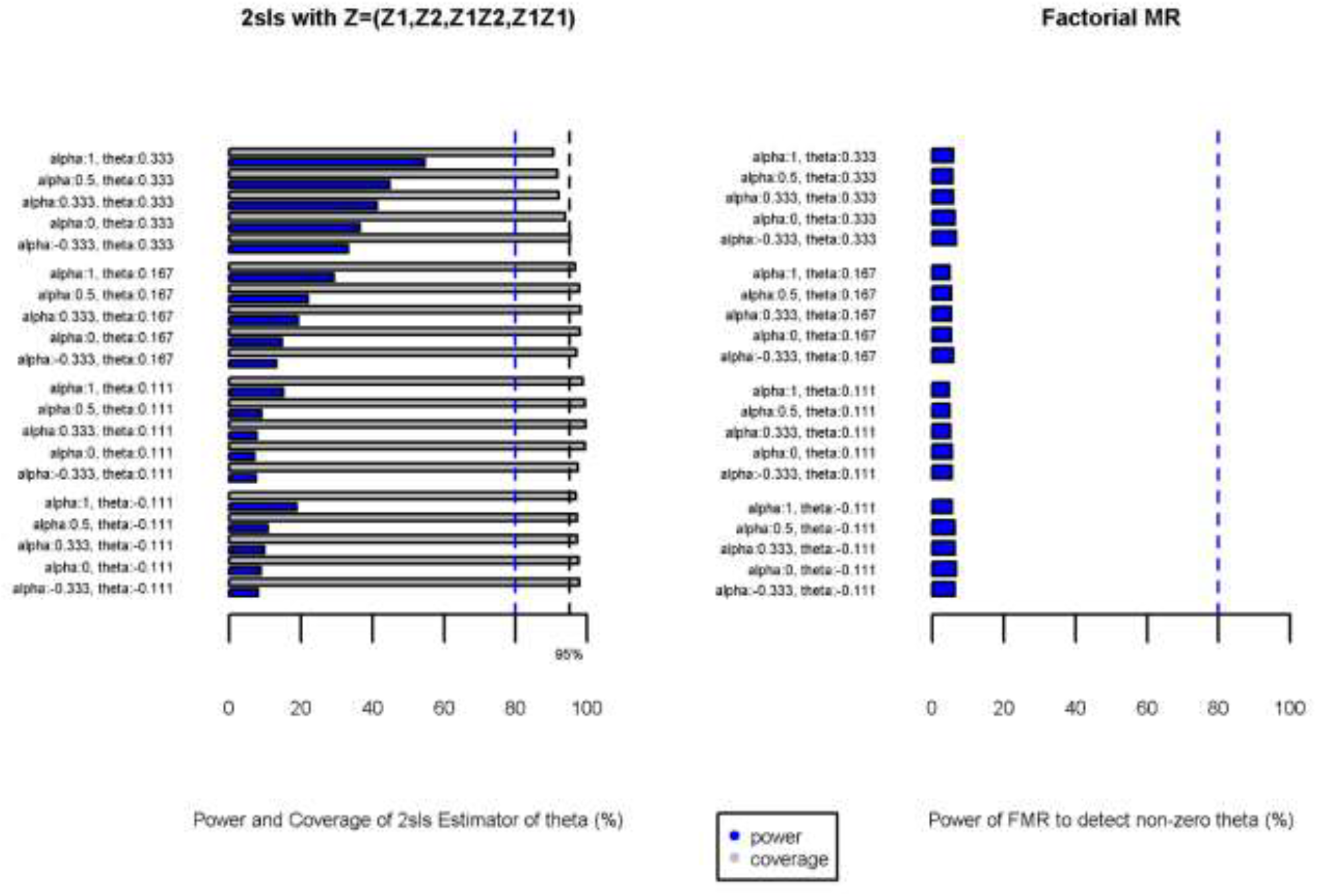
2SLS (power and coverage) versus factorial MR (power) when N=50,000. Theta is the interaction coefficient (A_1_A_2_), and alpha is the effect of A1 on A2.

**Figure 7:**
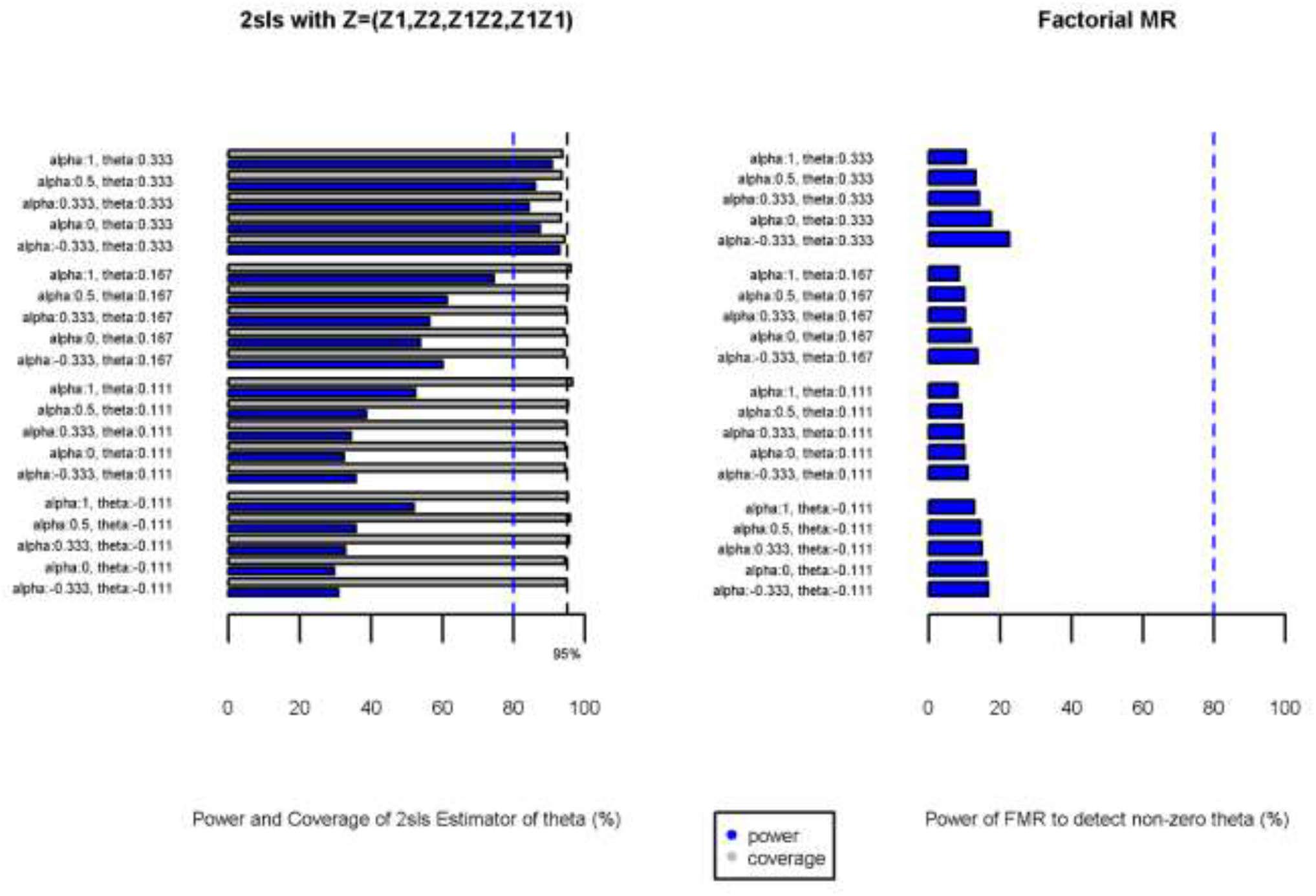
2SLS (power and coverage) versus factorial MR (power) when N=500,000. Theta is the interaction coefficient (A_1_A_2_), and alpha is the effect of A1 on A2.

#### Sensitivity analyses

An additional simulation was run at N=500,000 where A_2_’s instrument had a pleiotropic effect on A_1_. The results are presented in sections 7 and 8 of the Supplement. Under some parameter combinations the existence of pleiotropy negatively affected the performance of the 2SLS using Z=(Z_1_,Z_2_,Z_1_Z_2_,Z_1_Z_1_) and factorial MR estimators, potentially because the instruments were weaker. We also increased the instrument coefficients in the data generating model in a further simulation at N=50,000 to mimic stronger instrumental variables. As expected, the power of 2SLS using Z=(Z_1_,Z_2_,Z_1_Z_2_,Z_1_Z_1_) to detect the interaction was higher with greater instrumental variable strength; full results in Supplement section 9. Repeating the main observational and 2SLS (using Z=(Z_1_,Z_2_,Z_1_Z_2_,Z_1_Z_1_)) using robust standard errors did not materially affect the results (Supplementary Tables 10.1-10.4). Examination of the Sanderson-Windmeijer F-statistic revealed that in smaller sample sizes the instruments were weak to predict the interaction term (Supplementary Table 11.1, Supplementary Figures in Section 11).

### Illustrative example: education, BMI and blood pressure

Data were available for all variables for N=306,593 participants. Neither factorial MR nor the 2SLS approaches (using Z=(Z_1_,Z_2_,Z_1_Z_2_,Z_1_Z_1_) or Z=(Z_1_,Z_2_,Z_1_Z_2_)) detected an interaction between BMI and years of schooling on systolic blood pressure, in contrast to the (age and sex adjusted) observational linear regression analysis which suggested a positive additive interaction; β=0.029 mmHg higher systolic blood pressure per unit increase in BMI (kg/m^2^)*years of schooling (p<0.0005). The genetic risk score for BMI explained 1.6% of the variance in BMI, and the education genetic risk score explained 0.5% of the variance in years of education. Full results are in Supplement section 11.

## DISCUSSION

Our simulation study showed that factorial MR has very low power to detect additive interactions. In contrast, an extension to multivariable MR (MVMR, an approach to estimate the effects of two or more exposures on an outcome, which allows for overlapping instruments between exposures) (27, 28) using genetic risk scores for each exposure and their product as the instrument in 2SLS had greater power. The 2SLS approach was generally unbiased, with reasonable coverage and type I error, but required large sample sizes and strong instrumental variables. In addition to having greater statistical power than factorial MR, 2SLS also has the advantage of being comparable with and providing estimates on the same scale as multivariable linear regression. Factorial MR has an implicit simplicity, which may be attractive to researchers wishing to present findings to clinical or non-statistical audiences, but our results suggest that in most cases, the 2SLS approach will be preferable. Within the 2SLS framework, we tested two instrumental variables based on the genetic risk scores for two exposures (Z_1_ and Z_2_): Z=(Z_1_,Z_2_,Z_1_Z_2_,Z_1_Z_1_) and Z=(Z_1_,Z_2_,Z_1_Z_2_). The former includes the square of the genetic risk score for the first exposure, motivated by situations when exposure 1 has a causal effect on exposure 2. We found Z=(Z_1_,Z_2_,Z_1_Z_2_,Z_1_Z_1_) to exhibit less bias and have a smaller standard error and mean standard error at smaller sample sizes (N=50,000 and N=100,000); at N=500,000 the differences between the instrumental variables were minimal both in the presence and absence of causal effects of exposure 1 on exposure 2.

Our findings of low statistical power in factorial MR are similar to the literature on factorial randomised controlled trials, which indicates that a fourfold increase in sample size is generally necessary to detect with the same power an interaction of the same magnitude as the main effect (29). In our simulations, power of the 2SLS approaches increased when the simulation was repeated with genetic risk scores that explained a higher percentage of the variance in the exposures; this suggests that the increased power we see with higher sample sizes may be partially driven by increasing instrument strength.

Our simulations were limited to a continuous outcome and interactions on the additive scale; further work is required to test and develop approaches for binary outcomes and multiplicative interactions. Our approach uses individual participant data; assessing interactions within two-sample MR would be challenging and requires further methodological development. We have focused on the ability of the 2SLS approaches to detect additive interaction; we have not investigated the implications of these approaches for the main effects of each exposure. Our main analysis assumed no pleiotropy but a sensitivity analysis showed that pleiotropic effects of the instruments for one exposure on the other can affect the results from 2SLS, likely through making the instruments weaker. Our simulations modelled scenarios when the first exposure has a causal effect on the second exposure, but we did not explore the potential for bidirectional causal effects.

In our illustrative example, in contrast to the ordinary least squares estimate, neither factorial MR nor either of the 2SLS approaches provided evidence of an interaction of BMI and years of schooling on systolic blood pressure. This could be because the observed interaction is induced by confounding. Alternatively, this could be because the 2SLS and factorial MR approaches are underpowered to detect the interactive effect. The sample size for the analyses exceeded 300,000, and the genetic risk scores explained 1.5% (BMI) and 0.5% (years of education) of the variation in the exposures, suggesting power in this example was reasonable for the 2SLS approaches and unlikely to be the explanation for the null finding. Our findings are interesting in the context of a recent study (6) that examined the interaction of genetic risk for body size and education instrumented by policy reform on diastolic and systolic blood pressure, finding weak evidence for an interaction on diastolic but not systolic blood pressure.

In summary, we have shown that factorial MR has low statistical power in most realistic scenarios, and that 2SLS using genetic risk scores for each exposure is a more powerful technique to estimate an unconfounded additive interaction of two continuous exposures. However, large sample sizes and instrument strength are crucial to conducting an analysis that is sufficiently powered to detect such effects.

## Acknowledgements

The authors would like to thank Dr Rhian Daniel for helpful comments during the earlier stages of the analysis.

Author contributions
All authors contributed to study design, analysis, interpretation of results and critically revised the manuscript. TLN performed the majority of the analysis and wrote the first draft of the manuscript.

## REFERENCES

1. Lawlor DA, Harbord RM, Sterne JAC, Timpson N, Davey Smith G. Mendelian randomization: Using genes as instruments for making causal inferences in epidemiology. Statistics in Medicine. 2008;27(8):1133–63.

2. Hernán MA, Robins JM. Instruments for Causal Inference: An Epidemiologist’s Dream? Epidemiology. 2006;17(4):360–72.

3. Sanderson E, Davey Smith G, Windmeijer F, Bowden J. An examination of multivariable Mendelian randomization in the single-sample and two-sample summary data settings. Int J Epidemiol. 2018.

4. VanderWeele TJ. A Unification of Mediation and Interaction: A 4-Way Decomposition. Epidemiology. 2014;25(5):749–61.

5. Ference BA, Majeed F, Penumetcha R, Flack JM, Brook RD. Effect of Naturally Random Allocation to Lower Low-Density Lipoprotein Cholesterol on the Risk of Coronary Heart Disease Mediated by Polymorphisms in NPC1L1, HMGCR, or Both. Journal of the American College of Cardiology. 2015;65(15):1552–61.

6. Barcellos SH, Carvalho LS, Turley P. Education can reduce health differences related to genetic risk of obesity. Proceedings of the National Academy of Sciences. 2018;115(42):E9765.

7. Wooldridge JM. Chapter 6: Additional Single-Equation Topics. Econometric Analysis of Cross Section and Panel Data. Second Edition ed. Cambridge, Massachusetts: The MIT Press; 2010. p. 133–4.

8. Bond SJ, White IR, Sarah Walker A. Instrumental variables and interactions in the causal analysis of a complex clinical trial. Stat Med. 2007;26(7):1473–96.

9. Allen NE, Sudlow C, Peakman T, Collins R. UK Biobank Data: Come and Get It. Science Translational Medicine. 2014;6(224):224ed4.

10. Yzerbyt VY, Muller D, Judd CM. Adjusting researchers’ approach to adjustment: On the use of covariates when testing interactions. Journal of Experimental Social Psychology. 2004;40(3):424–31.

11. Keller MC. Gene-by-environment interaction studies have not properly controlled for potential confounders: The problem and the (simple) solution. Biological psychiatry. 2014;75(1):10.1016/j.biopsych.2013.09.006.

12. Henningsen A, Hamann JD. systemfit: A Package for Estimating Systems of Simultaneous Equations in R. Journal of Statistical Software. 2007;23(4):1–40.

13. Kleiber C, Zeileis A. Applied Econometrics with R. New York: Springer-Verlag; 2008.

14. Fox J, Weisberg S. An {R} Companion to Applied Regression, Second Edition. Thousand Oaks, CA: Sage; 2011.

15. Hughes RA, Sterne JAC, Tilling K. Comparison of imputation variance estimators. Statistical Methods in Medical Research. 2014;25(6):2541–57.

16. Burton A, Altman DG, Royston P, Holder RL. The design of simulation studies in medical statistics. Statistics in Medicine. 2006;25(24):4279–92.

17. Hemani G, Bowden J, Davey Smith G. Evaluating the potential role of pleiotropy in Mendelian randomization studies. Hum Mol Genet. 2018;27(R2):R195–r208.

18. Zeileis A, Hothorn T. Diagnostic Checking in Regression Relationships. R News. 2002;2(3):7–10.

19. Jiang Y, Small D. ivpack: Instrumental variable estimation. 2014.

20. Sanderson E, Windmeijer F. A weak instrument F-test in linear IV models with multiple endogenous variables. Journal of Econometrics. 2016;190(2):212–21.

21. Locke AE. Genetic studies of body mass index yield new insights for obesity biology. Nature. 2015;518.

22. Okbay A, Beauchamp JP, Fontana MA, Lee JJ, Pers TH, Rietveld CA, et al. Genome-wide association study identifies 74 loci associated with educational attainment. Nature. 2016;533:539.

23. Abecasis GR, Auton A, Brooks LD, DePristo MA, Durbin RM, Handsaker RE, et al. An integrated map of genetic variation from 1,092 human genomes. Nature. 2012;491(7422):56–65.

24. Mitchell R, Hemani G, Dudding T, Paternoster L. UK Biobank Genetic Data: MRC-IEU Quality Control, Version 1. 2017.

25. StataCorp. Stata Statistical Software: Release 15. College Station, TX: StataCorp LLC; 2017.

26. Baum CF, Schaffer ME, Stillman S. Enhanced routines for instrumental variables/generalized method of moments estimation and testing. The Stata Journal. 2007;7(4):465–506.

27. Burgess S, Thompson SG. Multivariable Mendelian Randomization: The Use of Pleiotropic Genetic Variants to Estimate Causal Effects. American Journal of Epidemiology. 2015;181(4):251–60.

28. Sanderson E, Davey Smith G, Windmeijer F, Bowden J. An examination of multivariable Mendelian randomization in the single sample and two-sample summary data settings. bioRxiv. 2018.

29. Brookes ST, Whitley E, Peters TJ, Mulheran PA, Egger M, Davey Smith G. Subgroup analyses in randomised controlled trials: quantifying the risks of false-positives and false-negatives. Health technology assessment (Winchester, England). 2001;5(33):1–56.

