## Supplementary Text for "Using genetic instruments to estimate interactions in Mendelian Randomization studies"

Supplemental Material

Teri-Louise North[1], Neil M Davies[1], Sean Harrison[1], Alice R Carter[1], Gibran Hemani[1], Eleanor Sanderson[1], Kate Tilling[1], Laura D Howe[1]

1 MRC Integrative Epidemiology Unit at the University of Bristol, Population Health Sciences, University of Bristol, Bristol, UK

### 1 PARAMETER PERMUTATIONS

| α | β | γ | θ | λ | τ |
|---|---|---|---|---|---|
| -0.333 | -0.333 | -0.333 | -0.111 | -0.111 | -0.111 |
| 0 | -0.333 | -0.333 | -0.111 | -0.111 | -0.111 |
| 0.333 | -0.333 | -0.333 | -0.111 | -0.111 | -0.111 |
| 0.5 | -0.333 | -0.333 | -0.111 | -0.111 | -0.111 |
| 1 | -0.333 | -0.333 | -0.111 | -0.111 | -0.111 |
| -0.333 | 0 | 0 | 0.000 | 0.000 | 0.000 |
| 0 | 0 | 0 | 0.000 | 0.000 | 0.000 |
| 0.333 | 0 | 0 | 0.000 | 0.000 | 0.000 |
| 0.5 | 0 | 0 | 0.000 | 0.000 | 0.000 |
| 1 | 0 | 0 | 0.000 | 0.000 | 0.000 |
| -0.333 | 0.333 | 0.333 | 0.111 | 0.111 | 0.111 |
| 0 | 0.333 | 0.333 | 0.111 | 0.111 | 0.111 |
| 0.333 | 0.333 | 0.333 | 0.111 | 0.111 | 0.111 |
| 0.5 | 0.333 | 0.333 | 0.111 | 0.111 | 0.111 |
| 1 | 0.333 | 0.333 | 0.111 | 0.111 | 0.111 |
| -0.333 | 0.5 | 0.5 | 0.167 | 0.167 | 0.167 |
| 0 | 0.5 | 0.5 | 0.167 | 0.167 | 0.167 |
| 0.333 | 0.5 | 0.5 | 0.167 | 0.167 | 0.167 |
| 0.5 | 0.5 | 0.5 | 0.167 | 0.167 | 0.167 |
| 1 | 0.5 | 0.5 | 0.167 | 0.167 | 0.167 |
| -0.333 | 1 | 1 | 0.333 | 0.333 | 0.333 |
| 0 | 1 | 1 | 0.333 | 0.333 | 0.333 |
| 0.333 | 1 | 1 | 0.333 | 0.333 | 0.333 |
| 0.5 | 1 | 1 | 0.333 | 0.333 | 0.333 |
| 1 | 1 | 1 | 0.333 | 0.333 | 0.333 |

### 2 ORDINARY LEAST SQUARES ASSOCIATION

#### 2.1 THETA=-0.111

| Modelled mediator coefficient | Modelled interaction coefficient | Sample size | Mean observational interaction effect estimate | Standard deviation of observational estimate | Mean estimated standard error of observational estimate | Standard error of the bias (standard deviation of observational estimate/sqrt(1000)) | Power of observational estimator (%) | Coverage of observational estimator (%) |
|---|---|---|---|---|---|---|---|---|
| α = -0.333 | θ = -0.111 | 10,000 | -0.22741 | 0.00702 | 0.006463 | 0.000222 | 100 | 0 |
| | | 20,000 | -0.22765 | 0.004721 | 0.004562 | 0.000149 | 100 | 0 |
| | | 30,000 | -0.22775 | 0.003883 | 0.003726 | 0.000123 | 100 | 0 |
| | | 40,000 | -0.22761 | 0.003451 | 0.003225 | 0.000109 | 100 | 0 |
| | | 50,000 | -0.22743 | 0.003007 | 0.002885 | 9.51E-05 | 100 | 0 |
| | | 60,000 | -0.22752 | 0.002805 | 0.002633 | 8.87E-05 | 100 | 0 |
| | | 70,000 | -0.22744 | 0.002672 | 0.002438 | 8.45E-05 | 100 | 0 |
| | | 80,000 | -0.22748 | 0.002503 | 0.00228 | 7.92E-05 | 100 | 0 |
| | | 90,000 | -0.22742 | 0.00222 | 0.002149 | 7.02E-05 | 100 | 0 |
| | | 100,000 | -0.22743 | 0.002063 | 0.00204 | 6.52E-05 | 100 | 0 |
| | | 500,000 | -0.22747 | 0.000959 | 0.000912 | 3.03E-05 | 100 | 0 |
| | | 1,000,000 | -0.22753 | 0.000681 | 0.000645 | 2.15E-05 | 100 | 0 |
| α = 0 | θ = -0.111 | 10,000 | -0.24269 | 0.005632 | 0.0052 | 0.000178 | 100 | 0 |
| No mediation | | 20,000 | -0.24272 | 0.003767 | 0.00367 | 0.000119 | 100 | 0 |
| | | 30,000 | -0.24283 | 0.003072 | 0.002997 | 9.72E-05 | 100 | 0 |
| | | 40,000 | -0.24274 | 0.002693 | 0.002594 | 8.52E-05 | 100 | 0 |
| | | 50,000 | -0.24255 | 0.002369 | 0.002321 | 7.49E-05 | 100 | 0 |
| | | 60,000 | -0.24261 | 0.00223 | 0.002119 | 7.05E-05 | 100 | 0 |
| | | 70,000 | -0.24255 | 0.002094 | 0.001962 | 6.62E-05 | 100 | 0 |
| | | 80,000 | -0.24262 | 0.00196 | 0.001835 | 6.20E-05 | 100 | 0 |
| | | 90,000 | -0.24256 | 0.001761 | 0.00173 | 5.57E-05 | 100 | 0 |
| | | 100,000 | -0.24256 | 0.001677 | 0.001641 | 5.30E-05 | 100 | 0 |
| | | 500,000 | -0.2426 | 0.000763 | 0.000734 | 2.41E-05 | 100 | 0 |

| | | | | | | | | |
|---|---|---|---|---|---|---|---|---|
| | | 1,000,000 | -0.24263 | 0.000534 | 0.000519 | 1.69E-05 | 100 | 0 |
| α = 0.333 | θ = -0.111 | 10,000 | -0.23253 | 0.004377 | 0.003991 | 0.000138 | 100 | 0 |
| | | 20,000 | -0.2325 | 0.002953 | 0.002817 | 9.34E-05 | 100 | 0 |
| | | 30,000 | -0.23258 | 0.002394 | 0.002301 | 7.57E-05 | 100 | 0 |
| | | 40,000 | -0.23254 | 0.002085 | 0.001991 | 6.59E-05 | 100 | 0 |
| | | 50,000 | -0.23239 | 0.001827 | 0.001782 | 5.78E-05 | 100 | 0 |
| | | 60,000 | -0.23243 | 0.001707 | 0.001626 | 5.40E-05 | 100 | 0 |
| | | 70,000 | -0.23239 | 0.00161 | 0.001506 | 5.09E-05 | 100 | 0 |
| | | 80,000 | -0.23246 | 0.001513 | 0.001409 | 4.78E-05 | 100 | 0 |
| | | 90,000 | -0.23241 | 0.001382 | 0.001328 | 4.37E-05 | 100 | 0 |
| | | 100,000 | -0.23241 | 0.001335 | 0.00126 | 4.22E-05 | 100 | 0 |
| | | 500,000 | -0.23243 | 0.000595 | 0.000563 | 1.88E-05 | 100 | 0 |
| | | 1,000,000 | -0.23244 | 0.000413 | 0.000398 | 1.31E-05 | 100 | 0 |
| α = 0.5 | θ = -0.111 | 10,000 | -0.22614 | 0.003901 | 0.003527 | 0.000123 | 100 | 0 |
| | | 20,000 | -0.22611 | 0.002642 | 0.00249 | 8.35E-05 | 100 | 0 |
| | | 30,000 | -0.22618 | 0.00214 | 0.002033 | 6.77E-05 | 100 | 0 |
| | | 40,000 | -0.22615 | 0.001861 | 0.00176 | 5.89E-05 | 100 | 0 |
| | | 50,000 | -0.22602 | 0.001625 | 0.001575 | 5.14E-05 | 100 | 0 |
| | | 60,000 | -0.22605 | 0.001512 | 0.001437 | 4.78E-05 | 100 | 0 |
| | | 70,000 | -0.22602 | 0.001432 | 0.001331 | 4.53E-05 | 100 | 0 |
| | | 80,000 | -0.22608 | 0.001349 | 0.001245 | 4.27E-05 | 100 | 0 |
| | | 90,000 | -0.22604 | 0.001237 | 0.001174 | 3.91E-05 | 100 | 0 |
| | | 100,000 | -0.22604 | 0.001201 | 0.001114 | 3.80E-05 | 100 | 0 |
| | | 500,000 | -0.22605 | 0.000533 | 0.000498 | 1.68E-05 | 100 | 0 |
| | | 1,000,000 | -0.22606 | 0.000369 | 0.000352 | 1.17E-05 | 100 | 0 |
| α = 1 | θ = -0.111 | 10,000 | -0.21067 | 0.002929 | 0.002577 | 9.26E-05 | 100 | 0 |
| | | 20,000 | -0.21064 | 0.002004 | 0.001819 | 6.34E-05 | 100 | 0 |
| | | 30,000 | -0.21068 | 0.001622 | 0.001486 | 5.13E-05 | 100 | 0 |
| | | 40,000 | -0.21067 | 0.001408 | 0.001286 | 4.45E-05 | 100 | 0 |
| | | 50,000 | -0.21058 | 0.00122 | 0.001151 | 3.86E-05 | 100 | 0 |
| | | 60,000 | -0.21061 | 0.001124 | 0.00105 | 3.56E-05 | 100 | 0 |
| | | 70,000 | -0.21059 | 0.001075 | 0.000972 | 3.40E-05 | 100 | 0 |

| | | | | | | | | |
|---|---|---|---|---|---|---|---|---|
| | | 80,000 | -0.21063 | 0.001019 | 0.00091 | 3.22E-05 | 100 | 0 |
| | | 90,000 | -0.2106 | 0.000936 | 0.000858 | 2.96E-05 | 100 | 0 |
| | | 100,000 | -0.2106 | 0.00092 | 0.000814 | 2.91E-05 | 100 | 0 |
| | | 500,000 | -0.21061 | 0.000404 | 0.000364 | 1.28E-05 | 100 | 0 |
| | | 1,000,000 | -0.21061 | 0.00028 | 0.000257 | 8.85E-06 | 100 | 0 |

#### 2.2 THETA=0

| Modelled mediator coefficient | Modelled interaction coefficient | Sample size | Mean observational interaction effect estimate | Standard deviation of observational estimate | Mean estimated standard error of observational estimate | Standard error of the bias (standard deviation of observational estimate/sqrt(1000)) | Type I error of observational estimator (%) | Coverage of observational estimator (%) |
|---|---|---|---|---|---|---|---|---|
| α = -0.333 | θ = 0 | 10,000 | -6.49E-05 | 0.006435 | 0.006363834 | 0.000203 | 4.8 | 95.2 |
| | | 20,000 | -0.00016 | 0.004422 | 0.004492431 | 0.00014 | 4.6 | 95.4 |
| | | 30,000 | -0.00026 | 0.003624 | 0.003668848 | 0.000115 | 4.8 | 95.2 |
| | | 40,000 | -0.00015 | 0.003227 | 0.003175481 | 0.000102 | 5 | 95 |
| | | 50,000 | 5.13E-05 | 0.002801 | 0.002840981 | 8.86E-05 | 4.6 | 95.4 |
| | | 60,000 | -3.77E-06 | 0.002648 | 0.002593147 | 8.37E-05 | 4.7 | 95.3 |
| | | 70,000 | 6.02E-05 | 0.002449 | 0.002401261 | 7.74E-05 | 4.7 | 95.3 |
| | | 80,000 | 5.83E-06 | 0.002283 | 0.002245631 | 7.22E-05 | 5.4 | 94.6 |
| | | 90,000 | 5.13E-05 | 0.002105 | 0.00211651 | 6.66E-05 | 4.8 | 95.2 |
| | | 100,000 | 8.12E-05 | 0.001922 | 0.002008939 | 6.08E-05 | 4 | 96 |
| | | 500,000 | 8.71E-06 | 0.000888 | 0.0008983 | 2.81E-05 | 4.5 | 95.5 |
| | | 1,000,000 | -3.41E-05 | 0.000631 | 0.000635118 | 2.00E-05 | 4.7 | 95.3 |
| α = 0 | θ = 0 | 10,000 | -0.0002 | 0.005286 | 0.005132689 | 0.000167 | 4.9 | 95.1 |
| No mediation | | 20,000 | -0.00011 | 0.003601 | 0.003622746 | 0.000114 | 4.1 | 95.9 |
| | | 30,000 | -0.0002 | 0.002952 | 0.002958611 | 9.34E-05 | 5.3 | 94.7 |
| | | 40,000 | -0.00015 | 0.002574 | 0.002560852 | 8.14E-05 | 4.5 | 95.5 |
| | | 50,000 | 4.61E-05 | 0.002298 | 0.002290979 | 7.27E-05 | 5.1 | 94.9 |
| | | 60,000 | 7.78E-06 | 0.002122 | 0.00209099 | 6.71E-05 | 4.6 | 95.4 |
| | | 70,000 | 6.08E-05 | 0.001948 | 0.001936354 | 6.16E-05 | 4.4 | 95.6 |
| | | 80,000 | -1.88E-05 | 0.001813 | 0.001811084 | 5.73E-05 | 4.5 | 95.5 |
| | | 90,000 | 2.14E-05 | 0.001691 | 0.001707118 | 5.35E-05 | 5.7 | 94.3 |
| | | 100,000 | 4.96E-05 | 0.001594 | 0.001619927 | 5.04E-05 | 4.5 | 95.5 |
| | | 500,000 | -6.20E-06 | 0.000716 | 0.00072437 | 2.27E-05 | 3.7 | 96.3 |

| | | | | | | | | |
|---|---|---|---|---|---|---|---|---|
| | | 1,000,000 | -2.47E-05 | 0.000504 | 0.000512135 | 1.59E-05 | 4.9 | 95.1 |
| α = 0.333 | θ = 0 | 10,000 | -0.00021 | 0.004084 | 0.003931463 | 0.000129 | 5.3 | 94.7 |
| | | 20,000 | -6.96E-05 | 0.002789 | 0.002775045 | 8.82E-05 | 5.3 | 94.7 |
| | | 30,000 | -0.00014 | 0.002281 | 0.002266252 | 7.21E-05 | 5.2 | 94.8 |
| | | 40,000 | -0.00012 | 0.001977 | 0.001961668 | 6.25E-05 | 5.4 | 94.6 |
| | | 50,000 | 3.44E-05 | 0.001773 | 0.001754907 | 5.61E-05 | 5.2 | 94.8 |
| | | 60,000 | 1.09E-05 | 0.001606 | 0.001601776 | 5.08E-05 | 4.3 | 95.7 |
| | | 70,000 | 4.84E-05 | 0.001484 | 0.00148331 | 4.69E-05 | 4.5 | 95.5 |
| | | 80,000 | -2.44E-05 | 0.001378 | 0.001387465 | 4.36E-05 | 4.7 | 95.3 |
| | | 90,000 | 5.53E-06 | 0.001298 | 0.00130794 | 4.10E-05 | 5.4 | 94.6 |
| | | 100,000 | 2.72E-05 | 0.001254 | 0.001240929 | 3.97E-05 | 5.5 | 94.5 |
| | | 500,000 | -1.06E-05 | 0.000551 | 0.000554875 | 1.74E-05 | 4.4 | 95.6 |
| | | 1,000,000 | -1.60E-05 | 0.000384 | 0.000392306 | 1.21E-05 | 4.6 | 95.4 |
| α = 0.5 | θ = 0 | 10,000 | -0.0002 | 0.003609 | 0.003467276 | 0.000114 | 5.2 | 94.8 |
| | | 20,000 | -5.53E-05 | 0.00247 | 0.002447499 | 7.81E-05 | 5.1 | 94.9 |
| | | 30,000 | -0.00011 | 0.002017 | 0.001998734 | 6.38E-05 | 5 | 95 |
| | | 40,000 | -0.0001 | 0.001748 | 0.001730134 | 5.53E-05 | 5.8 | 94.2 |
| | | 50,000 | 2.96E-05 | 0.001566 | 0.001547774 | 4.95E-05 | 5.2 | 94.8 |
| | | 60,000 | 1.10E-05 | 0.00141 | 0.001412752 | 4.46E-05 | 4.5 | 95.5 |
| | | 70,000 | 4.26E-05 | 0.001307 | 0.001308257 | 4.13E-05 | 4.8 | 95.2 |
| | | 80,000 | -2.42E-05 | 0.001214 | 0.001223753 | 3.84E-05 | 4.4 | 95.6 |
| | | 90,000 | 1.59E-06 | 0.001146 | 0.001153647 | 3.62E-05 | 5 | 95 |
| | | 100,000 | 2.05E-05 | 0.001116 | 0.00109449 | 3.53E-05 | 5.3 | 94.7 |
| | | 500,000 | -1.10E-05 | 0.000486 | 0.000489386 | 1.54E-05 | 4.3 | 95.7 |
| | | 1,000,000 | -1.31E-05 | 0.000338 | 0.000346007 | 1.07E-05 | 3.7 | 96.3 |
| α = 1 | θ = 0 | 10,000 | -0.00016 | 0.002617 | 0.002509598 | 8.28E-05 | 5.2 | 94.8 |
| | | 20,000 | -3.08E-05 | 0.001801 | 0.001771684 | 5.70E-05 | 4.8 | 95.2 |
| | | 30,000 | -7.01E-05 | 0.001467 | 0.00144679 | 4.64E-05 | 4.9 | 95.1 |
| | | 40,000 | -7.36E-05 | 0.001273 | 0.001252403 | 4.03E-05 | 6 | 94 |
| | | 50,000 | 1.99E-05 | 0.001135 | 0.001120399 | 3.59E-05 | 4.9 | 95.1 |
| | | 60,000 | 9.74E-06 | 0.001011 | 0.001022722 | 3.20E-05 | 4.2 | 95.8 |
| | | 70,000 | 3.02E-05 | 0.000945 | 0.000947055 | 2.99E-05 | 5.6 | 94.4 |

| | | | | | | | | |
|---|---|---|---|---|---|---|---|---|
| | | 80,000 | -2.08E-05 | 0.000877 | 0.000885915 | 2.77E-05 | 4.2 | 95.8 |
| | | 90,000 | -3.39E-06 | 0.000833 | 0.000835211 | 2.63E-05 | 4.9 | 95.1 |
| | | 100,000 | 9.59E-06 | 0.000821 | 0.000792326 | 2.60E-05 | 5.7 | 94.3 |
| | | 500,000 | -1.00E-05 | 0.000353 | 0.000354261 | 1.12E-05 | 5.2 | 94.8 |
| | | 1,000,000 | -7.77E-06 | 0.000245 | 0.000250475 | 7.74E-06 | 3.6 | 96.4 |

#### 2.3 THETA=0.111

| Modelled mediator coefficient | Modelled interaction coefficient | Sample size | Mean observational interaction effect estimate | Standard deviation of observational estimate | Mean estimated standard error of observational estimate | Standard error of the bias (standard deviation of observational estimate/sqrt(1000)) | Power of observational estimator (%) | Coverage of observational estimator (%) |
|---|---|---|---|---|---|---|---|---|
| α = -0.333 | θ = 0.111 | 10,000 | 0.227283 | 0.006788 | 0.006461 | 0.000215 | 100 | 0 |
| | | 20,000 | 0.227317 | 0.00484 | 0.004562 | 0.000153 | 100 | 0 |
| | | 30,000 | 0.227234 | 0.003946 | 0.003726 | 0.000125 | 100 | 0 |
| | | 40,000 | 0.227303 | 0.003475 | 0.003225 | 0.00011 | 100 | 0 |
| | | 50,000 | 0.227529 | 0.003052 | 0.002885 | 9.65E-05 | 100 | 0 |
| | | 60,000 | 0.227517 | 0.002883 | 0.002633 | 9.12E-05 | 100 | 0 |
| | | 70,000 | 0.227557 | 0.002598 | 0.002438 | 8.22E-05 | 100 | 0 |
| | | 80,000 | 0.227489 | 0.002392 | 0.00228 | 7.57E-05 | 100 | 0 |
| | | 90,000 | 0.227522 | 0.00232 | 0.002149 | 7.34E-05 | 100 | 0 |
| | | 100,000 | 0.227596 | 0.002102 | 0.00204 | 6.65E-05 | 100 | 0 |
| | | 500,000 | 0.227489 | 0.000949 | 0.000912 | 3.00E-05 | 100 | 0 |
| | | 1,000,000 | 0.227459 | 0.000678 | 0.000645 | 2.14E-05 | 100 | 0 |
| α = 0 | θ = 0.111 | 10,000 | 0.24229 | 0.005457 | 0.0052 | 0.000173 | 100 | 0 |
| No mediation | | 20,000 | 0.24249 | 0.003812 | 0.00367 | 0.000121 | 100 | 0 |
| | | 30,000 | 0.242427 | 0.003171 | 0.002998 | 0.0001 | 100 | 0 |
| | | 40,000 | 0.242437 | 0.002732 | 0.002595 | 8.64E-05 | 100 | 0 |
| | | 50,000 | 0.242646 | 0.002481 | 0.002321 | 7.85E-05 | 100 | 0 |
| | | 60,000 | 0.242627 | 0.002239 | 0.002118 | 7.08E-05 | 100 | 0 |
| | | 70,000 | 0.242671 | 0.00202 | 0.001962 | 6.39E-05 | 100 | 0 |
| | | 80,000 | 0.242578 | 0.001868 | 0.001835 | 5.91E-05 | 100 | 0 |
| | | 90,000 | 0.242598 | 0.001818 | 0.00173 | 5.75E-05 | 100 | 0 |
| | | 100,000 | 0.242658 | 0.001681 | 0.001641 | 5.32E-05 | 100 | 0 |
| | | 500,000 | 0.242587 | 0.00075 | 0.000734 | 2.37E-05 | 100 | 0 |

| | | | | | | | | |
|---|---|---|---|---|---|---|---|---|
| | | 1,000,000 | 0.242576 | 0.00053 | 0.000519 | 1.67E-05 | 100 | 0 |
| α = 0.333 | θ = 0.111 | 10,000 | 0.232112 | 0.004266 | 0.003991 | 0.000135 | 100 | 0 |
| | | 20,000 | 0.232361 | 0.002975 | 0.002817 | 9.41E-05 | 100 | 0 |
| | | 30,000 | 0.232313 | 0.002487 | 0.002301 | 7.86E-05 | 100 | 0 |
| | | 40,000 | 0.232301 | 0.002131 | 0.001991 | 6.74E-05 | 100 | 0 |
| | | 50,000 | 0.23246 | 0.001947 | 0.001781 | 6.16E-05 | 100 | 0 |
| | | 60,000 | 0.232453 | 0.001711 | 0.001626 | 5.41E-05 | 100 | 0 |
| | | 70,000 | 0.232486 | 0.001563 | 0.001506 | 4.94E-05 | 100 | 0 |
| | | 80,000 | 0.232408 | 0.001442 | 0.001408 | 4.56E-05 | 100 | 0 |
| | | 90,000 | 0.232422 | 0.001402 | 0.001328 | 4.43E-05 | 100 | 0 |
| | | 100,000 | 0.23246 | 0.00133 | 0.00126 | 4.21E-05 | 100 | 0 |
| | | 500,000 | 0.232408 | 0.00058 | 0.000563 | 1.83E-05 | 100 | 0 |
| | | 1,000,000 | 0.232411 | 0.00041 | 0.000398 | 1.30E-05 | 100 | 0 |
| α = 0.5 | θ = 0.111 | 10,000 | 0.225745 | 0.003811 | 0.003528 | 0.000121 | 100 | 0 |
| | | 20,000 | 0.225997 | 0.002663 | 0.00249 | 8.42E-05 | 100 | 0 |
| | | 30,000 | 0.225954 | 0.002225 | 0.002034 | 7.04E-05 | 100 | 0 |
| | | 40,000 | 0.225938 | 0.001905 | 0.00176 | 6.02E-05 | 100 | 0 |
| | | 50,000 | 0.226076 | 0.00174 | 0.001575 | 5.50E-05 | 100 | 0 |
| | | 60,000 | 0.226075 | 0.001519 | 0.001437 | 4.80E-05 | 100 | 0 |
| | | 70,000 | 0.226103 | 0.001396 | 0.001331 | 4.41E-05 | 100 | 0 |
| | | 80,000 | 0.226032 | 0.001287 | 0.001245 | 4.07E-05 | 100 | 0 |
| | | 90,000 | 0.226044 | 0.001251 | 0.001174 | 3.96E-05 | 100 | 0 |
| | | 100,000 | 0.226076 | 0.001195 | 0.001114 | 3.78E-05 | 100 | 0 |
| | | 500,000 | 0.226029 | 0.000516 | 0.000498 | 1.63E-05 | 100 | 0 |
| | | 1,000,000 | 0.226037 | 0.000366 | 0.000352 | 1.16E-05 | 100 | 0 |
| α = 1 | θ = 0.111 | 10,000 | 0.210343 | 0.002878 | 0.002577 | 9.10E-05 | 100 | 0 |
| | | 20,000 | 0.210576 | 0.002028 | 0.001819 | 6.41E-05 | 100 | 0 |
| | | 30,000 | 0.210544 | 0.001686 | 0.001486 | 5.33E-05 | 100 | 0 |
| | | 40,000 | 0.210524 | 0.001443 | 0.001286 | 4.56E-05 | 100 | 0 |
| | | 50,000 | 0.210618 | 0.001313 | 0.001151 | 4.15E-05 | 100 | 0 |
| | | 60,000 | 0.210629 | 0.001138 | 0.00105 | 3.60E-05 | 100 | 0 |
| | | 70,000 | 0.210647 | 0.00106 | 0.000973 | 3.35E-05 | 100 | 0 |

| | | | | | | | | |
|---|---|---|---|---|---|---|---|---|
| | | 80,000 | 0.210593 | 0.000974 | 0.00091 | 3.08E-05 | 100 | 0 |
| | | 90,000 | 0.210597 | 0.00095 | 0.000858 | 3.00E-05 | 100 | 0 |
| | | 100,000 | 0.210623 | 0.000915 | 0.000814 | 2.89E-05 | 100 | 0 |
| | | 500,000 | 0.210585 | 0.000388 | 0.000364 | 1.23E-05 | 100 | 0 |
| | | 1,000,000 | 0.210599 | 0.000276 | 0.000257 | 8.74E-06 | 100 | 0 |

#### 2.4 THETA=0.167

| Modelled mediator coefficient | Modelled interaction coefficient | Sample size | Mean observational interaction effect estimate | Standard deviation of observational estimate | Mean estimated standard error of observational estimate | Standard error of the bias (standard deviation of observational estimate/sqrt(1000)) | Power of observational estimator | Coverage of observational estimator |
|---|---|---|---|---|---|---|---|---|
| α = -0.333 | θ = 0.167 | 10,000 | 0.341299 | 0.00729 | 0.006582 | 0.000231 | 100 | 0 |
| | | 20,000 | 0.341399 | 0.005277 | 0.004647 | 0.000167 | 100 | 0 |
| | | 30,000 | 0.341322 | 0.004296 | 0.003796 | 0.000136 | 100 | 0 |
| | | 40,000 | 0.341371 | 0.003755 | 0.003285 | 0.000119 | 100 | 0 |
| | | 50,000 | 0.341609 | 0.003325 | 0.002939 | 0.000105 | 100 | 0 |
| | | 60,000 | 0.34162 | 0.003126 | 0.002683 | 9.89E-05 | 100 | 0 |
| | | 70,000 | 0.341648 | 0.0028 | 0.002484 | 8.86E-05 | 100 | 0 |
| | | 80,000 | 0.341572 | 0.002563 | 0.002323 | 8.11E-05 | 100 | 0 |
| | | 90,000 | 0.341599 | 0.00253 | 0.00219 | 8.00E-05 | 100 | 0 |
| | | 100,000 | 0.341696 | 0.002295 | 0.002078 | 7.26E-05 | 100 | 0 |
| | | 500,000 | 0.341571 | 0.001025 | 0.000929 | 3.24E-05 | 100 | 0 |
| | | 1,000,000 | 0.341547 | 0.000733 | 0.000657 | 2.32E-05 | 100 | 0 |
| α = 0 | θ = 0.167 | 10,000 | 0.363898 | 0.005728 | 0.005283 | 0.000181 | 100 | 0 |
| No mediation | | 20,000 | 0.364156 | 0.004045 | 0.003729 | 0.000128 | 100 | 0 |
| | | 30,000 | 0.364104 | 0.003392 | 0.003046 | 0.000107 | 100 | 0 |
| | | 40,000 | 0.364094 | 0.002905 | 0.002636 | 9.19E-05 | 100 | 0 |
| | | 50,000 | 0.36431 | 0.002656 | 0.002358 | 8.40E-05 | 100 | 0 |
| | | 60,000 | 0.3643 | 0.002374 | 0.002152 | 7.51E-05 | 100 | 0 |
| | | 70,000 | 0.36434 | 0.002133 | 0.001993 | 6.75E-05 | 100 | 0 |
| | | 80,000 | 0.364241 | 0.001968 | 0.001864 | 6.22E-05 | 100 | 0 |
| | | 90,000 | 0.364251 | 0.001945 | 0.001757 | 6.15E-05 | 100 | 0 |
| | | 100,000 | 0.364327 | 0.001784 | 0.001668 | 5.64E-05 | 100 | 0 |
| | | 500,000 | 0.364248 | 0.000795 | 0.000746 | 2.51E-05 | 100 | 0 |

| | | | | | | | | |
|---|---|---|---|---|---|---|---|---|
| | | 1,000,000 | 0.364241 | 0.000562 | 0.000527 | 1.78E-05 | 100 | 0 |
| α = 0.333 | θ = 0.167 | 10,000 | 0.348622 | 0.004524 | 0.004064 | 0.000143 | 100 | 0 |
| | | 20,000 | 0.348926 | 0.003185 | 0.002869 | 0.000101 | 100 | 0 |
| | | 30,000 | 0.348885 | 0.002693 | 0.002343 | 8.51E-05 | 100 | 0 |
| | | 40,000 | 0.34886 | 0.002294 | 0.002028 | 7.25E-05 | 100 | 0 |
| | | 50,000 | 0.349021 | 0.002105 | 0.001814 | 6.66E-05 | 100 | 0 |
| | | 60,000 | 0.349023 | 0.001832 | 0.001656 | 5.79E-05 | 100 | 0 |
| | | 70,000 | 0.349054 | 0.001674 | 0.001534 | 5.29E-05 | 100 | 0 |
| | | 80,000 | 0.348973 | 0.001544 | 0.001434 | 4.88E-05 | 100 | 0 |
| | | 90,000 | 0.34898 | 0.001515 | 0.001352 | 4.79E-05 | 100 | 0 |
| | | 100,000 | 0.349026 | 0.001422 | 0.001283 | 4.50E-05 | 100 | 0 |
| | | 500,000 | 0.348966 | 0.00062 | 0.000574 | 1.96E-05 | 100 | 0 |
| | | 1,000,000 | 0.348974 | 0.000442 | 0.000406 | 1.40E-05 | 100 | 0 |
| α = 0.5 | θ = 0.167 | 10,000 | 0.339057 | 0.004082 | 0.003602 | 0.000129 | 100 | 0 |
| | | 20,000 | 0.339363 | 0.002879 | 0.002542 | 9.10E-05 | 100 | 0 |
| | | 30,000 | 0.339327 | 0.002433 | 0.002076 | 7.69E-05 | 100 | 0 |
| | | 40,000 | 0.339298 | 0.002071 | 0.001797 | 6.55E-05 | 100 | 0 |
| | | 50,000 | 0.339438 | 0.001897 | 0.001608 | 6.00E-05 | 100 | 0 |
| | | 60,000 | 0.339447 | 0.001644 | 0.001468 | 5.20E-05 | 100 | 0 |
| | | 70,000 | 0.339473 | 0.001513 | 0.001359 | 4.78E-05 | 100 | 0 |
| | | 80,000 | 0.3394 | 0.001395 | 0.001271 | 4.41E-05 | 100 | 0 |
| | | 90,000 | 0.339405 | 0.001366 | 0.001198 | 4.32E-05 | 100 | 0 |
| | | 100,000 | 0.339444 | 0.00129 | 0.001137 | 4.08E-05 | 100 | 0 |
| | | 500,000 | 0.339389 | 0.000557 | 0.000508 | 1.76E-05 | 100 | 0 |
| | | 1,000,000 | 0.339401 | 0.000399 | 0.000359 | 1.26E-05 | 100 | 0 |
| α = 1 | θ = 0.167 | 10,000 | 0.315911 | 0.003193 | 0.002659 | 0.000101 | 100 | 0 |
| | | 20,000 | 0.316195 | 0.002272 | 0.001877 | 7.19E-05 | 100 | 0 |
| | | 30,000 | 0.316167 | 0.001904 | 0.001533 | 6.02E-05 | 100 | 0 |
| | | 40,000 | 0.316138 | 0.001619 | 0.001327 | 5.12E-05 | 100 | 0 |
| | | 50,000 | 0.316234 | 0.001477 | 0.001187 | 4.67E-05 | 100 | 0 |
| | | 60,000 | 0.316255 | 0.001276 | 0.001084 | 4.03E-05 | 100 | 0 |
| | | 70,000 | 0.316271 | 0.001194 | 0.001004 | 3.78E-05 | 100 | 0 |

| | | | | | | | | |
|---|---|---|---|---|---|---|---|---|
| | | 80,000 | 0.316216 | 0.0011 | 0.000939 | 3.48E-05 | 100 | 0 |
| | | 90,000 | 0.316213 | 0.001075 | 0.000885 | 3.40E-05 | 100 | 0 |
| | | 100,000 | 0.316246 | 0.001023 | 0.00084 | 3.24E-05 | 100 | 0 |
| | | 500,000 | 0.316199 | 0.000434 | 0.000375 | 1.37E-05 | 100 | 0 |
| | | 1,000,000 | 0.316218 | 0.000313 | 0.000265 | 9.88E-06 | 100 | 0 |

#### 2.5 THETA=0.333

| Modelled mediator coefficient | Modelled interaction coefficient | Sample size | Mean observational interaction effect estimate | Standard deviation of observational estimate | Mean estimated standard error of observational estimate | Standard error of the bias (standard deviation of observational estimate/sqrt(1000)) | Power of observational estimator | Coverage of observational estimator |
|---|---|---|---|---|---|---|---|---|
| α = -0.333 | θ = 0.333 | 10,000 | 0.682663 | 0.009651 | 0.007199 | 0.000305 | 100 | 0 |
| | | 20,000 | 0.682963 | 0.007144 | 0.005085 | 0.000226 | 100 | 0 |
| | | 30,000 | 0.682903 | 0.005803 | 0.004153 | 0.000184 | 100 | 0 |
| | | 40,000 | 0.682894 | 0.004991 | 0.003594 | 0.000158 | 100 | 0 |
| | | 50,000 | 0.683167 | 0.004502 | 0.003215 | 0.000142 | 100 | 0 |
| | | 60,000 | 0.683243 | 0.00417 | 0.002935 | 0.000132 | 100 | 0 |
| | | 70,000 | 0.683235 | 0.003737 | 0.002718 | 0.000118 | 100 | 0 |
| | | 80,000 | 0.683138 | 0.003384 | 0.002542 | 0.000107 | 100 | 0 |
| | | 90,000 | 0.683147 | 0.00341 | 0.002396 | 0.000108 | 100 | 0 |
| | | 100,000 | 0.68331 | 0.00312 | 0.002274 | 9.87E-05 | 100 | 0 |
| | | 500,000 | 0.683132 | 0.001364 | 0.001017 | 4.31E-05 | 100 | 0 |
| | | 1,000,000 | 0.683128 | 0.000981 | 0.000719 | 3.10E-05 | 100 | 0 |
| α = 0 | θ = 0.333 | 10,000 | 0.727997 | 0.007092 | 0.005711 | 0.000224 | 100 | 0 |
| No mediation | | 20,000 | 0.728426 | 0.005103 | 0.004032 | 0.000161 | 100 | 0 |
| | | 30,000 | 0.728408 | 0.004353 | 0.003293 | 0.000138 | 100 | 0 |
| | | 40,000 | 0.72834 | 0.003681 | 0.00285 | 0.000116 | 100 | 0 |
| | | 50,000 | 0.728575 | 0.0034 | 0.00255 | 0.000108 | 100 | 0 |
| | | 60,000 | 0.728593 | 0.002998 | 0.002327 | 9.48E-05 | 100 | 0 |
| | | 70,000 | 0.728619 | 0.002699 | 0.002155 | 8.54E-05 | 100 | 0 |
| | | 80,000 | 0.7285 | 0.002481 | 0.002016 | 7.85E-05 | 100 | 0 |
| | | 90,000 | 0.728481 | 0.002501 | 0.0019 | 7.91E-05 | 100 | 0 |
| | | 100,000 | 0.728604 | 0.002255 | 0.001803 | 7.13E-05 | 100 | 0 |
| | | 500,000 | 0.728502 | 0.001007 | 0.000806 | 3.19E-05 | 100 | 0 |

| | | | | | | | | |
|---|---|---|---|---|---|---|---|---|
| | | 1,000,000 | 0.728507 | 0.000713 | 0.00057 | 2.25E-05 | 100 | 0 |
| α = 0.333 | θ = 0.333 | 10,000 | 0.697454 | 0.005767 | 0.004439 | 0.000182 | 100 | 0 |
| | | 20,000 | 0.697921 | 0.004128 | 0.003134 | 0.000131 | 100 | 0 |
| | | 30,000 | 0.697906 | 0.003566 | 0.00256 | 0.000113 | 100 | 0 |
| | | 40,000 | 0.697838 | 0.003005 | 0.002216 | 9.50E-05 | 100 | 0 |
| | | 50,000 | 0.698008 | 0.002758 | 0.001982 | 8.72E-05 | 100 | 0 |
| | | 60,000 | 0.698036 | 0.002381 | 0.001809 | 7.53E-05 | 100 | 0 |
| | | 70,000 | 0.69806 | 0.002198 | 0.001675 | 6.95E-05 | 100 | 0 |
| | | 80,000 | 0.69797 | 0.002037 | 0.001567 | 6.44E-05 | 100 | 0 |
| | | 90,000 | 0.697954 | 0.002012 | 0.001477 | 6.36E-05 | 100 | 0 |
| | | 100,000 | 0.698024 | 0.00184 | 0.001401 | 5.82E-05 | 100 | 0 |
| | | 500,000 | 0.697942 | 0.000809 | 0.000627 | 2.56E-05 | 100 | 0 |
| | | 1,000,000 | 0.697964 | 0.000585 | 0.000443 | 1.85E-05 | 100 | 0 |
| α = 0.5 | θ = 0.333 | 10,000 | 0.678314 | 0.005351 | 0.003978 | 0.000169 | 100 | 0 |
| | | 20,000 | 0.678781 | 0.003833 | 0.002808 | 0.000121 | 100 | 0 |
| | | 30,000 | 0.678766 | 0.003304 | 0.002293 | 0.000104 | 100 | 0 |
| | | 40,000 | 0.678702 | 0.002782 | 0.001985 | 8.80E-05 | 100 | 0 |
| | | 50,000 | 0.678847 | 0.002543 | 0.001776 | 8.04E-05 | 100 | 0 |
| | | 60,000 | 0.678883 | 0.002194 | 0.001621 | 6.94E-05 | 100 | 0 |
| | | 70,000 | 0.678904 | 0.002046 | 0.001501 | 6.47E-05 | 100 | 0 |
| | | 80,000 | 0.678824 | 0.0019 | 0.001404 | 6.01E-05 | 100 | 0 |
| | | 90,000 | 0.678808 | 0.001862 | 0.001324 | 5.89E-05 | 100 | 0 |
| | | 100,000 | 0.678867 | 0.001714 | 0.001256 | 5.42E-05 | 100 | 0 |
| | | 500,000 | 0.678788 | 0.000747 | 0.000561 | 2.36E-05 | 100 | 0 |
| | | 1,000,000 | 0.678816 | 0.000544 | 0.000397 | 1.72E-05 | 100 | 0 |
| α = 1 | θ = 0.333 | 10,000 | 0.631984 | 0.004546 | 0.003065 | 0.000144 | 100 | 0 |
| | | 20,000 | 0.632421 | 0.003283 | 0.002164 | 0.000104 | 100 | 0 |
| | | 30,000 | 0.632404 | 0.002781 | 0.001767 | 8.80E-05 | 100 | 0 |
| | | 40,000 | 0.63235 | 0.00234 | 0.00153 | 7.40E-05 | 100 | 0 |
| | | 50,000 | 0.632448 | 0.002124 | 0.001368 | 6.72E-05 | 100 | 0 |
| | | 60,000 | 0.632501 | 0.001843 | 0.001249 | 5.83E-05 | 100 | 0 |
| | | 70,000 | 0.632512 | 0.001752 | 0.001157 | 5.54E-05 | 100 | 0 |

| | | | | | | | | |
|---|---|---|---|---|---|---|---|---|
| | | 80,000 | 0.632453 | 0.001632 | 0.001082 | 5.16E-05 | 100 | 0 |
| | | 90,000 | 0.632429 | 0.001582 | 0.00102 | 5.00E-05 | 100 | 0 |
| | | 100,000 | 0.632482 | 0.001476 | 0.000968 | 4.67E-05 | 100 | 0 |
| | | 500,000 | 0.632409 | 0.000631 | 0.000433 | 1.99E-05 | 100 | 0 |
| | | 1,000,000 | 0.632445 | 0.000463 | 0.000306 | 1.46E-05 | 100 | 0 |

### 3 2SLS INSTRUMENT DOES NOT ASSUME MEDIATION, Z=(Z1,Z2,Z1Z2)

#### 3.1 THETA=-0.111

| Modelled mediator coefficient | Modelled interaction coefficient | Sample size | Mean 2sls interaction effect estimate | Standard deviation of 2sls estimate | Mean estimated standard error of 2sls estimate | Standard error of the bias (standard deviation of 2sls estimate/sqrt(1000)) | Power of 2sls estimator (%) | Coverage of 2sls estimator (%) |
|---|---|---|---|---|---|---|---|---|
| α = -0.333 | θ = -0.111 | 10,000 | -0.21775 | 6.214008 | 43.15983 | 0.196504 | 1.2 | 100 |
| | | 20,000 | -0.069 | 1.723318 | 6.548343 | 0.054496 | 1.7 | 99.7 |
| | | 30,000 | 0.016721 | 3.693011 | 32.8285 | 0.116783 | 3.2 | 99.1 |
| | | 40,000 | -0.10449 | 0.405008 | 0.441858 | 0.012807 | 5.9 | 97.7 |
| | | 50,000 | 0.03879 | 5.227974 | 33.43623 | 0.165323 | 7.2 | 97.9 |
| | | 60,000 | -0.10669 | 0.274416 | 0.282685 | 0.008678 | 6.7 | 98.2 |
| | | 70,000 | -0.10058 | 0.250272 | 0.259073 | 0.007914 | 8 | 97.8 |
| | | 80,000 | -0.10868 | 0.243907 | 0.236284 | 0.007713 | 7.5 | 97.3 |
| | | 90,000 | -0.08746 | 0.227107 | 0.220484 | 0.007182 | 7.1 | 98 |
| | | 100,000 | -0.1003 | 0.202746 | 0.202602 | 0.006411 | 9.8 | 98.6 |
| | | 500,000 | -0.10677 | 0.084521 | 0.082203 | 0.002673 | 28.1 | 94.2 |
| | | 1,000,000 | -0.11088 | 0.056678 | 0.057986 | 0.001792 | 48.7 | 95.3 |
| α = 0 | θ = -0.111 | 10,000 | 0.246957 | 6.568474 | 62.61536 | 0.207713 | 1.5 | 99.6 |
| No mediation | | 20,000 | 1.096622 | 39.55999 | 1464.155 | 1.250997 | 2.1 | 99.6 |
| | | 30,000 | -0.02826 | 2.979395 | 42.48063 | 0.094217 | 3.7 | 98.9 |
| | | 40,000 | -0.09496 | 1.219632 | 2.056836 | 0.038568 | 6.3 | 98.4 |
| | | 50,000 | -0.11402 | 0.51638 | 0.66536 | 0.016329 | 7.4 | 97.5 |
| | | 60,000 | -0.08589 | 0.505889 | 0.614479 | 0.015998 | 6.7 | 98.5 |
| | | 70,000 | -0.08715 | 0.355384 | 0.370698 | 0.011238 | 8.4 | 98.6 |
| | | 80,000 | -0.10475 | 1.033444 | 3.087159 | 0.03268 | 8.3 | 97.5 |
| | | 90,000 | -0.06191 | 0.441419 | 0.376284 | 0.013959 | 7.7 | 98.5 |
| | | 100,000 | -0.09253 | 0.220696 | 0.222076 | 0.006979 | 10.1 | 99 |
| | | 500,000 | -0.10599 | 0.086511 | 0.083838 | 0.002736 | 27.8 | 94.3 |
| | | 1,000,000 | -0.1103 | 0.057625 | 0.058942 | 0.001822 | 47.7 | 95.7 |

| | | | | | | | | |
|---|---|---|---|---|---|---|---|---|
| α = 0.333 | θ = -0.111 | 10,000 | -0.36033 | 6.508636 | 90.21198 | 0.205821 | 1.9 | 99.6 |
| | | 20,000 | 0.515166 | 11.8878 | 251.7985 | 0.375925 | 3 | 99.5 |
| | | 30,000 | -0.32242 | 4.218115 | 172.7044 | 0.133388 | 4.4 | 99.1 |
| | | 40,000 | -0.04793 | 1.584554 | 7.390424 | 0.050108 | 6 | 98.8 |
| | | 50,000 | -0.57867 | 17.22369 | 381.2039 | 0.544661 | 7.6 | 97.7 |
| | | 60,000 | -0.16338 | 1.162284 | 2.872041 | 0.036755 | 6.9 | 98.5 |
| | | 70,000 | 1.031644 | 35.24122 | 1722.213 | 1.114425 | 8.4 | 99.1 |
| | | 80,000 | 0.053764 | 3.430512 | 12.64235 | 0.108482 | 8.6 | 98.4 |
| | | 90,000 | -0.18281 | 4.12814 | 24.30278 | 0.130543 | 7.5 | 98.6 |
| | | 100,000 | -0.08139 | 0.897594 | 2.551107 | 0.028384 | 10.1 | 99.3 |
| | | 500,000 | -0.10473 | 0.089978 | 0.086718 | 0.002845 | 27.6 | 95 |
| | | 1,000,000 | -0.10955 | 0.058984 | 0.060322 | 0.001865 | 47.1 | 96.4 |
| α = 0.5 | θ = -0.111 | 10,000 | -0.32521 | 5.340703 | 57.00212 | 0.168888 | 2.2 | 99.7 |
| | | 20,000 | -0.17623 | 6.479462 | 85.99288 | 0.204899 | 3.1 | 99.6 |
| | | 30,000 | -0.12533 | 3.023454 | 42.98291 | 0.09561 | 4.8 | 99.1 |
| | | 40,000 | -0.15743 | 2.543494 | 25.46778 | 0.080432 | 6.4 | 98.8 |
| | | 50,000 | -0.12348 | 2.230048 | 10.55006 | 0.07052 | 7.9 | 97.8 |
| | | 60,000 | -0.32028 | 7.239807 | 272.1581 | 0.228943 | 7 | 98.8 |
| | | 70,000 | -0.20027 | 2.297505 | 11.57199 | 0.072653 | 8.5 | 99.3 |
| | | 80,000 | -0.22864 | 2.213789 | 16.7742 | 0.070006 | 9.1 | 98.5 |
| | | 90,000 | -0.04359 | 1.285722 | 3.771445 | 0.040658 | 7.7 | 98.7 |
| | | 100,000 | -0.06435 | 0.473639 | 0.781673 | 0.014978 | 10.3 | 99.3 |
| | | 500,000 | -0.10388 | 0.092499 | 0.088789 | 0.002925 | 27.5 | 95.8 |
| | | 1,000,000 | -0.10911 | 0.059847 | 0.061197 | 0.001893 | 46.3 | 96.8 |
| α = 1 | θ = -0.111 | 10,000 | -0.29284 | 5.188695 | 88.12826 | 0.164081 | 3.2 | 99.7 |
| | | 20,000 | -0.46863 | 9.160373 | 294.736 | 0.289676 | 4.1 | 99.6 |
| | | 30,000 | -0.22865 | 2.631471 | 47.38844 | 0.083214 | 5.6 | 99.1 |
| | | 40,000 | -1.63516 | 69.79232 | 30492.19 | 2.207027 | 7.1 | 99.1 |
| | | 50,000 | -0.13721 | 8.255129 | 166.3684 | 0.26105 | 8.6 | 98 |
| | | 60,000 | -0.20744 | 2.236071 | 11.54497 | 0.070711 | 8.8 | 99.1 |
| | | 70,000 | -0.29327 | 4.868044 | 42.68373 | 0.153941 | 8.2 | 99.3 |
| | | 80,000 | 0.01058 | 22.58778 | 4609.322 | 0.714288 | 9.5 | 98.8 |

| | | | | | | | | |
|---|---|---|---|---|---|---|---|---|
| | | 90,000 | -0.25879 | 4.325907 | 55.24257 | 0.136797 | 8.8 | 98.6 |
| | | 100,000 | -0.26527 | 4.998768 | 28.97802 | 0.158075 | 10.5 | 99 |
| | | 500,000 | -0.09994 | 0.106611 | 0.099948 | 0.003371 | 27.9 | 96.9 |
| | | 1,000,000 | -0.10742 | 0.063393 | 0.064744 | 0.002005 | 46.2 | 97.7 |

#### 3.2 THETA=0

| Modelled mediator coefficient | Modelled interaction coefficient | Sample size | Mean 2sls interaction effect estimate | Standard deviation of 2sls estimate | Mean estimated standard error of 2sls estimate | Standard error of the bias (standard deviation of 2sls estimate/sqrt(1000)) | Type I error of 2sls estimator | Coverage of 2sls estimator |
|---|---|---|---|---|---|---|---|---|
| α = -0.333 | θ = 0 | 10,000 | -0.06625 | 6.257886 | 45.53554 | 0.197892 | 0 | 100 |
| | | 20,000 | -0.05187 | 1.652465 | 5.91879 | 0.052256 | 0.2 | 99.8 |
| | | 30,000 | 0.031233 | 4.999218 | 40.59871 | 0.158089 | 0.4 | 99.6 |
| | | 40,000 | -0.019 | 0.425931 | 0.452861 | 0.013469 | 1 | 99 |
| | | 50,000 | 0.201257 | 7.493741 | 48.00768 | 0.236973 | 1.1 | 98.9 |
| | | 60,000 | -0.0058 | 0.265096 | 0.275365 | 0.008383 | 1.2 | 98.8 |
| | | 70,000 | -0.00454 | 0.238511 | 0.250518 | 0.007542 | 2 | 98 |
| | | 80,000 | -0.00648 | 0.235611 | 0.230215 | 0.007451 | 2.6 | 97.4 |
| | | 90,000 | 0.011816 | 0.214749 | 0.213654 | 0.006791 | 2.1 | 97.9 |
| | | 100,000 | 0.002848 | 0.199567 | 0.198006 | 0.006311 | 1.6 | 98.4 |
| | | 500,000 | 0.003615 | 0.083695 | 0.080411 | 0.002647 | 6.3 | 93.7 |
| | | 1,000,000 | -0.00108 | 0.055738 | 0.056762 | 0.001763 | 4.7 | 95.3 |
| α = 0 | θ = 0 | 10,000 | 0.302075 | 6.575332 | 61.31404 | 0.20793 | 0 | 100 |
| No mediation | | 20,000 | 0.521476 | 36.51752 | 1523.227 | 1.154785 | 0.2 | 99.8 |
| | | 30,000 | -0.16634 | 4.580503 | 94.23084 | 0.144848 | 0.2 | 99.8 |
| | | 40,000 | -0.01389 | 1.045202 | 2.085443 | 0.033052 | 0.7 | 99.3 |
| | | 50,000 | -0.03516 | 0.585904 | 0.805202 | 0.018528 | 0.7 | 99.3 |
| | | 60,000 | -0.00411 | 0.452345 | 0.553792 | 0.014304 | 0.4 | 99.6 |
| | | 70,000 | -0.00557 | 0.318206 | 0.331378 | 0.010063 | 0.8 | 99.2 |
| | | 80,000 | 0.003372 | 0.337123 | 0.651364 | 0.010661 | 1.3 | 98.7 |
| | | 90,000 | 0.020326 | 0.297559 | 0.291984 | 0.00941 | 1.1 | 98.9 |
| | | 100,000 | 0.003015 | 0.215195 | 0.214225 | 0.006805 | 1 | 99 |
| | | 500,000 | 0.003522 | 0.085028 | 0.081221 | 0.002689 | 5.7 | 94.3 |
| | | 1,000,000 | -0.00113 | 0.056278 | 0.057176 | 0.00178 | 4.7 | 95.3 |

| | | | | | | | | |
|---|---|---|---|---|---|---|---|---|
| α = 0.333 | θ = 0 | 10,000 | -0.16879 | 5.890676 | 130.6215 | 0.18628 | 0 | 100 |
| | | 20,000 | 0.551993 | 9.280967 | 200.6147 | 0.29349 | 0.1 | 99.9 |
| | | 30,000 | -0.22368 | 8.6254 | 455.9818 | 0.272759 | 0.1 | 99.9 |
| | | 40,000 | -0.00494 | 0.965853 | 3.107193 | 0.030543 | 0.3 | 99.7 |
| | | 50,000 | -0.43591 | 14.5223 | 318.9475 | 0.459236 | 0.2 | 99.8 |
| | | 60,000 | -0.04154 | 0.826501 | 1.953673 | 0.026136 | 0.1 | 99.9 |
| | | 70,000 | 0.777269 | 24.56081 | 1200.354 | 0.776681 | 0.4 | 99.6 |
| | | 80,000 | 0.091533 | 2.358262 | 8.581851 | 0.074575 | 0.3 | 99.7 |
| | | 90,000 | -0.01449 | 3.532681 | 24.83627 | 0.111713 | 0.5 | 99.5 |
| | | 100,000 | 0.005468 | 0.766312 | 1.995738 | 0.024233 | 0.7 | 99.3 |
| | | 500,000 | 0.003446 | 0.087553 | 0.082996 | 0.002769 | 4.7 | 95.3 |
| | | 1,000,000 | -0.0012 | 0.057109 | 0.057865 | 0.001806 | 4.4 | 95.6 |
| α = 0.5 | θ = 0 | 10,000 | -0.18757 | 5.736502 | 58.64873 | 0.181404 | 0 | 100 |
| | | 20,000 | -0.13307 | 5.868711 | 82.26659 | 0.185585 | 0.1 | 99.9 |
| | | 30,000 | -0.10436 | 4.484523 | 68.65502 | 0.141813 | 0.1 | 99.9 |
| | | 40,000 | -0.19128 | 2.822528 | 21.53514 | 0.089256 | 0.3 | 99.7 |
| | | 50,000 | 0.020761 | 1.54377 | 6.689685 | 0.048818 | 0.1 | 99.9 |
| | | 60,000 | -0.21769 | 4.632296 | 198.8609 | 0.146486 | 0.1 | 99.9 |
| | | 70,000 | -0.05389 | 1.171219 | 5.266748 | 0.037037 | 0.4 | 99.6 |
| | | 80,000 | -0.0534 | 1.451717 | 7.088886 | 0.045907 | 0.2 | 99.8 |
| | | 90,000 | 0.014493 | 1.110803 | 3.291321 | 0.035127 | 0.5 | 99.5 |
| | | 100,000 | 0.018968 | 0.434499 | 0.628817 | 0.01374 | 0.5 | 99.5 |
| | | 500,000 | 0.003409 | 0.089473 | 0.084375 | 0.002829 | 4.3 | 95.7 |
| | | 1,000,000 | -0.00123 | 0.057656 | 0.058328 | 0.001823 | 4.1 | 95.9 |
| α = 1 | θ = 0 | 10,000 | -0.13832 | 5.399892 | 91.33763 | 0.17076 | 0 | 100 |
| | | 20,000 | -0.45335 | 13.82701 | 444.3984 | 0.437248 | 0.1 | 99.9 |
| | | 30,000 | 0.014092 | 1.989054 | 29.79733 | 0.062899 | 0.1 | 99.9 |
| | | 40,000 | -0.61841 | 40.9668 | 17496.97 | 1.295484 | 0.1 | 99.9 |
| | | 50,000 | 0.105713 | 8.622337 | 173.8918 | 0.272662 | 0 | 100 |
| | | 60,000 | -0.07933 | 2.378191 | 11.51237 | 0.075205 | 0 | 100 |
| | | 70,000 | -0.14894 | 5.168856 | 43.18671 | 0.163454 | 0 | 100 |
| | | 80,000 | -1.55432 | 33.823 | 5414.013 | 1.069577 | 0.1 | 99.9 |

| | | | | | | | | |
|---|---|---|---|---|---|---|---|---|
| | | 90,000 | -0.11091 | 2.640426 | 37.04351 | 0.083498 | 0.2 | 99.8 |
| | | 100,000 | -0.14681 | 4.102587 | 25.07565 | 0.129735 | 0.3 | 99.7 |
| | | 500,000 | 0.00305 | 0.102249 | 0.092846 | 0.003233 | 2.6 | 97.4 |
| | | 1,000,000 | -0.00137 | 0.059998 | 0.060316 | 0.001897 | 3.4 | 96.6 |

#### 3.3 THETA=0.111

| Modelled mediator coefficient | Modelled interaction coefficient | Sample size | Mean 2sls interaction effect estimate | Standard deviation of 2sls estimate | Mean estimated standard error of 2sls estimate | Standard error of the bias (standard deviation of 2sls estimate/sqrt(1000)) | Power of 2sls estimator | Coverage of 2sls estimator |
|---|---|---|---|---|---|---|---|---|
| α = -0.333 | θ = 0.111 | 10,000 | 0.085238 | 6.562822 | 49.52134 | 0.207535 | 1.7 | 100 |
| | | 20,000 | -0.03474 | 2.442886 | 11.2895 | 0.077251 | 2.8 | 99.4 |
| | | 30,000 | 0.045745 | 6.9286 | 48.51535 | 0.219102 | 3.6 | 99.2 |
| | | 40,000 | 0.066495 | 0.506448 | 0.519387 | 0.016015 | 4.5 | 98.2 |
| | | 50,000 | 0.363723 | 9.771656 | 62.66634 | 0.309007 | 4.6 | 98.8 |
| | | 60,000 | 0.095085 | 0.27349 | 0.282726 | 0.008649 | 5.7 | 98.3 |
| | | 70,000 | 0.091487 | 0.244835 | 0.257173 | 0.007742 | 5.3 | 97.8 |
| | | 80,000 | 0.095728 | 0.241232 | 0.235719 | 0.007628 | 8.5 | 96.5 |
| | | 90,000 | 0.111088 | 0.215324 | 0.217744 | 0.006809 | 9.1 | 97.6 |
| | | 100,000 | 0.105991 | 0.205774 | 0.202752 | 0.006507 | 10 | 97.7 |
| | | 500,000 | 0.114003 | 0.086158 | 0.0821 | 0.002725 | 30.7 | 94.1 |
| | | 1,000,000 | 0.108718 | 0.057131 | 0.058006 | 0.001807 | 46.1 | 95.3 |
| α = 0 | θ = 0.111 | 10,000 | 0.357192 | 7.15916 | 64.34084 | 0.226393 | 1.6 | 99.9 |
| No mediation | | 20,000 | -0.05367 | 37.16733 | 1600.504 | 1.175334 | 3.6 | 99.1 |
| | | 30,000 | -0.30441 | 9.394095 | 213.2775 | 0.297067 | 3.6 | 99.4 |
| | | 40,000 | 0.067182 | 1.255177 | 2.625237 | 0.039692 | 4.4 | 98.6 |
| | | 50,000 | 0.0437 | 0.757077 | 1.02221 | 0.023941 | 5.5 | 99.5 |
| | | 60,000 | 0.077682 | 0.634207 | 0.803016 | 0.020055 | 6 | 98.3 |
| | | 70,000 | 0.076 | 0.34817 | 0.338319 | 0.01101 | 5.7 | 98.2 |
| | | 80,000 | 0.111494 | 1.171376 | 3.544879 | 0.037042 | 9.5 | 97.2 |
| | | 90,000 | 0.102563 | 0.275337 | 0.287034 | 0.008707 | 9.3 | 98.3 |
| | | 100,000 | 0.098559 | 0.229209 | 0.223779 | 0.007248 | 10.1 | 98.1 |
| | | 500,000 | 0.113035 | 0.08841 | 0.083676 | 0.002796 | 30.5 | 94.5 |
| | | 1,000,000 | 0.108036 | 0.058265 | 0.058983 | 0.001842 | 46.4 | 95.5 |

| | | | | | | | | |
|---|---|---|---|---|---|---|---|---|
| α = 0.333 | θ = 0.111 | 10,000 | 0.02275 | 6.381498 | 181.659 | 0.201801 | 2.1 | 99.8 |
| | | 20,000 | 0.58882 | 7.255711 | 150.1406 | 0.229446 | 3.7 | 99.3 |
| | | 30,000 | -0.12494 | 14.10819 | 754.1116 | 0.44614 | 3.7 | 99.6 |
| | | 40,000 | 0.038054 | 1.322966 | 6.185259 | 0.041836 | 4.7 | 98.6 |
| | | 50,000 | -0.29315 | 11.95199 | 260.83 | 0.377955 | 5.2 | 99.6 |
| | | 60,000 | 0.080297 | 1.026853 | 2.613358 | 0.032472 | 7.1 | 99 |
| | | 70,000 | 0.522894 | 13.89466 | 678.7721 | 0.439388 | 5.8 | 98.9 |
| | | 80,000 | 0.129303 | 1.401817 | 4.726981 | 0.044329 | 10.4 | 97.9 |
| | | 90,000 | 0.153821 | 4.041784 | 25.48676 | 0.127812 | 9.5 | 98.9 |
| | | 100,000 | 0.092321 | 1.437649 | 4.349528 | 0.045462 | 10.4 | 98.4 |
| | | 500,000 | 0.111627 | 0.092241 | 0.086503 | 0.002917 | 30.7 | 95 |
| | | 1,000,000 | 0.107164 | 0.059868 | 0.060399 | 0.001893 | 45.6 | 95.6 |
| α = 0.5 | θ = 0.111 | 10,000 | -0.04992 | 6.317666 | 62.13505 | 0.199782 | 2.5 | 99.8 |
| | | 20,000 | -0.0899 | 5.804687 | 79.92608 | 0.18356 | 3.7 | 99.3 |
| | | 30,000 | -0.08339 | 6.90051 | 102.1386 | 0.218213 | 4 | 99.6 |
| | | 40,000 | -0.22512 | 4.708213 | 50.31435 | 0.148887 | 5.2 | 98.8 |
| | | 50,000 | 0.165003 | 1.792945 | 7.415969 | 0.056698 | 5 | 99.7 |
| | | 60,000 | -0.1151 | 7.856869 | 258.1645 | 0.248456 | 7.5 | 99 |
| | | 70,000 | 0.092482 | 1.30247 | 6.030621 | 0.041188 | 5.8 | 99.2 |
| | | 80,000 | 0.121836 | 2.911936 | 21.92397 | 0.092083 | 10.3 | 98.2 |
| | | 90,000 | 0.072576 | 1.545835 | 5.012182 | 0.048884 | 9.9 | 98.8 |
| | | 100,000 | 0.102284 | 0.748443 | 1.187633 | 0.023668 | 10.4 | 98.6 |
| | | 500,000 | 0.110696 | 0.095083 | 0.08857 | 0.003007 | 31.2 | 95.5 |
| | | 1,000,000 | 0.106644 | 0.060886 | 0.061301 | 0.001925 | 44.8 | 95.8 |
| α = 1 | θ = 0.111 | 10,000 | 0.016191 | 5.706895 | 95.16416 | 0.180468 | 3.2 | 99.7 |
| | | 20,000 | -0.43807 | 18.57751 | 594.795 | 0.587472 | 4.6 | 99.5 |
| | | 30,000 | 0.256829 | 4.11468 | 85.66557 | 0.130118 | 5.7 | 99.6 |
| | | 40,000 | 0.398352 | 16.43501 | 4503.089 | 0.519721 | 5.8 | 99.1 |
| | | 50,000 | 0.348632 | 9.45311 | 190.2508 | 0.298934 | 6.3 | 99.7 |
| | | 60,000 | 0.048776 | 2.693678 | 12.41903 | 0.085182 | 8.3 | 99.5 |
| | | 70,000 | -0.00461 | 5.685566 | 47.0415 | 0.179793 | 6.9 | 99.5 |
| | | 80,000 | -3.11921 | 70.25019 | 13769.61 | 2.221506 | 10 | 98.4 |

| | | | | | | | | |
|---|---|---|---|---|---|---|---|---|
| | | 90,000 | 0.036976 | 2.762198 | 22.36578 | 0.087348 | 9.9 | 99.3 |
| | | 100,000 | -0.02836 | 3.518813 | 22.27361 | 0.111275 | 11.2 | 98.6 |
| | | 500,000 | 0.10604 | 0.115306 | 0.10096 | 0.003646 | 31.7 | 96.4 |
| | | 1,000,000 | 0.104675 | 0.065148 | 0.064984 | 0.00206 | 43.3 | 96.6 |

#### 3.4 THETA=0.167

| Modelled mediator coefficient | Modelled interaction coefficient | Sample size | Mean 2sls interaction effect estimate | Standard deviation of 2sls estimate | Mean estimated standard error of 2sls estimate | Standard error of the bias (standard deviation of 2sls estimate/sqrt(1000)) | Power of 2sls estimator | Coverage of 2sls estimator |
|---|---|---|---|---|---|---|---|---|
| α = -0.333 | θ = 0.167 | 10,000 | 0.161211 | 6.804202 | 51.68335 | 0.215168 | 3.1 | 99.6 |
| | | 20,000 | -0.02615 | 2.983269 | 14.40362 | 0.094339 | 6.6 | 99 |
| | | 30,000 | 0.053023 | 7.996032 | 52.58625 | 0.252857 | 6.9 | 98.8 |
| | | 40,000 | 0.109369 | 0.562013 | 0.561665 | 0.017772 | 9.3 | 98 |
| | | 50,000 | 0.4452 | 10.91622 | 70.02417 | 0.345201 | 9.9 | 98.7 |
| | | 60,000 | 0.14568 | 0.284006 | 0.291775 | 0.008981 | 11.5 | 98.1 |
| | | 70,000 | 0.139648 | 0.254543 | 0.266112 | 0.008049 | 10.9 | 97.7 |
| | | 80,000 | 0.146983 | 0.249086 | 0.242707 | 0.007877 | 16.5 | 96.1 |
| | | 90,000 | 0.160873 | 0.220538 | 0.223801 | 0.006974 | 15.2 | 97.8 |
| | | 100,000 | 0.157717 | 0.212212 | 0.208499 | 0.006711 | 17.9 | 97.5 |
| | | 500,000 | 0.169364 | 0.088559 | 0.084209 | 0.0028 | 52.8 | 94.5 |
| | | 1,000,000 | 0.163782 | 0.058667 | 0.059522 | 0.001855 | 78.5 | 95.3 |
| α = 0 | θ = 0.167 | 10,000 | 0.384834 | 7.632824 | 66.67638 | 0.241371 | 4.5 | 99.6 |
| No mediation | | 20,000 | -0.34211 | 38.86641 | 1659.691 | 1.229064 | 6.9 | 98.6 |
| | | 30,000 | -0.37366 | 11.94794 | 273.6259 | 0.377827 | 7.9 | 99.2 |
| | | 40,000 | 0.107839 | 1.46571 | 2.949423 | 0.04635 | 10.7 | 98.1 |
| | | 50,000 | 0.083248 | 0.8638 | 1.166297 | 0.027316 | 11.3 | 99.3 |
| | | 60,000 | 0.118699 | 0.771383 | 0.946173 | 0.024393 | 13.4 | 97.9 |
| | | 70,000 | 0.116908 | 0.384944 | 0.369013 | 0.012173 | 11.9 | 98.1 |
| | | 80,000 | 0.165717 | 1.684155 | 5.070107 | 0.053258 | 17.4 | 96.5 |
| | | 90,000 | 0.143806 | 0.323509 | 0.328469 | 0.01023 | 17.4 | 98 |
| | | 100,000 | 0.146475 | 0.242708 | 0.23457 | 0.007675 | 19.4 | 97.4 |
| | | 500,000 | 0.167956 | 0.091799 | 0.086716 | 0.002903 | 51.9 | 94.4 |
| | | 1,000,000 | 0.162784 | 0.060435 | 0.061158 | 0.001911 | 75.9 | 95.6 |

| | | | | | | | | |
|---|---|---|---|---|---|---|---|---|
| α = 0.333 | θ = 0.167 | 10,000 | 0.118808 | 6.992638 | 210.8103 | 0.221127 | 5.6 | 99.2 |
| | | 20,000 | 0.607289 | 6.63452 | 125.3148 | 0.209802 | 8.1 | 98.6 |
| | | 30,000 | -0.07542 | 16.94599 | 904.9135 | 0.535879 | 10 | 99.2 |
| | | 40,000 | 0.059614 | 1.749855 | 8.837436 | 0.055335 | 12.3 | 98 |
| | | 50,000 | -0.22155 | 10.74209 | 236.7258 | 0.339695 | 11.9 | 99.2 |
| | | 60,000 | 0.141399 | 1.27745 | 3.261332 | 0.040397 | 14.5 | 98.4 |
| | | 70,000 | 0.395325 | 8.56771 | 417.2397 | 0.270935 | 12 | 98.4 |
| | | 80,000 | 0.148244 | 1.069799 | 2.941633 | 0.03383 | 17.9 | 96.8 |
| | | 90,000 | 0.238232 | 4.630255 | 25.86044 | 0.146422 | 17.7 | 98.2 |
| | | 100,000 | 0.135878 | 1.856925 | 5.776062 | 0.058721 | 19.9 | 97.7 |
| | | 500,000 | 0.16588 | 0.097008 | 0.090787 | 0.003068 | 50.9 | 94.8 |
| | | 1,000,000 | 0.161506 | 0.062844 | 0.063408 | 0.001987 | 71.8 | 96.1 |
| α = 0.5 | θ = 0.167 | 10,000 | 0.019104 | 6.664308 | 64.97602 | 0.210744 | 6.1 | 99.1 |
| | | 20,000 | -0.06825 | 5.989795 | 78.92715 | 0.189414 | 8.4 | 98.5 |
| | | 30,000 | -0.07288 | 8.224195 | 124.1627 | 0.260072 | 10 | 99.3 |
| | | 40,000 | -0.2421 | 5.837098 | 64.99095 | 0.184585 | 12.6 | 98 |
| | | 50,000 | 0.237341 | 2.21805 | 10.19671 | 0.070141 | 12.8 | 99.2 |
| | | 60,000 | -0.06365 | 10.42808 | 390.4894 | 0.329765 | 14.7 | 98.3 |
| | | 70,000 | 0.165889 | 1.858064 | 9.151722 | 0.058757 | 12.6 | 98.6 |
| | | 80,000 | 0.209719 | 3.889168 | 30.41479 | 0.122986 | 18.1 | 97.1 |
| | | 90,000 | 0.101705 | 1.887371 | 6.084278 | 0.059684 | 18.3 | 98 |
| | | 100,000 | 0.144068 | 0.948483 | 1.494762 | 0.029994 | 21 | 97.7 |
| | | 500,000 | 0.164501 | 0.100782 | 0.093644 | 0.003187 | 49.8 | 94.8 |
| | | 1,000,000 | 0.160744 | 0.064345 | 0.064802 | 0.002035 | 68.9 | 96.3 |
| α = 1 | θ = 0.167 | 10,000 | 0.09368 | 5.892411 | 97.23889 | 0.186334 | 9.8 | 98.8 |
| | | 20,000 | -0.4304 | 20.97362 | 671.0983 | 0.663244 | 10.6 | 98.2 |
| | | 30,000 | 0.378562 | 5.425829 | 114.4574 | 0.17158 | 12.8 | 99 |
| | | 40,000 | 0.908258 | 15.35535 | 2832.478 | 0.485579 | 15.2 | 98.6 |
| | | 50,000 | 0.470457 | 10.01062 | 198.9186 | 0.316564 | 13.8 | 98.8 |
| | | 60,000 | 0.113022 | 2.900672 | 13.53292 | 0.091727 | 16.9 | 98.3 |
| | | 70,000 | 0.067771 | 6.009432 | 50.98357 | 0.190035 | 13.9 | 98.9 |
| | | 80,000 | -3.90401 | 89.58272 | 17960.33 | 2.832854 | 19.4 | 97.1 |

| | | | | | | | | |
|---|---|---|---|---|---|---|---|---|
| | | 90,000 | 0.111139 | 3.553606 | 39.86445 | 0.112375 | 20.1 | 98.4 |
| | | 100,000 | 0.031041 | 3.398025 | 21.44655 | 0.107455 | 20.8 | 97.2 |
| | | 500,000 | 0.157689 | 0.127015 | 0.109916 | 0.004017 | 48.1 | 95.3 |
| | | 1,000,000 | 0.157859 | 0.070483 | 0.070328 | 0.002229 | 63.7 | 97.1 |

#### 3.5 THETA=0.333

| Modelled mediator coefficient | Modelled interaction coefficient | Sample size | Mean 2sls interaction effect estimate | Standard deviation of 2sls estimate | Mean estimated standard error of 2sls estimate | Standard error of the bias (standard deviation of 2sls estimate/sqrt(1000)) | Power of 2sls estimator | Coverage of 2sls estimator |
|---|---|---|---|---|---|---|---|---|
| α = -0.333 | θ = 0.333 | 10,000 | 0.388677 | 7.81111 | 59.5645 | 0.247009 | 10.1 | 98.2 |
| | | 20,000 | -0.00044 | 4.786057 | 23.87388 | 0.151348 | 16.9 | 96.9 |
| | | 30,000 | 0.074812 | 11.36517 | 66.83278 | 0.359398 | 20.9 | 97.3 |
| | | 40,000 | 0.237734 | 0.761812 | 0.713607 | 0.024091 | 24.2 | 97.1 |
| | | 50,000 | 0.689144 | 14.34763 | 92.07353 | 0.453712 | 25.6 | 97.8 |
| | | 60,000 | 0.297163 | 0.335516 | 0.33689 | 0.01061 | 30.5 | 97.2 |
| | | 70,000 | 0.28384 | 0.304217 | 0.30987 | 0.00962 | 31 | 97.8 |
| | | 80,000 | 0.300442 | 0.289113 | 0.278051 | 0.009143 | 35.8 | 95.4 |
| | | 90,000 | 0.309931 | 0.253031 | 0.255318 | 0.008002 | 38.5 | 97.3 |
| | | 100,000 | 0.312586 | 0.242481 | 0.237091 | 0.007668 | 39.5 | 96.6 |
| | | 500,000 | 0.335112 | 0.099686 | 0.09483 | 0.003152 | 90.5 | 94.1 |
| | | 1,000,000 | 0.328643 | 0.066105 | 0.0671 | 0.00209 | 99.3 | 96 |
| α = 0 | θ = 0.333 | 10,000 | 0.467593 | 9.547763 | 77.35779 | 0.301927 | 14.3 | 96.8 |
| No mediation | | 20,000 | -1.20569 | 48.13992 | 1841.103 | 1.522318 | 18.9 | 95.5 |
| | | 30,000 | -0.58098 | 19.72037 | 454.804 | 0.623613 | 23.3 | 96.1 |
| | | 40,000 | 0.229565 | 2.276794 | 4.355157 | 0.071999 | 26.8 | 95.6 |
| | | 50,000 | 0.201656 | 1.222505 | 1.663323 | 0.038659 | 27.2 | 96.2 |
| | | 60,000 | 0.241504 | 1.243631 | 1.478072 | 0.039327 | 32.4 | 95.8 |
| | | 70,000 | 0.23939 | 0.544596 | 0.509121 | 0.017222 | 29.6 | 96.6 |
| | | 80,000 | 0.328062 | 3.247758 | 9.683401 | 0.102703 | 35.7 | 94.2 |
| | | 90,000 | 0.267286 | 0.575975 | 0.480611 | 0.018214 | 37 | 96.9 |
| | | 100,000 | 0.289935 | 0.30158 | 0.284954 | 0.009537 | 38 | 95.9 |
| | | 500,000 | 0.33239 | 0.107332 | 0.101637 | 0.003394 | 85.4 | 93.9 |
| | | 1,000,000 | 0.326701 | 0.070686 | 0.071743 | 0.002235 | 98 | 96.3 |

| | | | | | | | | |
|---|---|---|---|---|---|---|---|---|
| α = 0.333 | θ = 0.333 | 10,000 | 0.406406 | 9.692701 | 300.7414 | 0.30651 | 17.3 | 96.3 |
| | | 20,000 | 0.662585 | 7.127646 | 114.5098 | 0.225396 | 21 | 95.4 |
| | | 30,000 | 0.072835 | 25.56029 | 1356.877 | 0.808287 | 24.7 | 96.1 |
| | | 40,000 | 0.124167 | 3.254435 | 16.93763 | 0.102914 | 27.6 | 95.2 |
| | | 50,000 | -0.0072 | 7.734129 | 167.1191 | 0.244575 | 28.3 | 95.8 |
| | | 60,000 | 0.324337 | 2.218865 | 6.158413 | 0.070167 | 32.5 | 95.1 |
| | | 70,000 | 0.013381 | 7.669513 | 370.5341 | 0.242531 | 29.9 | 96.2 |
| | | 80,000 | 0.204955 | 1.616596 | 5.710002 | 0.051121 | 35.2 | 93.6 |
| | | 90,000 | 0.490959 | 7.026869 | 39.10739 | 0.222209 | 36.3 | 96 |
| | | 100,000 | 0.266289 | 3.183921 | 10.13468 | 0.100684 | 37.8 | 95.2 |
| | | 500,000 | 0.328315 | 0.118434 | 0.111204 | 0.003745 | 79.8 | 93.7 |
| | | 1,000,000 | 0.324207 | 0.076521 | 0.077616 | 0.00242 | 94.9 | 96.5 |
| α = 0.5 | θ = 0.333 | 10,000 | 0.225774 | 7.86307 | 74.85897 | 0.248652 | 18.8 | 96 |
| | | 20,000 | -0.00343 | 7.254742 | 76.93808 | 0.229415 | 21.7 | 94.8 |
| | | 30,000 | -0.04139 | 12.34687 | 192.8554 | 0.390442 | 25 | 95.9 |
| | | 40,000 | -0.29291 | 9.424248 | 109.2822 | 0.298021 | 28.9 | 95.1 |
| | | 50,000 | 0.45392 | 3.894822 | 19.33617 | 0.123165 | 29 | 95.6 |
| | | 60,000 | 0.090382 | 18.88432 | 787.2233 | 0.597175 | 33.4 | 95.1 |
| | | 70,000 | 0.38567 | 3.882381 | 20.04184 | 0.122772 | 29.5 | 95.8 |
| | | 80,000 | 0.472839 | 6.986196 | 55.97751 | 0.220923 | 35.3 | 93.3 |
| | | 90,000 | 0.188918 | 3.072642 | 9.509641 | 0.097165 | 36.3 | 95.6 |
| | | 100,000 | 0.269167 | 1.589925 | 2.490781 | 0.050278 | 37.5 | 94.8 |
| | | 500,000 | 0.325593 | 0.126034 | 0.117429 | 0.003986 | 76.2 | 93.6 |
| | | 1,000,000 | 0.32272 | 0.080001 | 0.081074 | 0.00253 | 92.8 | 96.5 |
| α = 1 | θ = 0.333 | 10,000 | 0.325684 | 6.551942 | 104.6608 | 0.207191 | 23.5 | 94.5 |
| | | 20,000 | -0.40745 | 28.17452 | 901.663 | 0.890957 | 25 | 94.3 |
| | | 30,000 | 0.743033 | 9.53234 | 203.269 | 0.301439 | 27.1 | 93.8 |
| | | 40,000 | 2.434921 | 53.07075 | 22498.55 | 1.678245 | 30.9 | 94.3 |
| | | 50,000 | 0.835201 | 12.08151 | 225.2215 | 0.382051 | 30.7 | 95.1 |
| | | 60,000 | 0.305374 | 3.647052 | 18.05817 | 0.11533 | 34.1 | 94 |
| | | 70,000 | 0.28448 | 7.165697 | 64.43737 | 0.226599 | 29.7 | 95 |
| | | 80,000 | -6.2537 | 148.4059 | 30556.66 | 4.693005 | 35.4 | 92.8 |

| | | | | | | | | |
|---|---|---|---|---|---|---|---|---|
| | | 90,000 | 0.333184 | 6.799938 | 93.15192 | 0.215033 | 36.6 | 94.6 |
| | | 100,000 | 0.208895 | 3.823617 | 19.64807 | 0.120913 | 38.5 | 94.1 |
| | | 500,000 | 0.312328 | 0.173982 | 0.149289 | 0.005502 | 67.8 | 92.5 |
| | | 1,000,000 | 0.317093 | 0.093453 | 0.093978 | 0.002955 | 84.8 | 96.3 |

### 4 FACTORIAL MR

#### 4.1 THETA=-0.111

| Modelled mediator coefficient | Modelled interaction coefficient | Sample size | Power of FMR to detect an interaction (%) |
|---|---|---|---|
| α = -0.333 | θ = -0.111 | 10,000 | 5.1 |
| | | 20,000 | 4.4 |
| | | 30,000 | 4.2 |
| | | 40,000 | 7.1 |
| | | 50,000 | 6.3 |
| | | 60,000 | 7.8 |
| | | 70,000 | 6.3 |
| | | 80,000 | 6.5 |
| | | 90,000 | 7.7 |
| | | 100,000 | 6.5 |
| | | 500,000 | 16.6 |
| | | 1,000,000 | 29.2 |
| α = 0 | θ = -0.111 | 10,000 | 5.6 |
| No mediation | | 20,000 | 4.5 |
| | | 30,000 | 4.2 |
| | | 40,000 | 6.9 |
| | | 50,000 | 6.6 |
| | | 60,000 | 7.4 |
| | | 70,000 | 6.3 |
| | | 80,000 | 6.7 |
| | | 90,000 | 7.7 |
| | | 100,000 | 6.7 |
| | | 500,000 | 16.2 |
| | | 1,000,000 | 28.9 |
| α = 0.333 | θ = -0.111 | 10,000 | 5.5 |
| | | 20,000 | 4.9 |
| | | 30,000 | 4.6 |
| | | 40,000 | 6.5 |
| | | 50,000 | 6.2 |
| | | 60,000 | 7.2 |
| | | 70,000 | 6.4 |
| | | 80,000 | 6.7 |
| | | 90,000 | 7.4 |
| | | 100,000 | 6.9 |
| | | 500,000 | 14.9 |
| | | 1,000,000 | 26.9 |
| α = 0.5 | θ = -0.111 | 10,000 | 5.4 |
| | | 20,000 | 4.5 |
| | | 30,000 | 4.7 |
| | | 40,000 | 6.2 |
| | | 50,000 | 6.2 |
| | | 60,000 | 7.7 |
| | | 70,000 | 6.5 |
| | | 80,000 | 6.6 |

| | | | |
|---|---|---|---|
| | | 90,000 | 7.2 |
| | | 100,000 | 6.8 |
| | | 500,000 | 14.5 |
| | | 1,000,000 | 24.8 |
| α = 1 | θ = -0.111 | 10,000 | 5.6 |
| | | 20,000 | 4.2 |
| | | 30,000 | 4.5 |
| | | 40,000 | 5.9 |
| | | 50,000 | 5.4 |
| | | 60,000 | 7 |
| | | 70,000 | 6 |
| | | 80,000 | 6.7 |
| | | 90,000 | 6.6 |
| | | 100,000 | 7.7 |
| | | 500,000 | 12.8 |
| | | 1,000,000 | 20 |
| Power and type I error defined using F statistic and Wald test | | | |

#### 4.2 THETA=0

| Modelled mediator coefficient | Modelled interaction coefficient | Sample size | Type I error of FMR to detect an interaction (%) |
|---|---|---|---|
| α = -0.333 | θ = 0 | 10,000 | 5.2 |
| | | 20,000 | 4.7 |
| | | 30,000 | 5.8 |
| | | 40,000 | 5.9 |
| | | 50,000 | 5.7 |
| | | 60,000 | 5.1 |
| | | 70,000 | 4.2 |
| | | 80,000 | 4.6 |
| | | 90,000 | 4.5 |
| | | 100,000 | 6 |
| | | 500,000 | 5.8 |
| | | 1,000,000 | 4.7 |
| α = 0 | θ = 0 | 10,000 | 5.2 |
| No mediation | | 20,000 | 4.7 |
| | | 30,000 | 5.8 |
| | | 40,000 | 5.9 |
| | | 50,000 | 5.7 |
| | | 60,000 | 5.1 |
| | | 70,000 | 4.2 |
| | | 80,000 | 4.6 |
| | | 90,000 | 4.5 |
| | | 100,000 | 6 |
| | | 500,000 | 5.8 |
| | | 1,000,000 | 4.7 |
| α = 0.333 | θ = 0 | 10,000 | 5.2 |
| | | 20,000 | 4.7 |
| | | 30,000 | 5.8 |
| | | 40,000 | 5.9 |
| | | 50,000 | 5.7 |
| | | 60,000 | 5.1 |
| | | 70,000 | 4.2 |
| | | 80,000 | 4.6 |
| | | 90,000 | 4.5 |
| | | 100,000 | 6 |
| | | 500,000 | 5.8 |
| | | 1,000,000 | 4.7 |
| α = 0.5 | θ = 0 | 10,000 | 5.2 |
| | | 20,000 | 4.7 |
| | | 30,000 | 5.8 |
| | | 40,000 | 5.9 |
| | | 50,000 | 5.7 |
| | | 60,000 | 5.1 |
| | | 70,000 | 4.2 |
| | | 80,000 | 4.6 |

| | | | |
|---|---|---|---|
| | | 90,000 | 4.5 |
| | | 100,000 | 6 |
| | | 500,000 | 5.8 |
| | | 1,000,000 | 4.7 |
| α = 1 | θ = 0 | 10,000 | 5.2 |
| | | 20,000 | 4.7 |
| | | 30,000 | 5.8 |
| | | 40,000 | 5.9 |
| | | 50,000 | 5.7 |
| | | 60,000 | 5.1 |
| | | 70,000 | 4.2 |
| | | 80,000 | 4.6 |
| | | 90,000 | 4.5 |
| | | 100,000 | 6 |
| | | 500,000 | 5.8 |
| | | 1,000,000 | 4.7 |
| Power and type I error defined using F statistic and Wald test | | | |

#### 4.3 THETA=0.111

| Modelled mediator coefficient | Modelled interaction coefficient | Sample size | Power of FMR to detect an interaction (%) |
|---|---|---|---|
| α = -0.333 | θ = 0.111 | 10,000 | 5.2 |
| | | 20,000 | 6.3 |
| | | 30,000 | 6.3 |
| | | 40,000 | 5.7 |
| | | 50,000 | 5.4 |
| | | 60,000 | 6.1 |
| | | 70,000 | 4.8 |
| | | 80,000 | 5.8 |
| | | 90,000 | 5.8 |
| | | 100,000 | 6.5 |
| | | 500,000 | 11 |
| | | 1,000,000 | 15.2 |
| α = 0 | θ = 0.111 | 10,000 | 5.1 |
| No mediation | | 20,000 | 6.3 |
| | | 30,000 | 6.4 |
| | | 40,000 | 5.7 |
| | | 50,000 | 5.5 |
| | | 60,000 | 5.6 |
| | | 70,000 | 4.8 |
| | | 80,000 | 5.7 |
| | | 90,000 | 5.7 |
| | | 100,000 | 6.4 |
| | | 500,000 | 10 |
| | | 1,000,000 | 13.1 |
| α = 0.333 | θ = 0.111 | 10,000 | 4.7 |
| | | 20,000 | 6.4 |
| | | 30,000 | 6.1 |
| | | 40,000 | 5.5 |
| | | 50,000 | 5 |
| | | 60,000 | 5.4 |
| | | 70,000 | 4.9 |
| | | 80,000 | 5.6 |
| | | 90,000 | 5.3 |
| | | 100,000 | 6.5 |
| | | 500,000 | 9.6 |
| | | 1,000,000 | 11.5 |
| α = 0.5 | θ = 0.111 | 10,000 | 4.5 |
| | | 20,000 | 6.5 |
| | | 30,000 | 6 |
| | | 40,000 | 5.6 |
| | | 50,000 | 4.9 |
| | | 60,000 | 5.5 |
| | | 70,000 | 4.9 |
| | | 80,000 | 5.6 |

| | | | |
|---|---|---|---|
| | | 90,000 | 5.3 |
| | | 100,000 | 6 |
| | | 500,000 | 9.1 |
| | | 1,000,000 | 10.5 |
| α = 1 | θ = 0.111 | 10,000 | 4.5 |
| | | 20,000 | 6 |
| | | 30,000 | 6.2 |
| | | 40,000 | 5.2 |
| | | 50,000 | 4.6 |
| | | 60,000 | 5.1 |
| | | 70,000 | 4.7 |
| | | 80,000 | 5.6 |
| | | 90,000 | 4.5 |
| | | 100,000 | 5.8 |
| | | 500,000 | 8 |
| | | 1,000,000 | 9.1 |
| Power and type I error defined using F statistic and Wald test | | | |

#### 4.4 THETA=0.167

| Modelled mediator coefficient | Modelled interaction coefficient | Sample size | Power of FMR to detect an interaction (%) |
|---|---|---|---|
| α = -0.333 | θ = 0.167 | 10,000 | 4.9 |
| | | 20,000 | 6.5 |
| | | 30,000 | 6.6 |
| | | 40,000 | 6 |
| | | 50,000 | 5.9 |
| | | 60,000 | 6.4 |
| | | 70,000 | 5.9 |
| | | 80,000 | 6.5 |
| | | 90,000 | 5.8 |
| | | 100,000 | 6.6 |
| | | 500,000 | 13.8 |
| | | 1,000,000 | 21.7 |
| α = 0 | θ = 0.167 | 10,000 | 4.9 |
| No mediation | | 20,000 | 6.2 |
| | | 30,000 | 6.6 |
| | | 40,000 | 5.7 |
| | | 50,000 | 5.3 |
| | | 60,000 | 5.9 |
| | | 70,000 | 5.3 |
| | | 80,000 | 6.2 |
| | | 90,000 | 5.6 |
| | | 100,000 | 6 |
| | | 500,000 | 11.8 |
| | | 1,000,000 | 18.8 |
| α = 0.333 | θ = 0.167 | 10,000 | 4.4 |
| | | 20,000 | 5.9 |
| | | 30,000 | 6.7 |
| | | 40,000 | 5.4 |
| | | 50,000 | 5.3 |
| | | 60,000 | 5.3 |
| | | 70,000 | 5.4 |
| | | 80,000 | 5.7 |
| | | 90,000 | 5.3 |
| | | 100,000 | 6 |
| | | 500,000 | 10.2 |
| | | 1,000,000 | 15.8 |
| α = 0.5 | θ = 0.167 | 10,000 | 4.5 |
| | | 20,000 | 5.8 |
| | | 30,000 | 6.3 |
| | | 40,000 | 5.8 |
| | | 50,000 | 5.3 |
| | | 60,000 | 5.5 |
| | | 70,000 | 5.2 |
| | | 80,000 | 5.9 |

| | | | |
|---|---|---|---|
| | | 90,000 | 5.4 |
| | | 100,000 | 6.1 |
| | | 500,000 | 9.9 |
| | | 1,000,000 | 14.1 |
| α = 1 | θ = 0.167 | 10,000 | 4.8 |
| | | 20,000 | 5.9 |
| | | 30,000 | 6.2 |
| | | 40,000 | 5.8 |
| | | 50,000 | 4.9 |
| | | 60,000 | 5.4 |
| | | 70,000 | 5.5 |
| | | 80,000 | 5.8 |
| | | 90,000 | 5.5 |
| | | 100,000 | 5.7 |
| | | 500,000 | 8.4 |
| | | 1,000,000 | 11.5 |
| Power and type I error defined using F statistic and Wald test | | | |

#### 4.5 THETA=0.333

| Modelled mediator coefficient | Modelled interaction coefficient | Sample size | Power of FMR to detect an interaction (%) |
|---|---|---|---|
| α = -0.333 | θ = 0.333 | 10,000 | 4.8 |
| | | 20,000 | 6.7 |
| | | 30,000 | 6.8 |
| | | 40,000 | 6.2 |
| | | 50,000 | 6.7 |
| | | 60,000 | 7.2 |
| | | 70,000 | 6.3 |
| | | 80,000 | 6.5 |
| | | 90,000 | 7.1 |
| | | 100,000 | 7.3 |
| | | 500,000 | 22.5 |
| | | 1,000,000 | 37 |
| α = 0 | θ = 0.333 | 10,000 | 4.5 |
| No mediation | | 20,000 | 6.1 |
| | | 30,000 | 6.4 |
| | | 40,000 | 5.9 |
| | | 50,000 | 6.3 |
| | | 60,000 | 6.9 |
| | | 70,000 | 6.2 |
| | | 80,000 | 6.3 |
| | | 90,000 | 7 |
| | | 100,000 | 7.2 |
| | | 500,000 | 17.4 |
| | | 1,000,000 | 28.1 |
| α = 0.333 | θ = 0.333 | 10,000 | 4.5 |
| | | 20,000 | 5.9 |
| | | 30,000 | 6.5 |
| | | 40,000 | 5.8 |
| | | 50,000 | 5.8 |
| | | 60,000 | 6.5 |
| | | 70,000 | 5.8 |
| | | 80,000 | 6 |
| | | 90,000 | 6.7 |
| | | 100,000 | 6.8 |
| | | 500,000 | 14.1 |
| | | 1,000,000 | 21.8 |
| α = 0.5 | θ = 0.333 | 10,000 | 4.8 |
| | | 20,000 | 5.4 |
| | | 30,000 | 6.7 |
| | | 40,000 | 5.7 |
| | | 50,000 | 5.6 |
| | | 60,000 | 6.5 |
| | | 70,000 | 6 |
| | | 80,000 | 5.7 |

| | | | |
|---|---|---|---|
| | | 90,000 | 6.8 |
| | | 100,000 | 6.5 |
| | | 500,000 | 13.2 |
| | | 1,000,000 | 20.3 |
| α = 1 | θ = 0.333 | 10,000 | 4.9 |
| | | 20,000 | 5.4 |
| | | 30,000 | 6.7 |
| | | 40,000 | 5.7 |
| | | 50,000 | 5.8 |
| | | 60,000 | 6.4 |
| | | 70,000 | 5 |
| | | 80,000 | 5.2 |
| | | 90,000 | 6.2 |
| | | 100,000 | 5.9 |
| | | 500,000 | 10.4 |
| | | 1,000,000 | 14.8 |
| Power and type I error defined using F statistic and Wald test | | | |

### 5 2SLS INSTRUMENT ASSUMES MEDIATION, Z=(Z1,Z2,Z1Z2,Z1Z1)

#### 5.1 THETA=-0.111

| Modelled mediator coefficient | Modelled interaction coefficient | Sample size | Mean 2sls interaction effect estimate | Standard deviation of 2sls estimate | Mean estimated standard error of 2sls estimate | Standard error of the bias (standard deviation of 2sls estimate/sqrt(1000)) | Power of 2sls estimator (%) | Coverage of 2sls estimator (%) |
|---|---|---|---|---|---|---|---|---|
| α = -0.333 | θ = -0.111 | 10,000 | -0.18706 | 1.076519 | 1.48131 | 0.034043 | 2.3 | 99.5 |
| | | 20,000 | -0.11597 | 0.598166 | 0.646303 | 0.018916 | 3.4 | 99.4 |
| | | 30,000 | -0.10924 | 0.454739 | 0.465493 | 0.01438 | 5.2 | 98.4 |
| | | 40,000 | -0.12039 | 0.316235 | 0.320225 | 0.01 | 8.8 | 97.6 |
| | | 50,000 | -0.12401 | 0.279135 | 0.281334 | 0.008827 | 8.1 | 97.8 |
| | | 60,000 | -0.11514 | 0.2304 | 0.239271 | 0.007286 | 9.1 | 98 |
| | | 70,000 | -0.11283 | 0.220268 | 0.224387 | 0.006965 | 9.5 | 97.3 |
| | | 80,000 | -0.11624 | 0.216432 | 0.206372 | 0.006844 | 10.1 | 97.5 |
| | | 90,000 | -0.10729 | 0.195853 | 0.19348 | 0.006193 | 9.3 | 97.3 |
| | | 100,000 | -0.10355 | 0.183141 | 0.18348 | 0.005791 | 10.9 | 97.6 |
| | | 500,000 | -0.10763 | 0.079258 | 0.076689 | 0.002506 | 30.9 | 94.8 |
| | | 1,000,000 | -0.11182 | 0.053775 | 0.054206 | 0.001701 | 56.4 | 95.4 |
| α = 0 | θ = -0.111 | 10,000 | -0.20216 | 0.986655 | 1.498846 | 0.031201 | 3.7 | 99.1 |
| No mediation | | 20,000 | -0.12065 | 0.74251 | 0.979235 | 0.02348 | 4.1 | 99 |
| | | 30,000 | -0.13328 | 0.72618 | 0.786817 | 0.022964 | 5.7 | 98.3 |
| | | 40,000 | -0.14241 | 0.433761 | 0.456131 | 0.013717 | 8.1 | 97.9 |
| | | 50,000 | -0.14925 | 0.334484 | 0.339586 | 0.010577 | 8.8 | 97.7 |
| | | 60,000 | -0.12349 | 0.295504 | 0.306463 | 0.009345 | 8.9 | 98.1 |
| | | 70,000 | -0.11151 | 0.255225 | 0.267764 | 0.008071 | 9.9 | 98.1 |
| | | 80,000 | -0.11771 | 0.304805 | 0.27821 | 0.009639 | 10.2 | 98.3 |
| | | 90,000 | -0.10535 | 0.229623 | 0.223587 | 0.007261 | 8.9 | 98.2 |
| | | 100,000 | -0.10737 | 0.208313 | 0.205176 | 0.006587 | 10.5 | 98.1 |
| | | 500,000 | -0.10839 | 0.086107 | 0.083045 | 0.002723 | 29.7 | 94.5 |
| | | 1,000,000 | -0.11173 | 0.057225 | 0.05864 | 0.00181 | 49.6 | 96 |

| | | | | | | | | |
|---|---|---|---|---|---|---|---|---|
| α = 0.333 | θ = -0.111 | 10,000 | -0.21154 | 0.765203 | 1.298475 | 0.024198 | 5.1 | 99 |
| | | 20,000 | -0.16806 | 0.575466 | 0.806974 | 0.018198 | 5.8 | 99.2 |
| | | 30,000 | -0.15945 | 0.840079 | 1.161619 | 0.026566 | 6.8 | 98.3 |
| | | 40,000 | -0.14832 | 0.417362 | 0.511085 | 0.013198 | 10 | 98.7 |
| | | 50,000 | -0.15869 | 0.52521 | 0.466656 | 0.016609 | 9.8 | 97.3 |
| | | 60,000 | -0.1373 | 0.353551 | 0.368441 | 0.01118 | 10.6 | 98.6 |
| | | 70,000 | -0.11488 | 0.340849 | 0.356105 | 0.010779 | 12 | 98 |
| | | 80,000 | -0.12547 | 0.227594 | 0.245195 | 0.007197 | 11.4 | 98.5 |
| | | 90,000 | -0.10681 | 0.255881 | 0.259467 | 0.008092 | 12 | 97.9 |
| | | 100,000 | -0.12069 | 0.357219 | 0.293045 | 0.011296 | 13.1 | 98.2 |
| | | 500,000 | -0.10922 | 0.081828 | 0.079741 | 0.002588 | 32.8 | 95.6 |
| | | 1,000,000 | -0.11149 | 0.054341 | 0.055932 | 0.001718 | 53 | 95.2 |
| α = 0.5 | θ = -0.111 | 10,000 | -0.1875 | 0.699966 | 1.109288 | 0.022135 | 5.5 | 99.1 |
| | | 20,000 | -0.16198 | 0.465365 | 0.593426 | 0.014716 | 6.5 | 98.7 |
| | | 30,000 | -0.16132 | 0.604987 | 0.789674 | 0.019131 | 8.7 | 98.2 |
| | | 40,000 | -0.14376 | 0.358825 | 0.423346 | 0.011347 | 10.7 | 98.8 |
| | | 50,000 | -0.14444 | 0.34384 | 0.382541 | 0.010873 | 10.8 | 97.2 |
| | | 60,000 | -0.14044 | 0.279346 | 0.309524 | 0.008834 | 12.2 | 98.5 |
| | | 70,000 | -0.11983 | 0.287092 | 0.290708 | 0.009079 | 12.5 | 97.9 |
| | | 80,000 | -0.12279 | 0.242507 | 0.267138 | 0.007669 | 13.5 | 98.8 |
| | | 90,000 | -0.1048 | 0.223925 | 0.234768 | 0.007081 | 13.4 | 97.7 |
| | | 100,000 | -0.1122 | 0.206471 | 0.199921 | 0.006529 | 14.2 | 98.1 |
| | | 500,000 | -0.10963 | 0.076283 | 0.075222 | 0.002412 | 35.8 | 95.8 |
| | | 1,000,000 | -0.11126 | 0.051235 | 0.052565 | 0.00162 | 59.5 | 95.2 |
| α = 1 | θ = -0.111 | 10,000 | -0.18726 | 0.490068 | 0.950795 | 0.015497 | 9.8 | 98.6 |
| | | 20,000 | -0.14977 | 0.397464 | 0.608196 | 0.012569 | 10.5 | 98.3 |
| | | 30,000 | -0.14452 | 0.364937 | 0.491593 | 0.01154 | 15 | 97.8 |
| | | 40,000 | -0.13845 | 0.305051 | 0.36495 | 0.009647 | 15.6 | 98 |
| | | 50,000 | -0.13884 | 0.25266 | 0.3258 | 0.00799 | 18.8 | 96.9 |
| | | 60,000 | -0.12785 | 0.262517 | 0.296885 | 0.008302 | 17.4 | 98 |
| | | 70,000 | -0.12724 | 0.495161 | 0.495209 | 0.015658 | 17.9 | 97.8 |
| | | 80,000 | -0.12058 | 0.228902 | 0.234561 | 0.007239 | 19.9 | 97.9 |

| | | | | | | | | |
|---|---|---|---|---|---|---|---|---|
| | | 90,000 | -0.10064 | 0.249745 | 0.248568 | 0.007898 | 20.9 | 96.7 |
| | | 100,000 | -0.11763 | 0.162328 | 0.163181 | 0.005133 | 23.2 | 97.7 |
| | | 500,000 | -0.11067 | 0.058644 | 0.059315 | 0.001854 | 52 | 95.3 |
| | | 1,000,000 | -0.11069 | 0.040609 | 0.041169 | 0.001284 | 76.9 | 95.7 |

#### 5.2 THETA=0

| Modelled mediator coefficient | Modelled interaction coefficient | Sample size | Mean 2sls interaction effect estimate | Standard deviation of 2sls estimate | Mean estimated standard error of 2sls estimate | Standard error of the bias (standard deviation of 2sls estimate/sqrt(1000)) | Type I error of 2sls estimator (%) | Coverage of 2sls estimator (%) |
|---|---|---|---|---|---|---|---|---|
| α = -0.333 | θ = 0 | 10,000 | -0.03339 | 1.028665 | 1.436209 | 0.032529 | 0.1 | 99.9 |
| | | 20,000 | 0.005382 | 0.601347 | 0.642348 | 0.019016 | 0.3 | 99.7 |
| | | 30,000 | 0.001548 | 0.441551 | 0.454041 | 0.013963 | 0.8 | 99.2 |
| | | 40,000 | -0.00575 | 0.310096 | 0.312888 | 0.009806 | 1.6 | 98.4 |
| | | 50,000 | -0.01317 | 0.276796 | 0.276383 | 0.008753 | 1.9 | 98.1 |
| | | 60,000 | -5.92E-05 | 0.224856 | 0.23396 | 0.007111 | 1.6 | 98.4 |
| | | 70,000 | -0.00413 | 0.214716 | 0.219423 | 0.00679 | 2 | 98 |
| | | 80,000 | -0.0034 | 0.20984 | 0.201842 | 0.006636 | 2.5 | 97.5 |
| | | 90,000 | 0.001843 | 0.188239 | 0.188744 | 0.005953 | 2.4 | 97.6 |
| | | 100,000 | 0.005428 | 0.179205 | 0.179321 | 0.005667 | 2 | 98 |
| | | 500,000 | 0.003899 | 0.078714 | 0.075032 | 0.002489 | 5.1 | 94.9 |
| | | 1,000,000 | -0.00128 | 0.052679 | 0.05307 | 0.001666 | 4.5 | 95.5 |
| α = 0 | θ = 0 | 10,000 | -0.01961 | 1.010714 | 1.537807 | 0.031962 | 0 | 100 |
| No mediation | | 20,000 | 0.024202 | 0.689174 | 0.90307 | 0.021794 | 0.4 | 99.6 |
| | | 30,000 | -0.00248 | 0.67781 | 0.752259 | 0.021434 | 0.2 | 99.8 |
| | | 40,000 | -0.01519 | 0.43524 | 0.456942 | 0.013763 | 0.7 | 99.3 |
| | | 50,000 | -0.02921 | 0.344766 | 0.343028 | 0.010902 | 0.8 | 99.2 |
| | | 60,000 | -0.00605 | 0.306988 | 0.308644 | 0.009708 | 0.8 | 99.2 |
| | | 70,000 | -0.00076 | 0.239685 | 0.253623 | 0.007579 | 1.2 | 98.8 |
| | | 80,000 | -0.00409 | 0.281678 | 0.264636 | 0.008907 | 1.3 | 98.7 |
| | | 90,000 | 0.005483 | 0.21645 | 0.214791 | 0.006845 | 1.7 | 98.3 |
| | | 100,000 | 0.003484 | 0.203867 | 0.198973 | 0.006447 | 1.3 | 98.7 |
| | | 500,000 | 0.003411 | 0.084527 | 0.080513 | 0.002673 | 6 | 94 |
| | | 1,000,000 | -0.0014 | 0.056028 | 0.056911 | 0.001772 | 4.6 | 95.4 |

| | | | | | | | | |
|---|---|---|---|---|---|---|---|---|
| α = 0.333 | θ = 0 | 10,000 | -0.01446 | 0.773106 | 1.325373 | 0.024448 | 0.1 | 99.9 |
| | | 20,000 | -0.00123 | 0.533524 | 0.757063 | 0.016871 | 0.4 | 99.6 |
| | | 30,000 | -0.00839 | 0.790478 | 1.116682 | 0.024997 | 0 | 100 |
| | | 40,000 | -0.00612 | 0.444997 | 0.536436 | 0.014072 | 0.4 | 99.6 |
| | | 50,000 | -0.02799 | 0.483952 | 0.445904 | 0.015304 | 0.3 | 99.7 |
| | | 60,000 | -0.01033 | 0.336787 | 0.353051 | 0.01065 | 0.4 | 99.6 |
| | | 70,000 | 0.002013 | 0.264741 | 0.299273 | 0.008372 | 0.3 | 99.7 |
| | | 80,000 | -0.00888 | 0.230955 | 0.237422 | 0.007303 | 0.9 | 99.1 |
| | | 90,000 | 0.007127 | 0.257261 | 0.255802 | 0.008135 | 0.4 | 99.6 |
| | | 100,000 | -0.00234 | 0.265298 | 0.249449 | 0.008389 | 0.8 | 99.2 |
| | | 500,000 | 0.002546 | 0.078625 | 0.076435 | 0.002486 | 5.2 | 94.8 |
| | | 1,000,000 | -0.00113 | 0.0529 | 0.053714 | 0.001673 | 4.7 | 95.3 |
| α = 0.5 | θ = 0 | 10,000 | 0.004062 | 0.664833 | 1.066352 | 0.021024 | 0.1 | 99.9 |
| | | 20,000 | 0.004738 | 0.439981 | 0.557946 | 0.013913 | 0.4 | 99.6 |
| | | 30,000 | -0.01706 | 0.531479 | 0.720319 | 0.016807 | 0 | 100 |
| | | 40,000 | -0.00276 | 0.346847 | 0.402602 | 0.010968 | 0.2 | 99.8 |
| | | 50,000 | -0.01849 | 0.357505 | 0.486435 | 0.011305 | 0.2 | 99.8 |
| | | 60,000 | -0.01426 | 0.270824 | 0.296974 | 0.008564 | 0.4 | 99.6 |
| | | 70,000 | 0.000592 | 0.253393 | 0.269395 | 0.008013 | 0.3 | 99.7 |
| | | 80,000 | -0.00565 | 0.23951 | 0.260929 | 0.007574 | 0.7 | 99.3 |
| | | 90,000 | 0.010253 | 0.206037 | 0.21955 | 0.006515 | 0.5 | 99.5 |
| | | 100,000 | 0.004624 | 0.192501 | 0.189997 | 0.006087 | 1 | 99 |
| | | 500,000 | 0.001997 | 0.072593 | 0.071647 | 0.002296 | 4.7 | 95.3 |
| | | 1,000,000 | -0.00083 | 0.049471 | 0.050173 | 0.001564 | 4.7 | 95.3 |
| α = 1 | θ = 0 | 10,000 | -0.0037 | 0.467823 | 0.905348 | 0.014794 | 0.1 | 99.9 |
| | | 20,000 | 0.018115 | 0.35524 | 0.513046 | 0.011234 | 0.4 | 99.6 |
| | | 30,000 | 0.003369 | 0.373595 | 0.510629 | 0.011814 | 0 | 100 |
| | | 40,000 | -0.00253 | 0.306245 | 0.35916 | 0.009684 | 0.4 | 99.6 |
| | | 50,000 | -0.00935 | 0.23839 | 0.295923 | 0.007539 | 0.2 | 99.8 |
| | | 60,000 | -0.00144 | 0.239451 | 0.279512 | 0.007572 | 0.3 | 99.7 |
| | | 70,000 | -0.00055 | 0.391558 | 0.40733 | 0.012382 | 0.6 | 99.4 |
| | | 80,000 | -0.00416 | 0.207396 | 0.209482 | 0.006558 | 0.7 | 99.3 |

| | | | | | | | | |
|---|---|---|---|---|---|---|---|---|
| | | 90,000 | 0.012204 | 0.197349 | 0.211747 | 0.006241 | 0.5 | 99.5 |
| | | 100,000 | -0.00019 | 0.150885 | 0.152935 | 0.004771 | 1.1 | 98.9 |
| | | 500,000 | 0.000492 | 0.055034 | 0.05541 | 0.00174 | 4 | 96 |
| | | 1,000,000 | -3.96E-05 | 0.037923 | 0.038476 | 0.001199 | 4.1 | 95.9 |

#### 5.3 THETA=0.111

| Modelled mediator coefficient | Modelled interaction coefficient | Sample size | Mean 2sls interaction effect estimate | Standard deviation of 2sls estimate | Mean estimated standard error of 2sls estimate | Standard error of the bias (standard deviation of 2sls estimate/sqrt(1000)) | Power of 2sls estimator (%) | Coverage of 2sls estimator (%) |
|---|---|---|---|---|---|---|---|---|
| α = -0.333 | θ = 0.111 | 10,000 | 0.120269 | 1.010892 | 1.417101 | 0.031967 | 2.4 | 99.5 |
| | | 20,000 | 0.126731 | 0.628127 | 0.658104 | 0.019863 | 3.7 | 99.2 |
| | | 30,000 | 0.112331 | 0.451049 | 0.464658 | 0.014263 | 4.3 | 98.9 |
| | | 40,000 | 0.108884 | 0.318172 | 0.319714 | 0.010061 | 6.6 | 97.8 |
| | | 50,000 | 0.097668 | 0.290143 | 0.287094 | 0.009175 | 7.5 | 97.5 |
| | | 60,000 | 0.115022 | 0.230645 | 0.239489 | 0.007294 | 7.8 | 97.6 |
| | | 70,000 | 0.104572 | 0.21962 | 0.224498 | 0.006945 | 6.8 | 98 |
| | | 80,000 | 0.109442 | 0.213109 | 0.206112 | 0.006739 | 10.5 | 97.2 |
| | | 90,000 | 0.110973 | 0.190894 | 0.192683 | 0.006037 | 10.4 | 97 |
| | | 100,000 | 0.114406 | 0.183568 | 0.183199 | 0.005805 | 10.6 | 97.5 |
| | | 500,000 | 0.115423 | 0.081204 | 0.07659 | 0.002568 | 35.8 | 94.4 |
| | | 1,000,000 | 0.109258 | 0.053775 | 0.054225 | 0.001701 | 51.9 | 95.4 |
| α = 0 | θ = 0.111 | 10,000 | 0.162937 | 1.06187 | 1.605094 | 0.033579 | 3 | 99.5 |
| No mediation | | 20,000 | 0.169053 | 0.680691 | 0.876537 | 0.021525 | 4.7 | 99 |
| | | 30,000 | 0.128316 | 0.67302 | 0.75667 | 0.021283 | 4.9 | 98.9 |
| | | 40,000 | 0.112028 | 0.464982 | 0.481922 | 0.014704 | 6.7 | 98.4 |
| | | 50,000 | 0.090828 | 0.378685 | 0.365357 | 0.011975 | 7.1 | 99.4 |
| | | 60,000 | 0.11139 | 0.341494 | 0.327837 | 0.010799 | 8 | 98.1 |
| | | 70,000 | 0.11 | 0.253138 | 0.265954 | 0.008005 | 6.7 | 98.5 |
| | | 80,000 | 0.109532 | 0.274905 | 0.266797 | 0.008693 | 11.3 | 96.9 |
| | | 90,000 | 0.11632 | 0.217853 | 0.22034 | 0.006889 | 11 | 97.9 |
| | | 100,000 | 0.114334 | 0.211933 | 0.205414 | 0.006702 | 11.4 | 97.9 |
| | | 500,000 | 0.115209 | 0.087647 | 0.082886 | 0.002772 | 32.3 | 94.4 |
| | | 1,000,000 | 0.108941 | 0.058118 | 0.058694 | 0.001838 | 46.9 | 95 |

| | | | | | | | | |
|---|---|---|---|---|---|---|---|---|
| α = 0.333 | θ = 0.111 | 10,000 | 0.182614 | 0.823528 | 1.381805 | 0.026042 | 4 | 99.6 |
| | | 20,000 | 0.165589 | 0.529343 | 0.752904 | 0.016739 | 5.2 | 98.9 |
| | | 30,000 | 0.142673 | 0.76978 | 1.102976 | 0.024343 | 6.6 | 99 |
| | | 40,000 | 0.13608 | 0.499585 | 0.584051 | 0.015798 | 7.8 | 98.4 |
| | | 50,000 | 0.102723 | 0.460935 | 0.44649 | 0.014576 | 7.7 | 99.6 |
| | | 60,000 | 0.116631 | 0.34323 | 0.363245 | 0.010854 | 9.6 | 98.2 |
| | | 70,000 | 0.118904 | 0.246106 | 0.27617 | 0.007783 | 9.1 | 98.7 |
| | | 80,000 | 0.107707 | 0.25778 | 0.257697 | 0.008152 | 14 | 97.6 |
| | | 90,000 | 0.121068 | 0.291152 | 0.275773 | 0.009207 | 12.2 | 98 |
| | | 100,000 | 0.11602 | 0.213249 | 0.223246 | 0.006744 | 14.6 | 97.5 |
| | | 500,000 | 0.114313 | 0.081735 | 0.079503 | 0.002585 | 34.4 | 94.9 |
| | | 1,000,000 | 0.10923 | 0.055636 | 0.056011 | 0.001759 | 51.1 | 94.9 |
| α = 0.5 | θ = 0.111 | 10,000 | 0.195619 | 0.655741 | 1.049814 | 0.020736 | 5 | 99.2 |
| | | 20,000 | 0.171457 | 0.454953 | 0.587361 | 0.014387 | 5.9 | 98.7 |
| | | 30,000 | 0.127195 | 0.518937 | 0.704378 | 0.01641 | 8.2 | 99 |
| | | 40,000 | 0.138243 | 0.366566 | 0.418836 | 0.011592 | 9.5 | 98.4 |
| | | 50,000 | 0.107457 | 0.466333 | 0.629951 | 0.014747 | 9 | 99.3 |
| | | 60,000 | 0.11191 | 0.294334 | 0.32035 | 0.009308 | 11.1 | 98.3 |
| | | 70,000 | 0.121018 | 0.250276 | 0.271187 | 0.007914 | 10.9 | 98.6 |
| | | 80,000 | 0.111488 | 0.262529 | 0.281733 | 0.008302 | 15.3 | 97 |
| | | 90,000 | 0.125309 | 0.208127 | 0.223658 | 0.006582 | 14.9 | 98 |
| | | 100,000 | 0.121444 | 0.199154 | 0.197452 | 0.006298 | 16.7 | 97.2 |
| | | 500,000 | 0.113621 | 0.075993 | 0.074991 | 0.002403 | 38.7 | 95.2 |
| | | 1,000,000 | 0.109603 | 0.052219 | 0.05263 | 0.001651 | 56.7 | 95.4 |
| α = 1 | θ = 0.111 | 10,000 | 0.179866 | 0.479524 | 0.92671 | 0.015164 | 9 | 99.1 |
| | | 20,000 | 0.186004 | 0.383159 | 0.591125 | 0.012117 | 11.1 | 98.3 |
| | | 30,000 | 0.151255 | 0.418301 | 0.57858 | 0.013228 | 15.2 | 98.7 |
| | | 40,000 | 0.133393 | 0.331091 | 0.390104 | 0.01047 | 16.8 | 98.3 |
| | | 50,000 | 0.120147 | 0.308455 | 0.394251 | 0.009754 | 15.1 | 98.8 |
| | | 60,000 | 0.124971 | 0.269574 | 0.295066 | 0.008525 | 17.3 | 98.2 |
| | | 70,000 | 0.126138 | 0.323431 | 0.355051 | 0.010228 | 17.4 | 98.4 |
| | | 80,000 | 0.112269 | 0.225509 | 0.228095 | 0.007131 | 21.4 | 96.5 |

| | | | | | | | | |
|---|---|---|---|---|---|---|---|---|
| | | 90,000 | 0.125045 | 0.194023 | 0.217156 | 0.006136 | 22.1 | 97.1 |
| | | 100,000 | 0.117242 | 0.163908 | 0.164104 | 0.005183 | 24.9 | 97.1 |
| | | 500,000 | 0.111657 | 0.060065 | 0.059355 | 0.001899 | 52.5 | 96.4 |
| | | 1,000,000 | 0.110615 | 0.040422 | 0.041157 | 0.001278 | 74.7 | 95.8 |

#### 5.4 THETA=0.167

| Modelled mediator coefficient | Modelled interaction coefficient | Sample size | Mean 2sls interaction effect estimate | Standard deviation of 2sls estimate | Mean estimated standard error of 2sls estimate | Standard error of the bias (standard deviation of 2sls estimate/sqrt(1000)) | Power of 2sls estimator (%) | Coverage of 2sls estimator (%) |
|---|---|---|---|---|---|---|---|---|
| α = -0.333 | θ = 0.167 | 10,000 | 0.197331 | 1.013906 | 1.416338 | 0.032063 | 5.4 | 99.2 |
| | | 20,000 | 0.187588 | 0.649627 | 0.673347 | 0.020543 | 8.1 | 98.4 |
| | | 30,000 | 0.167889 | 0.464047 | 0.477021 | 0.014674 | 9.3 | 98.4 |
| | | 40,000 | 0.166374 | 0.327316 | 0.328466 | 0.010351 | 12.1 | 97.3 |
| | | 50,000 | 0.153254 | 0.302185 | 0.297196 | 0.009556 | 13.1 | 97.1 |
| | | 60,000 | 0.172735 | 0.237619 | 0.246105 | 0.007514 | 15.6 | 97.5 |
| | | 70,000 | 0.159085 | 0.225859 | 0.230669 | 0.007142 | 14 | 97.9 |
| | | 80,000 | 0.166032 | 0.218373 | 0.211477 | 0.006906 | 18.7 | 96.7 |
| | | 90,000 | 0.165702 | 0.196023 | 0.197819 | 0.006199 | 17.8 | 96.7 |
| | | 100,000 | 0.169058 | 0.188743 | 0.188055 | 0.005969 | 20.7 | 96.9 |
| | | 500,000 | 0.171353 | 0.083522 | 0.078537 | 0.002641 | 60.1 | 94.3 |
| | | 1,000,000 | 0.164694 | 0.05512 | 0.055633 | 0.001743 | 84.4 | 95.7 |
| α = 0 | θ = 0.167 | 10,000 | 0.254486 | 1.096615 | 1.650447 | 0.034678 | 7.7 | 98.8 |
| No mediation | | 20,000 | 0.241696 | 0.694182 | 0.884161 | 0.021952 | 9.7 | 98.3 |
| | | 30,000 | 0.193913 | 0.687636 | 0.787755 | 0.021745 | 11.9 | 98.2 |
| | | 40,000 | 0.175827 | 0.489114 | 0.502759 | 0.015467 | 14.4 | 97.3 |
| | | 50,000 | 0.151028 | 0.402886 | 0.382987 | 0.01274 | 14.7 | 98 |
| | | 60,000 | 0.170286 | 0.365595 | 0.343382 | 0.011561 | 16.5 | 97 |
| | | 70,000 | 0.165544 | 0.269808 | 0.27966 | 0.008532 | 14.8 | 98.1 |
| | | 80,000 | 0.166513 | 0.278154 | 0.274003 | 0.008796 | 19.4 | 95.9 |
| | | 90,000 | 0.171905 | 0.22407 | 0.228294 | 0.007086 | 21.1 | 97.7 |
| | | 100,000 | 0.169925 | 0.220342 | 0.213114 | 0.006968 | 21.8 | 97.1 |
| | | 500,000 | 0.171276 | 0.090857 | 0.085828 | 0.002873 | 53.7 | 94.2 |
| | | 1,000,000 | 0.164275 | 0.060318 | 0.060837 | 0.001907 | 77.3 | 95.4 |

| | | | | | | | | |
|---|---|---|---|---|---|---|---|---|
| α = 0.333 | θ = 0.167 | 10,000 | 0.281449 | 0.862732 | 1.423515 | 0.027282 | 9.7 | 98.5 |
| | | 20,000 | 0.24925 | 0.542047 | 0.767064 | 0.017141 | 13 | 97.7 |
| | | 30,000 | 0.218431 | 0.771141 | 1.106324 | 0.024386 | 16.8 | 97.8 |
| | | 40,000 | 0.207394 | 0.534682 | 0.617173 | 0.016908 | 18.8 | 97 |
| | | 50,000 | 0.168274 | 0.45721 | 0.454628 | 0.014458 | 19.3 | 98.1 |
| | | 60,000 | 0.180303 | 0.35498 | 0.376225 | 0.011225 | 19.8 | 96.8 |
| | | 70,000 | 0.177525 | 0.263748 | 0.30285 | 0.00834 | 18.4 | 97.8 |
| | | 80,000 | 0.166177 | 0.278174 | 0.275033 | 0.008797 | 23.7 | 95.4 |
| | | 90,000 | 0.178209 | 0.317524 | 0.292757 | 0.010041 | 24.6 | 96.8 |
| | | 100,000 | 0.175376 | 0.212486 | 0.218545 | 0.006719 | 27.1 | 95.7 |
| | | 500,000 | 0.170365 | 0.08551 | 0.083281 | 0.002704 | 56.3 | 94.7 |
| | | 1,000,000 | 0.164576 | 0.058432 | 0.058739 | 0.001848 | 79.2 | 95.3 |
| α = 0.5 | θ = 0.167 | 10,000 | 0.291685 | 0.661444 | 1.050285 | 0.020917 | 11.7 | 98.5 |
| | | 20,000 | 0.255066 | 0.476864 | 0.614482 | 0.01508 | 15.6 | 97.5 |
| | | 30,000 | 0.199539 | 0.537757 | 0.711098 | 0.017005 | 18.8 | 97.6 |
| | | 40,000 | 0.208956 | 0.387175 | 0.444912 | 0.012244 | 22.2 | 97 |
| | | 50,000 | 0.17062 | 0.540904 | 0.719625 | 0.017105 | 21.9 | 97.9 |
| | | 60,000 | 0.175187 | 0.316329 | 0.342877 | 0.010003 | 21.7 | 97.1 |
| | | 70,000 | 0.181411 | 0.260993 | 0.28036 | 0.008253 | 20.9 | 97.1 |
| | | 80,000 | 0.170233 | 0.282157 | 0.299607 | 0.008923 | 25.6 | 95 |
| | | 90,000 | 0.183009 | 0.216731 | 0.233047 | 0.006854 | 26.1 | 96.5 |
| | | 100,000 | 0.180029 | 0.209878 | 0.207586 | 0.006637 | 29.4 | 95.4 |
| | | 500,000 | 0.1696 | 0.080158 | 0.079073 | 0.002535 | 61.2 | 95.3 |
| | | 1,000,000 | 0.164985 | 0.055127 | 0.055536 | 0.001743 | 82.4 | 95.4 |
| α = 1 | θ = 0.167 | 10,000 | 0.271922 | 0.497693 | 0.95017 | 0.015738 | 18.8 | 97.7 |
| | | 20,000 | 0.2702 | 0.420568 | 0.645974 | 0.0133 | 23.7 | 96.4 |
| | | 30,000 | 0.22542 | 0.451278 | 0.623109 | 0.014271 | 27.6 | 96.7 |
| | | 40,000 | 0.201557 | 0.351069 | 0.416972 | 0.011102 | 29.9 | 96.5 |
| | | 50,000 | 0.185087 | 0.362523 | 0.455346 | 0.011464 | 29.2 | 96.7 |
| | | 60,000 | 0.188366 | 0.300819 | 0.319382 | 0.009513 | 31 | 96.7 |
| | | 70,000 | 0.189674 | 0.310527 | 0.337741 | 0.00982 | 33.4 | 97 |
| | | 80,000 | 0.170657 | 0.247531 | 0.252153 | 0.007828 | 35.5 | 94.8 |

| | | | | | | | | |
|---|---|---|---|---|---|---|---|---|
| | | 90,000 | 0.181635 | 0.21289 | 0.228958 | 0.006732 | 36.6 | 95.6 |
| | | 100,000 | 0.176135 | 0.178366 | 0.177016 | 0.00564 | 40.8 | 94.7 |
| | | 500,000 | 0.167407 | 0.06535 | 0.06394 | 0.002067 | 74.4 | 96.1 |
| | | 1,000,000 | 0.166108 | 0.043419 | 0.044296 | 0.001373 | 92.9 | 95.7 |

#### 5.5 THETA=0.333

| Modelled mediator coefficient | Modelled interaction coefficient | Sample size | Mean 2sls interaction effect estimate | Standard deviation of 2sls estimate | Mean estimated standard error of 2sls estimate | Standard error of the bias (standard deviation of 2sls estimate/sqrt(1000)) | Power of 2sls estimator (%) | Coverage of 2sls estimator (%) |
|---|---|---|---|---|---|---|---|---|
| α = -0.333 | θ = 0.333 | 10,000 | 0.428057 | 1.069232 | 1.458777 | 0.033812 | 16.6 | 97.1 |
| | | 20,000 | 0.369795 | 0.740297 | 0.749296 | 0.02341 | 22.4 | 95.5 |
| | | 30,000 | 0.334231 | 0.53054 | 0.53955 | 0.016777 | 27.1 | 96.8 |
| | | 40,000 | 0.338499 | 0.371777 | 0.373049 | 0.011757 | 31.8 | 95.6 |
| | | 50,000 | 0.319681 | 0.354974 | 0.342252 | 0.011225 | 33.1 | 95.3 |
| | | 60,000 | 0.34553 | 0.272033 | 0.278466 | 0.008602 | 37.8 | 96.1 |
| | | 70,000 | 0.322296 | 0.257233 | 0.261428 | 0.008134 | 37.8 | 97.1 |
| | | 80,000 | 0.335462 | 0.246562 | 0.238683 | 0.007797 | 42.4 | 95.2 |
| | | 90,000 | 0.329562 | 0.224228 | 0.223987 | 0.007091 | 44.2 | 96.9 |
| | | 100,000 | 0.332689 | 0.214283 | 0.212504 | 0.006776 | 46.3 | 96.5 |
| | | 500,000 | 0.338807 | 0.094062 | 0.088349 | 0.002975 | 92.9 | 94.2 |
| | | 1,000,000 | 0.330671 | 0.061866 | 0.062674 | 0.001956 | 99.7 | 95.9 |
| α = 0 | θ = 0.333 | 10,000 | 0.528586 | 1.23056 | 1.833369 | 0.038914 | 22 | 94.1 |
| No mediation | | 20,000 | 0.45919 | 0.796727 | 0.981171 | 0.025195 | 28 | 93.2 |
| | | 30,000 | 0.39031 | 0.79043 | 0.924352 | 0.024996 | 31.8 | 94.7 |
| | | 40,000 | 0.366844 | 0.58864 | 0.59584 | 0.018614 | 35.8 | 94 |
| | | 50,000 | 0.331268 | 0.495326 | 0.457067 | 0.015664 | 36.5 | 93.9 |
| | | 60,000 | 0.346623 | 0.45599 | 0.408503 | 0.01442 | 38.5 | 93.2 |
| | | 70,000 | 0.331844 | 0.347033 | 0.341126 | 0.010974 | 38.6 | 95.3 |
| | | 80,000 | 0.337116 | 0.312211 | 0.314922 | 0.009873 | 41.4 | 92.4 |
| | | 90,000 | 0.338328 | 0.261122 | 0.268376 | 0.008257 | 43.8 | 94.7 |
| | | 100,000 | 0.336367 | 0.259293 | 0.250686 | 0.0082 | 45.6 | 95.2 |
| | | 500,000 | 0.339141 | 0.105754 | 0.100298 | 0.003344 | 87.4 | 93.2 |
| | | 1,000,000 | 0.329946 | 0.070574 | 0.071263 | 0.002232 | 98.2 | 95.8 |

| | | | | | | | | |
|---|---|---|---|---|---|---|---|---|
| α = 0.333 | θ = 0.333 | 10,000 | 0.577363 | 1.022258 | 1.624879 | 0.032327 | 30.9 | 92.8 |
| | | 20,000 | 0.499734 | 0.630736 | 0.848212 | 0.019946 | 36.4 | 92.3 |
| | | 30,000 | 0.44525 | 0.820973 | 1.148082 | 0.025961 | 40 | 91.9 |
| | | 40,000 | 0.420911 | 0.660508 | 0.747598 | 0.020887 | 41 | 92.9 |
| | | 50,000 | 0.364533 | 0.478234 | 0.51281 | 0.015123 | 41.3 | 92.1 |
| | | 60,000 | 0.370939 | 0.41781 | 0.436842 | 0.013212 | 42.5 | 91.9 |
| | | 70,000 | 0.353037 | 0.389211 | 0.42966 | 0.012308 | 43 | 93.6 |
| | | 80,000 | 0.341237 | 0.357078 | 0.345618 | 0.011292 | 46.1 | 90.3 |
| | | 90,000 | 0.34929 | 0.419244 | 0.365197 | 0.013258 | 48.1 | 92.4 |
| | | 100,000 | 0.353088 | 0.304584 | 0.306412 | 0.009632 | 50.5 | 92.4 |
| | | 500,000 | 0.338183 | 0.10356 | 0.101379 | 0.003275 | 84.2 | 93.3 |
| | | 1,000,000 | 0.330284 | 0.071051 | 0.071642 | 0.002247 | 97.2 | 95.1 |
| α = 0.5 | θ = 0.333 | 10,000 | 0.579308 | 0.716971 | 1.088362 | 0.022673 | 34.8 | 91.7 |
| | | 20,000 | 0.505395 | 0.585861 | 0.735026 | 0.018527 | 41.6 | 92 |
| | | 30,000 | 0.41614 | 0.674192 | 0.859305 | 0.02132 | 42.1 | 91.1 |
| | | 40,000 | 0.420671 | 0.479873 | 0.554046 | 0.015175 | 44.5 | 91.5 |
| | | 50,000 | 0.359729 | 0.798667 | 1.031265 | 0.025256 | 44.8 | 91.7 |
| | | 60,000 | 0.36464 | 0.409033 | 0.433782 | 0.012935 | 45.3 | 91.3 |
| | | 70,000 | 0.362229 | 0.330984 | 0.343779 | 0.010467 | 45.7 | 92.8 |
| | | 80,000 | 0.346116 | 0.362177 | 0.377255 | 0.011453 | 47.8 | 90.1 |
| | | 90,000 | 0.355765 | 0.266053 | 0.285936 | 0.008413 | 51.3 | 91.4 |
| | | 100,000 | 0.355435 | 0.263591 | 0.258795 | 0.008335 | 53.6 | 91.1 |
| | | 500,000 | 0.337203 | 0.099789 | 0.098314 | 0.003156 | 86 | 93.5 |
| | | 1,000,000 | 0.330799 | 0.06827 | 0.069106 | 0.002159 | 97.5 | 94.9 |
| α = 1 | θ = 0.333 | 10,000 | 0.547539 | 0.591245 | 1.054379 | 0.018697 | 43.7 | 88.8 |
| | | 20,000 | 0.522284 | 0.587953 | 0.879207 | 0.018593 | 51.3 | 89.3 |
| | | 30,000 | 0.447471 | 0.577175 | 0.791724 | 0.018252 | 51.9 | 87.6 |
| | | 40,000 | 0.40564 | 0.432167 | 0.523054 | 0.013666 | 53.5 | 88.5 |
| | | 50,000 | 0.37952 | 0.555151 | 0.665846 | 0.017555 | 54.5 | 90.6 |
| | | 60,000 | 0.37817 | 0.429324 | 0.456015 | 0.013576 | 54.8 | 90.2 |
| | | 70,000 | 0.379901 | 0.369908 | 0.418037 | 0.011698 | 55.6 | 91.1 |
| | | 80,000 | 0.34547 | 0.344038 | 0.351551 | 0.010879 | 55.2 | 89.1 |

| | | | | | | | | |
|---|---|---|---|---|---|---|---|---|
| | | 90,000 | 0.351067 | 0.318713 | 0.339428 | 0.010079 | 59.4 | 90.5 |
| | | 100,000 | 0.352462 | 0.241217 | 0.235006 | 0.007628 | 60.8 | 89.7 |
| | | 500,000 | 0.334322 | 0.087996 | 0.084452 | 0.002783 | 90.8 | 93.7 |
| | | 1,000,000 | 0.332255 | 0.057 | 0.058392 | 0.001802 | 98.9 | 95.3 |

### 6 ALLOWING FOR PLEIOTROPIC EFFECTS OF THE INSTRUMENTS: Z=(Z1,Z2,Z1Z2,Z1Z1), N=500,000

#### 6.1 Standalone model – no pleiotropic effect

| Modelled mediator coefficient, α | Modelled interaction coefficient, θ | Mean 2sls interaction effect estimate | Standard deviation of 2sls estimate | Mean estimated standard error of 2sls estimate | Standard error of the bias (standard deviation of 2sls estimate/sqrt(1000)) | Power/ Type I error of 2sls estimator (%) | Coverage of 2sls estimator (%) |
|---|---|---|---|---|---|---|---|
| -0.333 | -0.111 | -0.105 | 0.064 | 0.077 | 0.002 | 29.8 | 95.7 |
| 0 | | -0.104 | 0.073 | 0.084 | 0.002 | 27.8 | 95.9 |
| 0.333 | | -0.105 | 0.071 | 0.081 | 0.002 | 31.8 | 96.2 |
| 0.5 | | -0.105 | 0.067 | 0.076 | 0.002 | 35.6 | 96.2 |
| 1 | | -0.108 | 0.052 | 0.060 | 0.002 | 50.7 | 96 |
| -0.333 | 0 | 0.005 | 0.065 | 0.075 | 0.002 | 4.2 [type I error] | 95.8 |
| 0 | | 0.006 | 0.074 | 0.081 | 0.002 | 4 [type I error] | 96 |
| 0.333 | | 0.005 | 0.071 | 0.077 | 0.002 | 3.8 [type I error] | 96.2 |
| 0.5 | | 0.004 | 0.065 | 0.073 | 0.002 | 3.8 [type I error] | 96.2 |
| 1 | | 0.002 | 0.048 | 0.056 | 0.002 | 3.2 [type I error] | 96.8 |
| -0.333 | 0.111 | 0.115 | 0.069 | 0.077 | 0.002 | 33.8 | 95.5 |
| 0 | | 0.116 | 0.080 | 0.083 | 0.003 | 32.4 | 95.6 |
| 0.333 | | 0.115 | 0.075 | 0.080 | 0.002 | 34.9 | 96.6 |
| 0.5 | | 0.114 | 0.068 | 0.076 | 0.002 | 39.2 | 96.6 |
| 1 | | 0.111 | 0.050 | 0.060 | 0.002 | 51.8 | 96.4 |
| -0.333 | 0.167 | 0.171 | 0.072 | 0.079 | 0.002 | 59.7 | 95.1 |
| 0 | | 0.171 | 0.084 | 0.086 | 0.003 | 54.3 | 96.3 |
| 0.333 | | 0.170 | 0.078 | 0.084 | 0.002 | 56.4 | 96.6 |
| 0.5 | | 0.169 | 0.071 | 0.080 | 0.002 | 60.3 | 96.1 |
| 1 | | 0.166 | 0.054 | 0.065 | 0.002 | 72.4 | 96.4 |
| -0.333 | 0.333 | 0.337 | 0.085 | 0.089 | 0.003 | 94.5 | 94.2 |
| 0 | | 0.336 | 0.101 | 0.101 | 0.003 | 85.7 | 95 |
| 0.333 | | 0.334 | 0.094 | 0.103 | 0.003 | 84.3 | 95.7 |
| 0.5 | | 0.333 | 0.086 | 0.100 | 0.003 | 85.3 | 95.7 |
| 1 | | 0.331 | 0.071 | 0.086 | 0.002 | 89.6 | 94.8 |

#### 6.2 Standalone model – pleiotropic effect

| Modelled mediator coefficient, α | Modelled interaction coefficient, θ | Mean 2sls interaction effect estimate | Standard deviation of 2sls estimate | Mean estimated standard error of 2sls estimate | Standard error of the bias (standard deviation of 2sls estimate/sqrt(1000)) | Power/ Type I error of 2sls estimator (%) | Coverage of 2sls estimator (%) |
|---|---|---|---|---|---|---|---|
| -0.333 | -0.111 | -0.099 | 0.142 | 0.176 | 0.004 | 10.7 | 97.9 |
| 0 | | -0.101 | 0.105 | 0.120 | 0.003 | 18.5 | 96.9 |
| 0.333 | | -0.104 | 0.076 | 0.087 | 0.002 | 29.2 | 96 |
| 0.5 | | -0.105 | 0.066 | 0.076 | 0.002 | 34.8 | 96.4 |
| 1 | | -0.106 | 0.048 | 0.055 | 0.002 | 54 | 96.2 |
| -0.333 | 0 | 0.010 | 0.143 | 0.172 | 0.005 | 2 [type I error] | 98 |
| 0 | | 0.009 | 0.108 | 0.116 | 0.003 | 3.2 [type I error] | 96.8 |
| 0.333 | | 0.006 | 0.077 | 0.083 | 0.002 | 3.4 [type I error] | 96.6 |
| 0.5 | | 0.005 | 0.067 | 0.072 | 0.002 | 3.5 [type I error] | 96.5 |
| 1 | | 0.003 | 0.047 | 0.051 | 0.001 | 3.6 [type I error] | 96.4 |
| -0.333 | 0.111 | 0.120 | 0.154 | 0.175 | 0.005 | 13.8 | 97.3 |
| 0 | | 0.118 | 0.116 | 0.119 | 0.004 | 20.1 | 96.6 |
| 0.333 | | 0.116 | 0.083 | 0.086 | 0.003 | 32.4 | 96.8 |
| 0.5 | | 0.115 | 0.072 | 0.075 | 0.002 | 39.3 | 96.7 |
| 1 | | 0.113 | 0.051 | 0.055 | 0.002 | 58.5 | 96.5 |
| -0.333 | 0.167 | 0.176 | 0.163 | 0.180 | 0.005 | 22.5 | 96.3 |
| 0 | | 0.173 | 0.123 | 0.123 | 0.004 | 36.4 | 96.7 |
| 0.333 | | 0.171 | 0.088 | 0.090 | 0.003 | 52.2 | 96.8 |
| 0.5 | | 0.170 | 0.076 | 0.080 | 0.002 | 59.9 | 96.5 |
| 1 | | 0.168 | 0.054 | 0.059 | 0.002 | 78.1 | 96.6 |
| -0.333 | 0.333 | 0.341 | 0.197 | 0.203 | 0.006 | 48.3 | 94.8 |
| 0 | | 0.338 | 0.147 | 0.145 | 0.005 | 67.8 | 94.7 |
| 0.333 | | 0.335 | 0.107 | 0.111 | 0.003 | 80.7 | 95.1 |
| 0.5 | | 0.334 | 0.093 | 0.099 | 0.003 | 84.9 | 95.1 |
| 1 | | 0.333 | 0.069 | 0.078 | 0.002 | 93.9 | 95.1 |

### 7 ALLOWING FOR PLEIOTROPIC EFFECTS OF THE INSTRUMENTS: FACTORIAL MR, N=500,000

#### 7.1 Standalone model – no pleiotropic effect

| Modelled mediator coefficient, α | Modelled interaction coefficient, θ | Power/Type I error of FMR to detect an interaction (%) |
|---|---|---|
| -0.333 | -0.111 | 17.7 |
| 0 | | 15.3 |
| 0.333 | | 13.2 |
| 0.5 | | 12.7 |
| 1 | | 10.2 |
| -0.333 | 0 | 5.5 [type I error] |
| 0 | | 5.5 [type I error] |
| 0.333 | | 5.5 [type I error] |
| 0.5 | | 5.5 [type I error] |
| 1 | | 5.5 [type I error] |
| -0.333 | 0.111 | 11.7 |
| 0 | | 11 |
| 0.333 | | 9.7 |
| 0.5 | | 9.1 |
| 1 | | 8.4 |
| -0.333 | 0.167 | 15.4 |
| 0 | | 13.7 |
| 0.333 | | 11.8 |
| 0.5 | | 10.8 |
| 1 | | 9.4 |
| -0.333 | 0.333 | 22.3 |
| 0 | | 17.8 |
| 0.333 | | 15.1 |
| 0.5 | | 13.8 |
| 1 | | 10.6 |
| Power and type I error defined using F statistic and Wald test | | |

#### 7.2 Standalone model – pleiotropic effect

| Modelled mediator coefficient, α | Modelled interaction coefficient, θ | Power/Type I error of FMR to detect an interaction (%) |
|---|---|---|
| -0.333 | -0.111 | 6.4 |
| 0 | | 10.1 |
| 0.333 | | 13.3 |
| 0.5 | | 15 |
| 1 | | 19.1 |
| -0.333 | 0 | 5.5 [type I error] |
| 0 | | 5.5 [type I error] |
| 0.333 | | 5.5 [type I error] |
| 0.5 | | 5.5 [type I error] |
| 1 | | 5.5 [type I error] |
| -0.333 | 0.111 | 6.2 |
| 0 | | 7.9 |
| 0.333 | | 9.5 |
| 0.5 | | 10.2 |
| 1 | | 12.5 |
| -0.333 | 0.167 | 7.3 |
| 0 | | 8.8 |
| 0.333 | | 11.2 |
| 0.5 | | 12.6 |
| 1 | | 15.3 |
| -0.333 | 0.333 | 9.1 |
| 0 | | 12.1 |
| 0.333 | | 14.4 |
| 0.5 | | 15.6 |
| 1 | | 18.2 |
| Power and type I error defined using F statistic and Wald test | | |

### 8 INCREASING THE VARIANCE EXPLAINED BY THE INSTRUMENTS, N=50,000, 2SLS INSTRUMENT ASSUMES MEDIATION

| Modelled mediator coefficient, α | Modelled interaction coefficient, θ | Mean 2sls interaction effect estimate | Standard deviation of 2sls estimate | Mean estimated standard error of 2sls estimate | Standard error of the bias (standard deviation of 2sls estimate/sqrt(1000)) | Power/ Type I error of 2sls estimator (%) | Coverage of 2sls estimator (%) |
|---|---|---|---|---|---|---|---|
| -0.333 | -0.111 | -0.109 | 0.050 | 0.049 | 0.002 | 61.9 | 94.9 |
| 0 | | -0.109 | 0.055 | 0.053 | 0.002 | 55.6 | 94.9 |
| 0.333 | | -0.109 | 0.052 | 0.051 | 0.002 | 58.8 | 95.8 |
| 0.5 | | -0.109 | 0.049 | 0.047 | 0.002 | 64 | 95.4 |
| 1 | | -0.109 | 0.038 | 0.036 | 0.001 | 81.2 | 95.1 |
| -0.333 | 0 | 0.001 | 0.048 | 0.048 | 0.002 | 4.4 | 95.6 |
| 0 | | 0.001 | 0.052 | 0.052 | 0.002 | 4.6 | 95.4 |
| 0.333 | | 0.001 | 0.049 | 0.048 | 0.002 | 4.8 | 95.2 |
| 0.5 | | 0.001 | 0.046 | 0.045 | 0.001 | 4.9 | 95.1 |
| 1 | | 0.001 | 0.034 | 0.034 | 0.001 | 4.8 | 95.2 |
| -0.333 | 0.111 | 0.112 | 0.049 | 0.049 | 0.002 | 63.6 | 95.4 |
| 0 | | 0.112 | 0.053 | 0.053 | 0.002 | 56 | 95.1 |
| 0.333 | | 0.112 | 0.050 | 0.050 | 0.002 | 61.4 | 95.9 |
| 0.5 | | 0.111 | 0.047 | 0.047 | 0.001 | 66.6 | 95.9 |
| 1 | | 0.111 | 0.036 | 0.036 | 0.001 | 83.2 | 95.6 |
| -0.333 | 0.167 | 0.167 | 0.050 | 0.050 | 0.002 | 89.5 | 95.3 |
| 0 | | 0.167 | 0.055 | 0.055 | 0.002 | 84.5 | 95.5 |
| 0.333 | | 0.167 | 0.053 | 0.053 | 0.002 | 84.3 | 95.5 |
| 0.5 | | 0.167 | 0.050 | 0.050 | 0.002 | 87.1 | 95.7 |
| 1 | | 0.166 | 0.039 | 0.039 | 0.001 | 95.2 | 96 |
| -0.333 | 0.333 | 0.333 | 0.057 | 0.057 | 0.002 | 99.8 | 95.6 |
| 0 | | 0.333 | 0.065 | 0.065 | 0.002 | 98.9 | 95.3 |
| 0.333 | | 0.332 | 0.065 | 0.065 | 0.002 | 98.6 | 95.6 |
| 0.5 | | 0.332 | 0.062 | 0.062 | 0.002 | 99 | 95.3 |
| 1 | | 0.332 | 0.052 | 0.052 | 0.002 | 99.8 | 94.6 |

### 9 HETEROSKEDASTIC ROBUST STANDARD ERRORS

#### 9.1 OLS REGRESSION, N=50,000

| Modelled mediator coefficient | Modelled interaction coefficient | Mean estimated standard error of 2sls estimate | Power/Type I Error of 2sls estimator (%) | Coverage of 2sls estimator (%) |
|---|---|---|---|---|
| -0.333 | -0.111 | 0.00305 | 100 | 0 |
| -0.333 | 0.000 | 0.00284 | 4.6 | 95.4 |
| -0.333 | 0.111 | 0.00305 | 100 | 0 |
| -0.333 | 0.167 | 0.00330 | 100 | 0 |
| -0.333 | 0.333 | 0.00441 | 100 | 0 |
| 0 | -0.111 | 0.00241 | 100 | 0 |
| 0 | 0.000 | 0.00229 | 5.1 | 94.9 |
| 0 | 0.111 | 0.00241 | 100 | 0 |
| 0 | 0.167 | 0.00256 | 100 | 0 |
| 0 | 0.333 | 0.00324 | 100 | 0 |
| 0.333 | -0.111 | 0.00187 | 100 | 0 |
| 0.333 | 0.000 | 0.00175 | 5.2 | 94.8 |
| 0.333 | 0.111 | 0.00187 | 100 | 0 |
| 0.333 | 0.167 | 0.00201 | 100 | 0 |
| 0.333 | 0.333 | 0.00262 | 100 | 0 |
| 0.5 | -0.111 | 0.00167 | 100 | 0 |
| 0.5 | 0.000 | 0.00155 | 5.1 | 94.9 |
| 0.5 | 0.111 | 0.00167 | 100 | 0 |
| 0.5 | 0.167 | 0.00181 | 100 | 0 |
| 0.5 | 0.333 | 0.00243 | 100 | 0 |
| 1 | -0.111 | 0.00126 | 100 | 0 |
| 1 | 0.000 | 0.00112 | 4.7 | 95.3 |
| 1 | 0.111 | 0.00126 | 100 | 0 |
| 1 | 0.167 | 0.00141 | 100 | 0 |
| 1 | 0.333 | 0.00205 | 100 | 0 |

#### 9.2 OLS REGRESSION, N=500,000

| Modelled mediator coefficient | Modelled interaction coefficient | Mean estimated standard error of 2sls estimate | Power/Type I Error of 2sls estimator (%) | Coverage of 2sls estimator (%) |
|---|---|---|---|---|
| -0.333 | -0.111 | 0.00097 | 100 | 0 |
| -0.333 | 0.000 | 0.00090 | 4.5 | 95.5 |
| -0.333 | 0.111 | 0.00097 | 100 | 0 |
| -0.333 | 0.167 | 0.00105 | 100 | 0 |
| -0.333 | 0.333 | 0.00140 | 100 | 0 |
| 0 | -0.111 | 0.00076 | 100 | 0 |
| 0 | 0.000 | 0.00072 | 3.6 | 96.4 |
| 0 | 0.111 | 0.00076 | 100 | 0 |
| 0 | 0.167 | 0.00081 | 100 | 0 |
| 0 | 0.333 | 0.00102 | 100 | 0 |
| 0.333 | -0.111 | 0.00059 | 100 | 0 |
| 0.333 | 0.000 | 0.00055 | 4.3 | 95.7 |
| 0.333 | 0.111 | 0.00059 | 100 | 0 |
| 0.333 | 0.167 | 0.00063 | 100 | 0 |
| 0.333 | 0.333 | 0.00083 | 100 | 0 |
| 0.5 | -0.111 | 0.00053 | 100 | 0 |
| 0.5 | 0.000 | 0.00049 | 4 | 96 |
| 0.5 | 0.111 | 0.00053 | 100 | 0 |
| 0.5 | 0.167 | 0.00057 | 100 | 0 |
| 0.5 | 0.333 | 0.00077 | 100 | 0 |
| 1 | -0.111 | 0.00040 | 100 | 0 |
| 1 | 0.000 | 0.00035 | 5.2 | 94.8 |
| 1 | 0.111 | 0.00040 | 100 | 0 |
| 1 | 0.167 | 0.00045 | 100 | 0 |
| 1 | 0.333 | 0.00065 | 100 | 0 |

#### 9.3 2SLS INSTRUMENT ASSUMES MEDIATION, Z=(Z1,Z2,Z1Z2,Z1Z1),N=50,000

| Modelled mediator coefficient | Modelled interaction coefficient | Mean estimated standard error of 2sls estimate | Power/Type I Error of 2sls estimator (%) | Coverage of 2sls estimator (%) |
|---|---|---|---|---|
| -0.333 | -0.111 | 0.283 | 8.1 | 98.1 |
| -0.333 | 0.000 | 0.278 | 1.8 | 98.2 |
| -0.333 | 0.111 | 0.289 | 7.3 | 97.8 |
| -0.333 | 0.167 | 0.299 | 12.9 | 97.6 |
| -0.333 | 0.333 | 0.344 | 33.5 | 95.5 |
| 0 | -0.111 | 0.342 | 8.7 | 97.7 |
| 0 | 0.000 | 0.345 | 0.7 | 99.3 |
| 0 | 0.111 | 0.368 | 7 | 99.5 |
| 0 | 0.167 | 0.385 | 14.7 | 98.1 |
| 0 | 0.333 | 0.460 | 36.8 | 94.1 |
| 0.333 | -0.111 | 0.472 | 9.4 | 97.6 |
| 0.333 | 0.000 | 0.450 | 0.2 | 99.8 |
| 0.333 | 0.111 | 0.451 | 7.4 | 99.6 |
| 0.333 | 0.167 | 0.459 | 19 | 98.4 |
| 0.333 | 0.333 | 0.517 | 41 | 92.1 |
| 0.5 | -0.111 | 0.383 | 10.3 | 97.3 |
| 0.5 | 0.000 | 0.490 | 0.2 | 99.8 |
| 0.5 | 0.111 | 0.635 | 8.3 | 99.3 |
| 0.5 | 0.167 | 0.726 | 21.3 | 98 |
| 0.5 | 0.333 | 1.041 | 44.2 | 91.9 |
| 1 | -0.111 | 0.328 | 18.2 | 97.1 |
| 1 | 0.000 | 0.298 | 0.2 | 99.8 |
| 1 | 0.111 | 0.397 | 15 | 98.8 |
| 1 | 0.167 | 0.459 | 28.9 | 96.8 |
| 1 | 0.333 | 0.671 | 54.6 | 90.8 |

#### 9.4 2SLS INSTRUMENT ASSUMES MEDIATION, Z=(Z1,Z2,Z1Z2,Z1Z1),N=500,000

| Modelled mediator coefficient | Modelled interaction coefficient | Mean estimated standard error of 2sls estimate | Power/Type I Error of 2sls estimator (%) | Coverage of 2sls estimator (%) |
|---|---|---|---|---|
| -0.333 | -0.111 | 0.0768 | 30.9 | 95 |
| -0.333 | 0.000 | 0.0751 | 5 | 95 |
| -0.333 | 0.111 | 0.0766 | 35.9 | 94.4 |
| -0.333 | 0.167 | 0.0786 | 60.1 | 94.3 |
| -0.333 | 0.333 | 0.0885 | 92.9 | 94.4 |
| 0 | -0.111 | 0.0831 | 29.9 | 94.5 |
| 0 | 0.000 | 0.0806 | 5.9 | 94.1 |
| 0 | 0.111 | 0.0830 | 32.3 | 94.4 |
| 0 | 0.167 | 0.0859 | 53.5 | 94.2 |
| 0 | 0.333 | 0.1005 | 87.3 | 93.2 |
| 0.333 | -0.111 | 0.0799 | 32.5 | 95.6 |
| 0.333 | 0.000 | 0.0765 | 5 | 95 |
| 0.333 | 0.111 | 0.0796 | 34.3 | 94.9 |
| 0.333 | 0.167 | 0.0834 | 56.3 | 94.6 |
| 0.333 | 0.333 | 0.1017 | 84.2 | 93.4 |
| 0.5 | -0.111 | 0.0753 | 35.9 | 95.8 |
| 0.5 | 0.000 | 0.0717 | 4.7 | 95.3 |
| 0.5 | 0.111 | 0.0751 | 38.8 | 95.5 |
| 0.5 | 0.167 | 0.0792 | 61.1 | 95.3 |
| 0.5 | 0.333 | 0.0987 | 86 | 93.5 |
| 1 | -0.111 | 0.0594 | 52.1 | 95.4 |
| 1 | 0.000 | 0.0555 | 3.9 | 96.1 |
| 1 | 0.111 | 0.0595 | 52.3 | 96.5 |
| 1 | 0.167 | 0.0641 | 74.2 | 96 |
| 1 | 0.333 | 0.0849 | 90.7 | 93.9 |

### 10 Examination of Sanderson-Windmeijer F Statistics

| Modelled mediator coefficient | F_statistic_2SLS_assuming_no_mediation | F_statistic_2SLS_assuming_mediation | N |
|---|---|---|---|
| -0.333 | 2.68 | 1.94 | 10000 |
| -0.333 | 4.88 | 3.20 | 20000 |
| -0.333 | 6.53 | 4.12 | 30000 |
| -0.333 | 9.10 | 5.50 | 40000 |
| -0.333 | 10.59 | 6.45 | 50000 |
| -0.333 | 12.64 | 7.66 | 60000 |
| -0.333 | 14.36 | 8.47 | 70000 |
| -0.333 | 16.56 | 9.76 | 80000 |
| -0.333 | 17.99 | 10.69 | 90000 |
| -0.333 | 19.76 | 11.55 | 1.00E+05 |
| -0.333 | 97.46 | 55.64 | 5.00E+05 |
| -0.333 | 191.17 | 109.02 | 1.00E+06 |
| 0 | 2.12 | 1.55 | 10000 |
| 0 | 3.54 | 2.28 | 20000 |
| 0 | 4.63 | 2.84 | 30000 |
| 0 | 6.29 | 3.64 | 40000 |
| 0 | 7.23 | 4.14 | 50000 |
| 0 | 8.60 | 4.81 | 60000 |
| 0 | 9.51 | 5.27 | 70000 |
| 0 | 11.18 | 6.09 | 80000 |
| 0 | 12.08 | 6.58 | 90000 |
| 0 | 13.28 | 7.13 | 1.00E+05 |
| 0 | 63.61 | 32.34 | 5.00E+05 |
| 0 | 123.87 | 62.49 | 1.00E+06 |
| 0.333 | 1.69 | 1.38 | 10000 |
| 0.333 | 2.51 | 1.87 | 20000 |
| 0.333 | 3.16 | 2.28 | 30000 |
| 0.333 | 4.12 | 2.78 | 40000 |
| 0.333 | 4.67 | 3.12 | 50000 |
| 0.333 | 5.48 | 3.50 | 60000 |
| 0.333 | 5.90 | 3.84 | 70000 |
| 0.333 | 7.02 | 4.39 | 80000 |
| 0.333 | 7.53 | 4.78 | 90000 |
| 0.333 | 8.27 | 5.20 | 1.00E+05 |

| | | | |
|---|---|---|---|
| 0.333 | 37.67 | 21.85 | 5.00E+05 |
| 0.333 | 72.64 | 41.85 | 1.00E+06 |
| 0.5 | 1.55 | 1.35 | 10000 |
| 0.5 | 2.18 | 1.78 | 20000 |
| 0.5 | 2.69 | 2.16 | 30000 |
| 0.5 | 3.43 | 2.59 | 40000 |
| 0.5 | 3.87 | 2.91 | 50000 |
| 0.5 | 4.49 | 3.22 | 60000 |
| 0.5 | 4.77 | 3.54 | 70000 |
| 0.5 | 5.70 | 4.02 | 80000 |
| 0.5 | 6.09 | 4.40 | 90000 |
| 0.5 | 6.68 | 4.80 | 1.00E+05 |
| 0.5 | 29.51 | 19.62 | 5.00E+05 |
| 0.5 | 56.55 | 37.54 | 1.00E+06 |
| 1 | 1.32 | 1.32 | 10000 |
| 1 | 1.63 | 1.71 | 20000 |
| 1 | 1.91 | 2.06 | 30000 |
| 1 | 2.29 | 2.40 | 40000 |
| 1 | 2.54 | 2.72 | 50000 |
| 1 | 2.85 | 2.94 | 60000 |
| 1 | 2.92 | 3.27 | 70000 |
| 1 | 3.50 | 3.64 | 80000 |
| 1 | 3.70 | 4.08 | 90000 |
| 1 | 4.03 | 4.47 | 1.00E+05 |
| 1 | 15.91 | 17.55 | 5.00E+05 |
| 1 | 29.83 | 33.69 | 1.00E+06 |

#### 10.1 F Statistic 2SLS using Z=Z1,Z2,Z1Z2 against sample size

A plot showing  the correlation of the Sanderson-Windmeijer F statistic(1) and sample size for the 2SLS estimator using Z=Z1,Z2,Z1Z2. Plot created using the *scatterplot* function from the *car* R package(2).

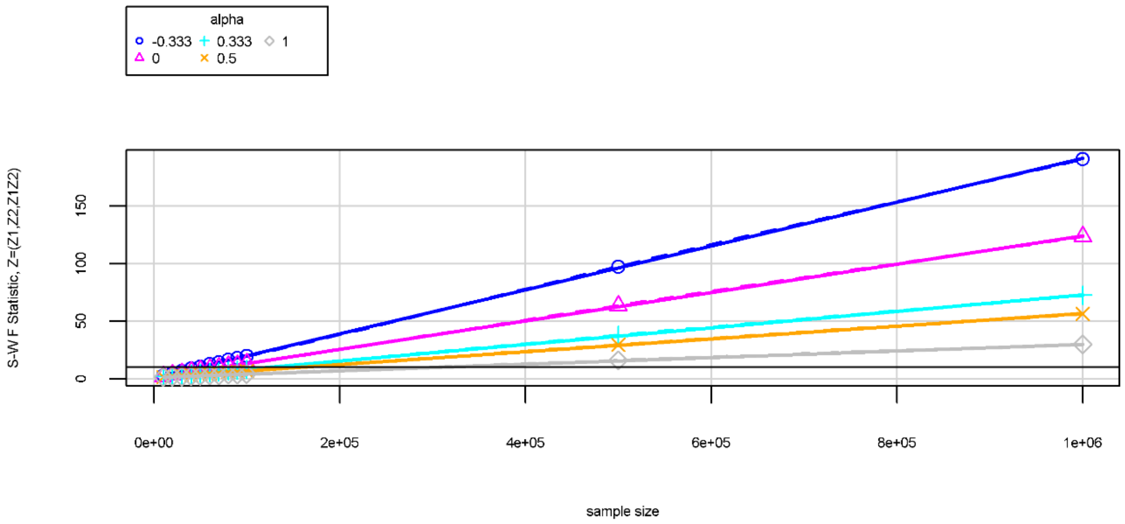

#### 10.2 F statistic 2SLS using Z=Z1,Z2,Z1Z2,Z1Z1 against sample size

A plot showing  the correlation of the Sanderson-Windmeijer F statistic(1) and sample size for the 2SLS estimator assuming  mediation. Plot created using the *scatterplot* function from the *car* R package(2)

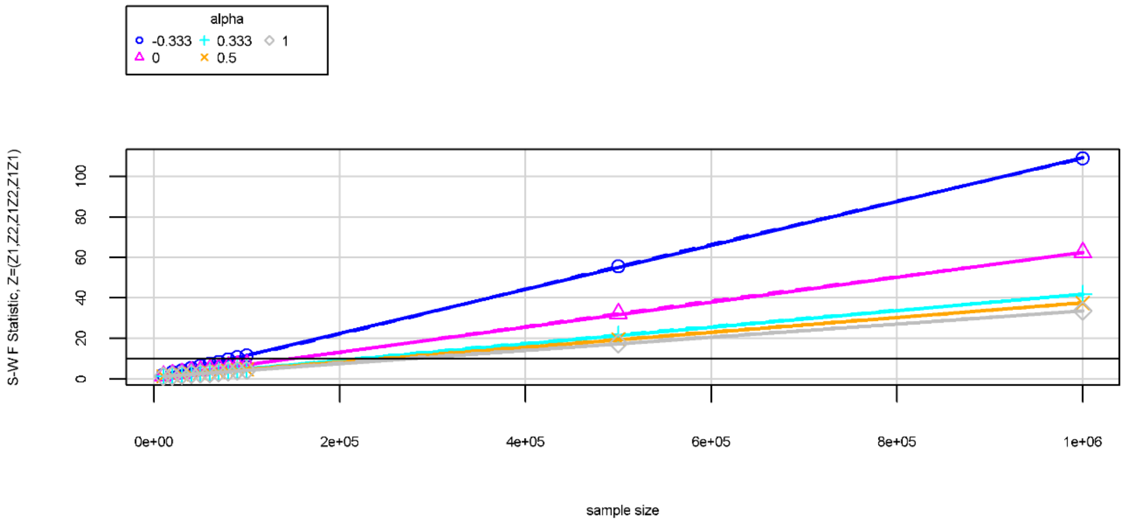

### 11 BIOBANK ILLUSTRATION

Models are described in detail in a subsequent section.

(*) adjusted for genotyping array

| Model | Estimated interactive effect (kg/m2 *years) | 95% Confidence interval | P-VALUE |
|---|---|---|---|
| 1 OLS | 0.029 | (0.026,0.031) | <0.0005 |
| 2 Instrument assumes mediation Z=(Z1,Z2,Z1Z2,Z1Z1) | -0.037 | (-0.357, 0.283) | 0.820 |
| 3 Instrument assumes mediation Z=(Z1,Z2,Z1Z2,Z1Z1)* | -0.037 | (-0.357, 0.283) | 0.820 |
| 4 Instrument does not assume mediation Z=(Z1,Z2,Z1Z2) | -0.050 | (-0.371,0.271) | 0.761 |
| 5 Instrument does not assume mediation Z=(Z1,Z2,Z1Z2)* | -0.050 | (-0.371,0.271) | 0.761 |
| 8 FMR | NA | NA | 0.250 |
| 9 FMR* | NA | NA | 0.251 |

### 12 SUPPLEMENTAL METHODS

#### 12.1 Inclusion of $Z_1Z_1$ term in the instrument

If we substitute the A1 equation into the A2 equation and multiply we get

$$
\begin{aligned}
A_1A_2 = {} & C^2 + CE + 0.18CZ_2 + \alpha C^2 + \alpha CV + 0.14C\alpha Z_1 + \\
& VC + VE + 0.18VZ_2 + \alpha VC + \alpha V^2 + 0.14V\alpha Z_1 + \\
& 0.14CZ_1 + 0.14EZ_1 + 0.14*0.18\mathbf{Z_1Z_2} + 0.14\alpha CZ_1 + \\
& 0.14\alpha VZ_1 + 0.14*0.14\alpha\mathbf{Z_1Z_1}
\end{aligned}
$$

We see that when mediation is assumed ($\alpha$ is assumed to be non-zero), there are $Z_1Z_2$ and $Z_1Z_1$ terms. When mediation is not assumed, the $Z_1Z_1$ term vanishes.

#### 12.2 Biobank example

The models tested are listed below. Principal components refer to principal components of population stratification.

1. Linear regression of systolic blood pressure on BMI, years of schooling, BMI*years of schooling, age and sex
2. Instrumental variable regression with
    a. Outcome: systolic blood pressure
    b. Instrumented variables: BMI, years of schooling, BMI*years of schooling
    c. Included instruments: age, sex, 40 principal components
    d. Excluded instruments: years of schooling PRS, BMI PRS, years of schooling PRS*BMI PRS, years of schooling PRS* years of schooling PRS
3. Instrumental variable regression with
    a. Outcome: systolic blood pressure
    b. Instrumented variables: BMI, years of schooling, BMI*years of schooling
    c. Included instruments: age, sex, 40 principal components, genotyping array
    d. Excluded instruments: years of schooling PRS, BMI PRS, years of schooling PRS*BMI PRS, years of schooling PRS* years of schooling PRS
4. Instrumental variable regression with
    a. Outcome: systolic blood pressure
    b. Instrumented variables: BMI, years of schooling, BMI*years of schooling
    c. Included instruments: age, sex, 40 principal components
    d. Excluded instruments: years of schooling PRS, BMI PRS, years of schooling PRS*BMI PRS
5. Instrumental variable regression with
    a. Outcome: systolic blood pressure
    b. Instrumented variables: BMI, years of schooling, BMI*years of schooling
    c. Included instruments: age, sex, 40 principal components, genotyping array
    d. Excluded instruments: years of schooling PRS, BMI PRS, years of schooling PRS*BMI PRS
6. Instrumental variable regression with
    a. Outcome: systolic blood pressure
    b. Instrumented variables: BMI, years of schooling
    c. Included instruments: age, sex, 40 principal components
    d. Excluded instruments: years of schooling PRS, BMI PRS
7. Instrumental variable regression with
    a. Outcome: systolic blood pressure
    b. Instrumented variables: BMI, years of schooling
    c. Included instruments: age, sex, 40 principal components, genotyping array
    d. Excluded instruments: years of schooling PRS, BMI PRS
8. Linear regression of systolic blood pressure on the three non-reference categories of FMR (low-high, high-low and high-high PRS scores), adjusted for age, sex, 40 principal components, followed by test of linear combination of FMR category coefficients
9. Linear regression of systolic blood pressure on the three non-reference categories of FMR (low-high, high-low and high-high PRS scores), adjusted for age, sex, 40 principal

components and genotyping array, followed by test of linear combination of FMR category coefficients

#### 12.3 UK Biobank - Further Information and Quality Control

UK Biobank is a population-based health research resource consisting of approximately 500,000 people, aged between 38 years and 73 years, who were recruited between the years 2006 and 2010 from across the UK(3). Particularly focused on identifying determinants of human diseases in middle-aged and older individuals, participants provided a range of information (such as demographics, health status, lifestyle measures, cognitive testing, personality self-report, and physical and mental health measures) via questionnaires and interviews; anthropometric measures, BP readings and samples of blood, urine and saliva were also taken (data available at www.ukbiobank.ac.uk). A full description of the study design, participants and quality control (QC) methods have been described in detail previously(4). UK Biobank received ethical approval from the Research Ethics Committee (REC reference for UK Biobank is 11/NW/0382).

The full data release contains the cohort of successfully genotyped samples (n=488,377). 49,979 individuals were genotyped using the UK BiLEVE array and 438,398 using the UK Biobank axiom array. Pre-imputation QC, phasing and imputation are described elsewhere(5). In brief, prior to phasing, multiallelic SNPs or those with MAF ≤1% were removed. Phasing of genotype data was performed using a modified version of the SHAPEIT2 algorithm(6). Genotype imputation to a reference set combining the UK10K haplotype and HRC reference panels(7) was performed using IMPUTE2 algorithms(8). The analyses presented here were restricted to autosomal variants within the HRC site list using a graded filtering with varying imputation quality for different allele frequency ranges. Therefore, rarer genetic variants are required to have a higher imputation INFO score (Info>0.3 for MAF >3%; Info>0.6 for MAF 1-3%; Info>0.8 for MAF 0.5-1%; Info>0.9 for MAF 0.1-0.5%) with MAF and Info scores having been recalculated on an in house derived ‘European’ subset.

Individuals with sex-mismatch (derived by comparing genetic sex and reported sex) or individuals with sex-chromosome aneuploidy were excluded from the analysis (n=814).
We restricted the sample to individuals of white British ancestry who self-report as "White British" and who have very similar ancestral backgrounds according to the PCA (n=409,703), as described by Bycroft(5).

Estimated kinship coefficients using the KING toolset(9) identified 107,162 pairs of individuals(5). An in-house algorithm was then applied to this list and preferentially removed the individuals related to the greatest number of other individuals until no related pairs remain. These individuals were excluded (n=79,448). Additionally 2 individuals were removed due to them relating to a very large number (>200) of individuals.

Quality Control filtering of the UK Biobank data was conducted by R.Mitchell, G.Hemani, T.Dudding, L.Paternoster as described in the published protocol (doi: 10.5523/bris.3074krb6t2frj29yh2b03x3wxj)(10).

### 13 SUPPLEMENTAL FIGURES

#### 13.1 Ordinary Least Squares: bias at N=50,000

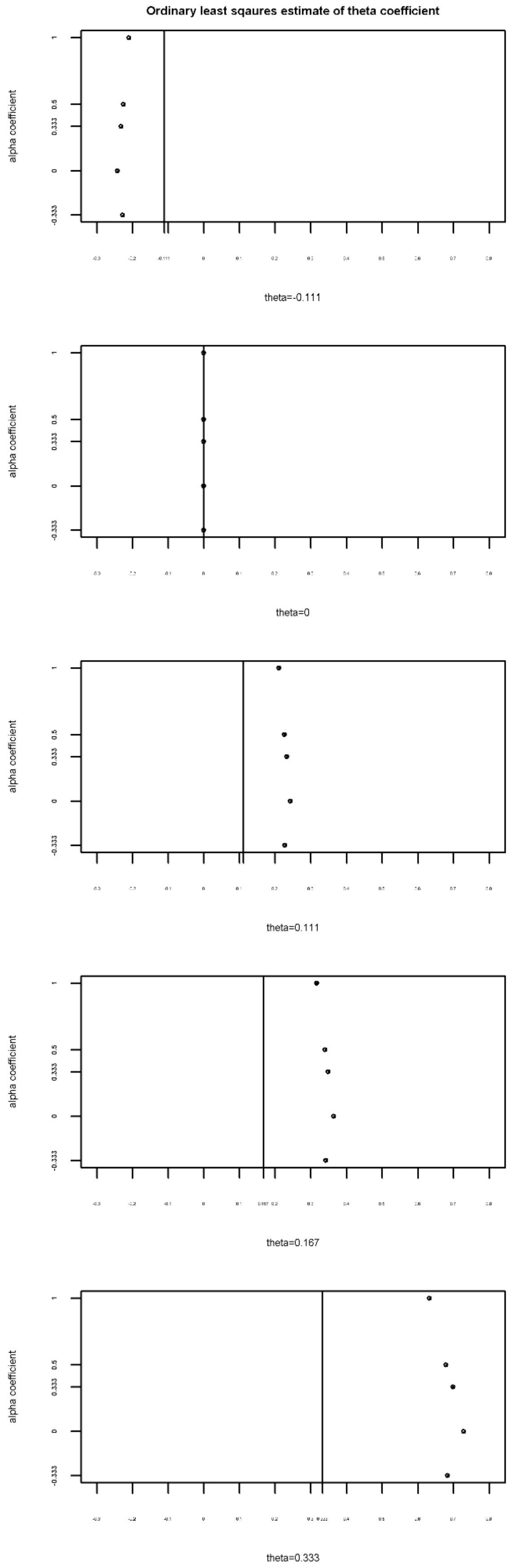

#### 13.2 Ordinary Least Squares bias: at N=500,00

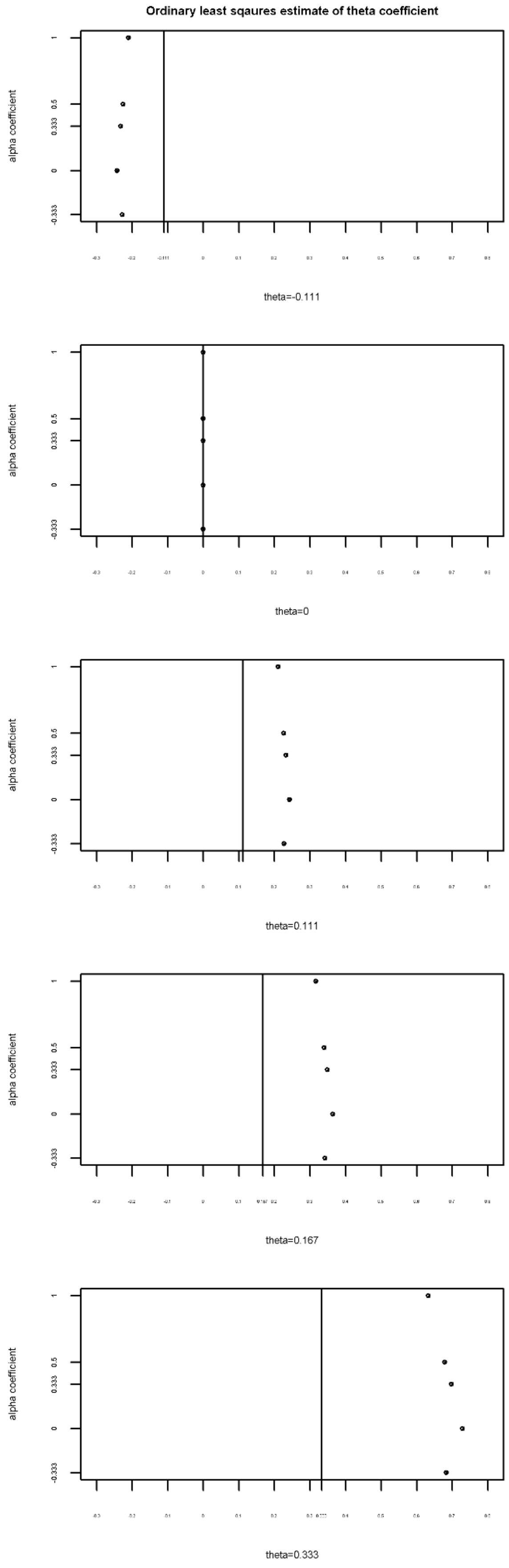

#### 13.3 2SLS using Z=Z1,Z2,Z1Z2: bias at N=50,000, Z=(Z1,Z2,Z1Z2)

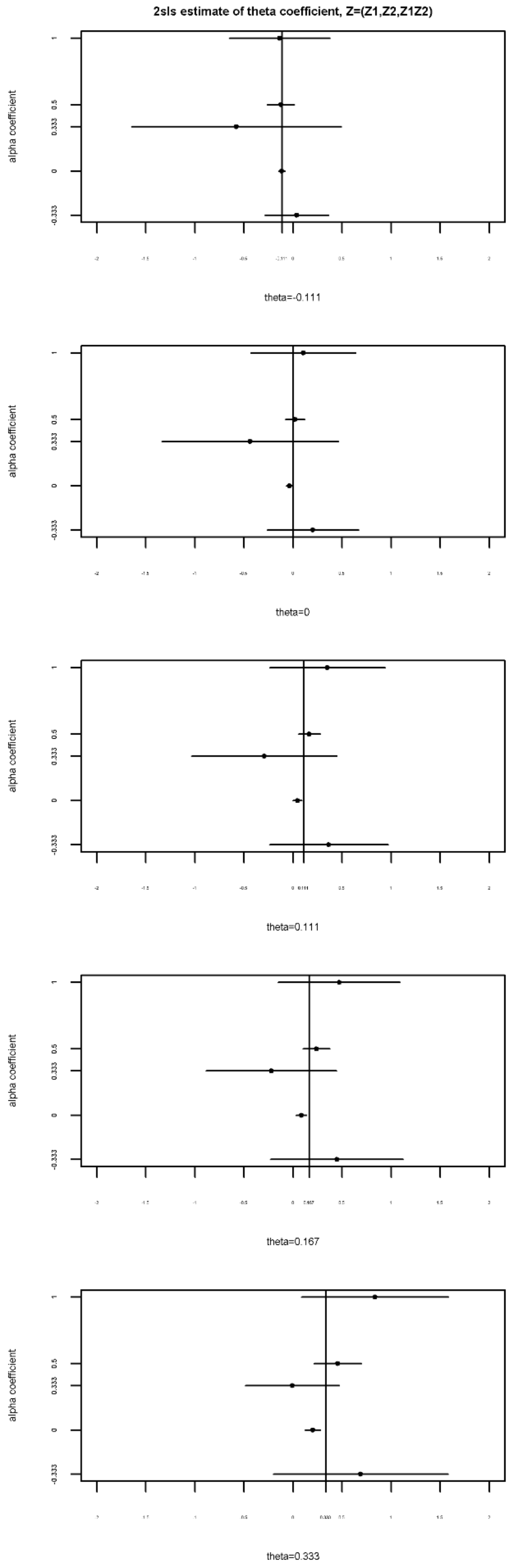

#### 13.4 2SLS using Z=Z1,Z2,Z1Z2: bias at N=100,000, Z=(Z1,Z2,Z1Z2)

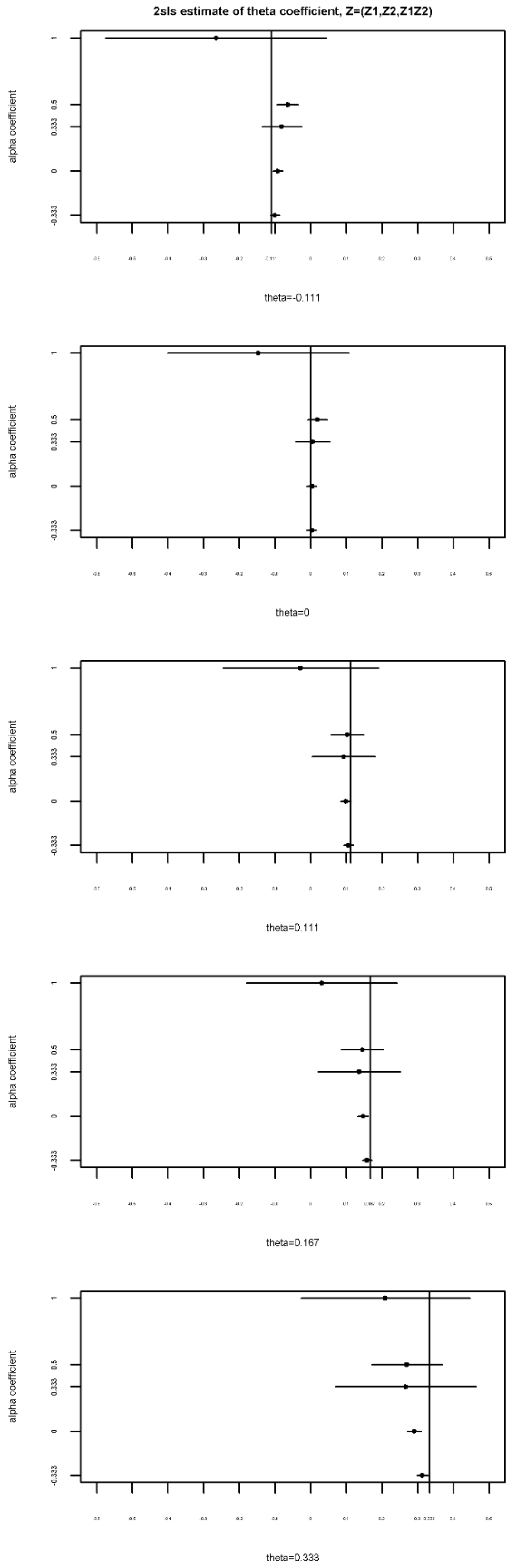

#### 13.5 2SLS using Z=Z1,Z2,Z1Z2: bias at N=500,000, Z=(Z1,Z2,Z1Z2)

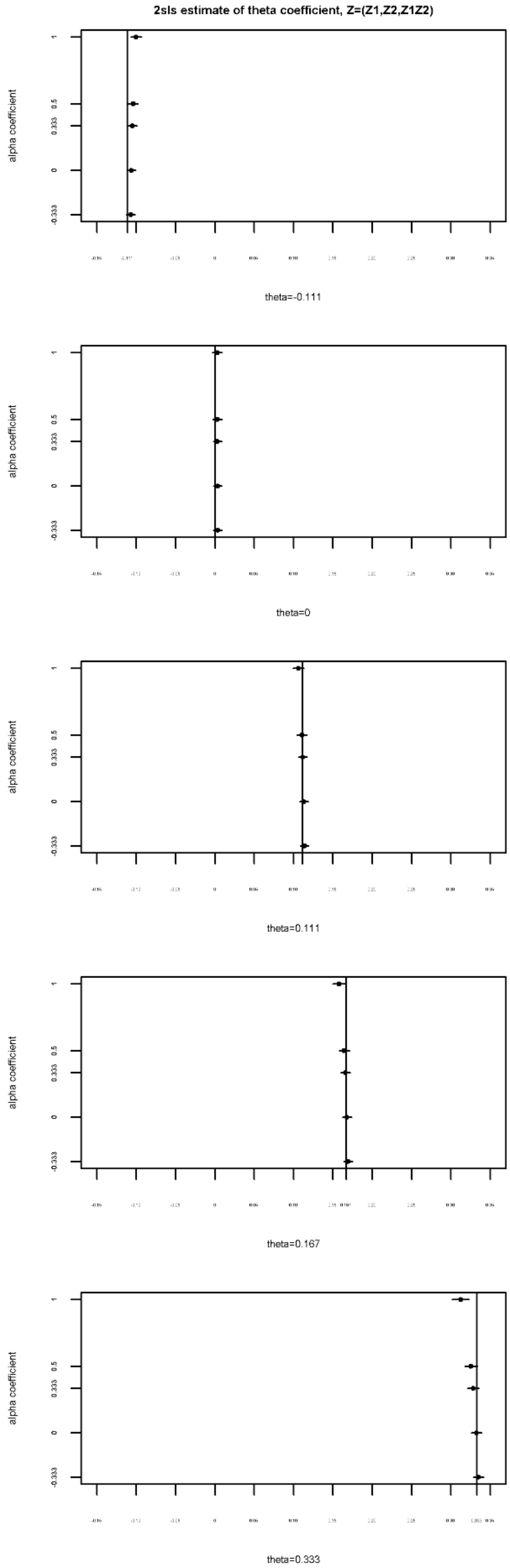

#### 13.6 2SLS using Z=Z1,Z2,Z1Z2,Z1Z1 (Z=(Z1,Z2,Z1Z2,Z1Z1),power and coverage) versus FMR (power) when N=100,000

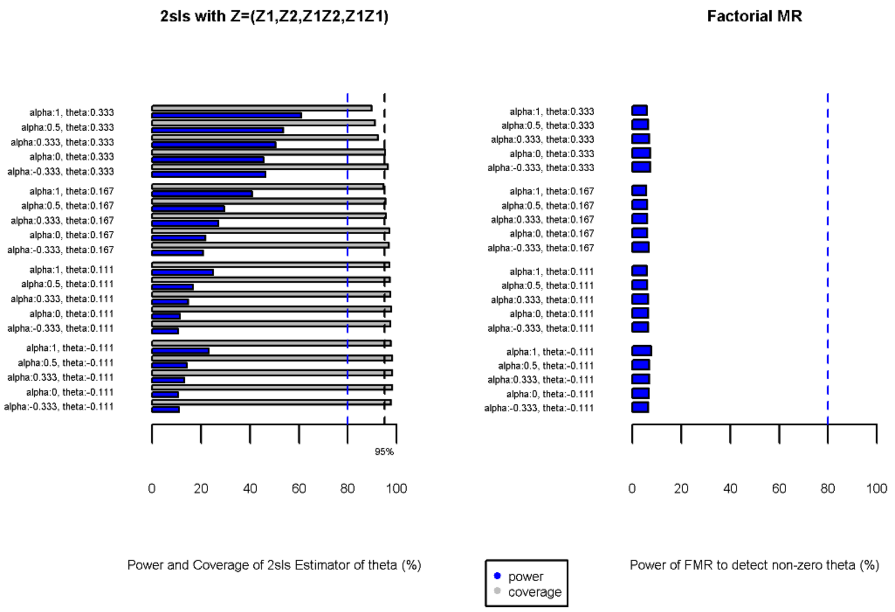

#### 13.7 Type I error rate of 2SLS using Z=Z1,Z2,Z1Z2,Z1Z1 (Z=Z1,Z2,Z1Z2,Z1Z1) versus FMR at N=50,000

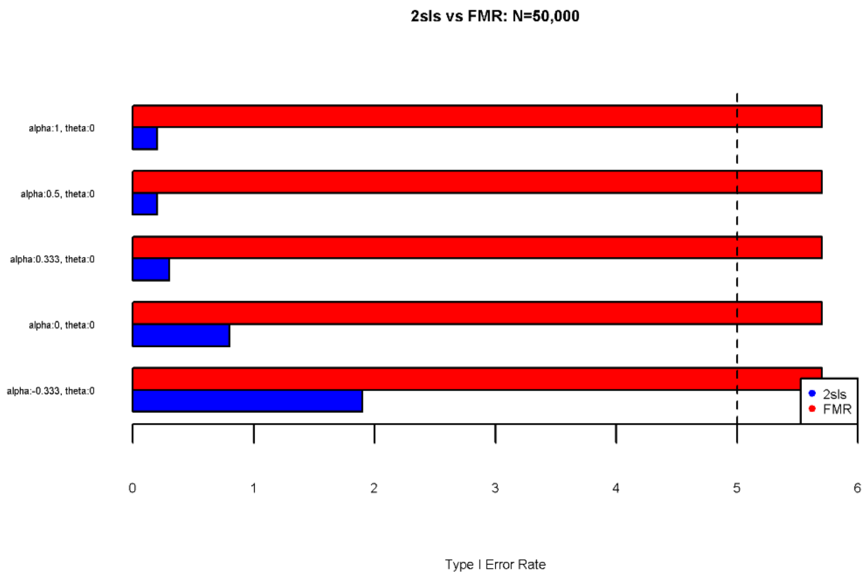

#### 13.8 Type I error rate of 2SLS using Z=Z1,Z2,Z1Z2,Z1Z1 (Z=Z1,Z2,Z1Z2,Z1Z1) versus FMR at N=100,000

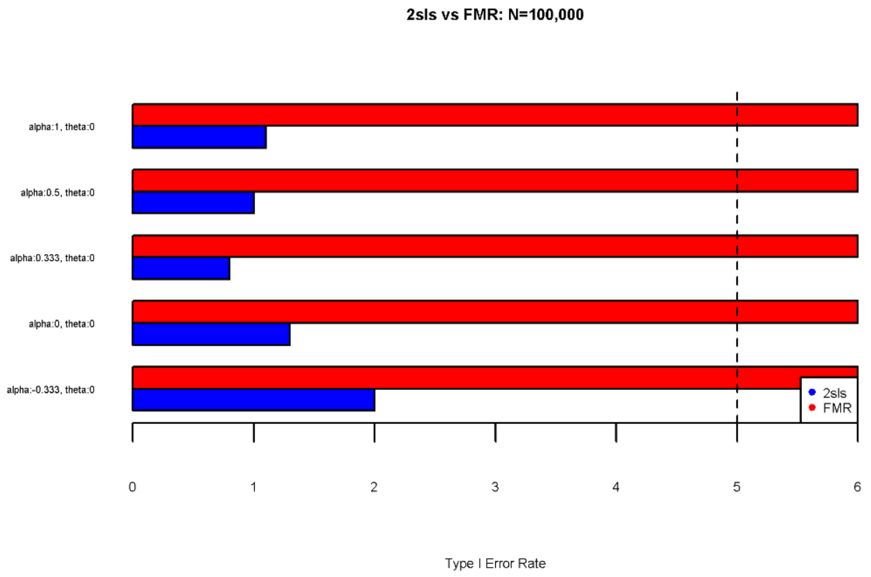

#### 13.9 Type I error rate of 2SLS using Z=Z1,Z2,Z1Z2,Z1Z1 (Z=Z1,Z2,Z1Z2,Z1Z1) versus FMR at N=500,000

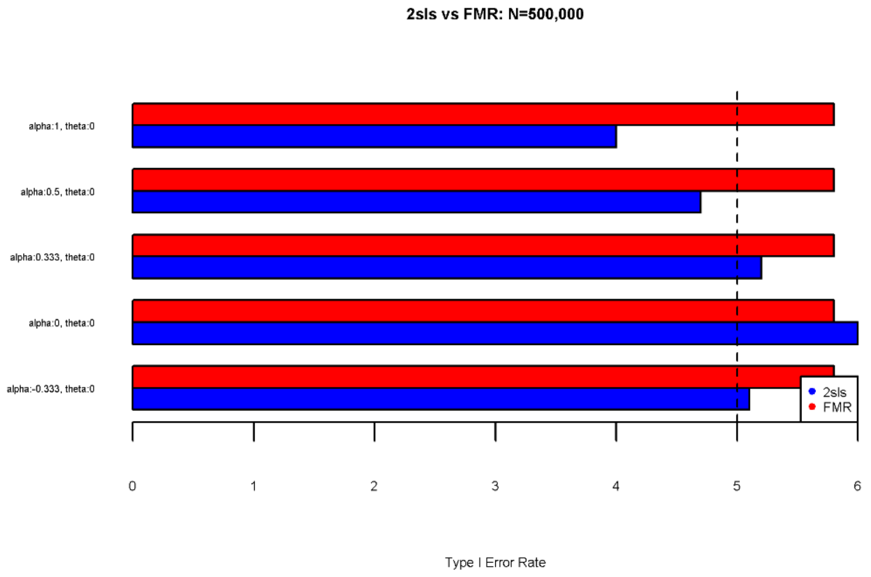

### 14 Supplemental References

1. Sanderson E, Windmeijer F. A weak instrument F-test in linear IV models with multiple endogenous variables. Journal of Econometrics. 2016;190(2):212-21.
2. Fox J, Weisberg S. An {R} Companion to Applied Regression, Second Edition. Thousand Oaks, CA: Sage; 2011.
3. Allen NE, Sudlow C, Peakman T, Collins R. UK Biobank Data: Come and Get It. Science Translational Medicine. 2014;6(224):224ed4.
4. Collins R. What makes UK Biobank special? The Lancet. 2012;379(9822):1173-4.
5. Bycroft C, Freeman C, Petkova D, Band G, Elliott LT, Sharp K, et al. Genome-wide genetic data on ~500,000 UK Biobank participants. bioRxiv. 2017.
6. O'Connell J, Sharp K, Shrine N, Wain L, Hall I, Tobin M, et al. Haplotype estimation for biobank-scale data sets. Nature Genetics. 2016;48:817.
7. Huang J, Howie B, McCarthy S, Memari Y, Walter K, Min JL, et al. Improved imputation of low-frequency and rare variants using the UK10K haplotype reference panel. Nature Communications. 2015;6:8111.
8. Howie B, Marchini J, Stephens M. Genotype Imputation with Thousands of Genomes. G3: Genes|Genomes|Genetics. 2011;1(6):457.
9. Manichaikul A, Mychaleckyj JC, Rich SS, Daly K, Sale M, Chen W-M. Robust relationship inference in genome-wide association studies. Bioinformatics. 2010;26(22):2867-73.
10. Mitchell R, Hemani G, Dudding T, Paternoster L. UK Biobank Genetic Data: MRC-IEU Quality Control, Version 1. 2017.
